# X chromosomes show relaxed selection and complete somatic dosage compensation across *Timema* stick insect species

**DOI:** 10.1101/2021.11.28.470265

**Authors:** Darren J. Parker, Kamil S. Jaron, Zoé Dumas, Marc Robinson-Rechavi, Tanja Schwander

**Affiliations:** Department of Ecology and Evolution, University of Lausanne, Lausanne, Switzerland; Swiss Institute of Bioinformatics, Lausanne, Switzerland; School of Natural Sciences, Bangor University, United Kingdom; Institute of Evolutionary Biology, School of Biological Sciences, University of Edinburgh, Edinburgh, EH9 3FL

## Abstract

Sex chromosomes have evolved repeatedly across the tree of life. As they are present in different copy numbers in males and females, they are expected to experience different selection pressures than the autosomes, with consequences including a faster rate of evolution, increased accumulation of sexually antagonistic alleles, and the evolution of dosage compensation. Whether these consequences are general or linked to idiosyncrasies of specific taxa is not clear as relatively few taxa have been studied thus far. Here we use wholegenome sequencing to identify and characterize the evolution of the X chromosome in five species of *Timema* stick insects with XX:X0 sex determination. The X chromosome had a similar size (approximately 11% of the genome) and gene content across all five species, suggesting that the X chromosome originated prior to the diversification of the genus. Genes on the X showed evidence of relaxed selection (elevated dN/dS) and a slower evolutionary rate (dN + dS) than genes on the autosomes, likely due to sex-biased mutation rates. Genes on the X also showed almost complete dosage compensation in somatic tissues (heads and legs), but dosage compensation was absent in the reproductive tracts. Contrary to prediction, sex-biased genes showed little enrichment on the X, suggesting that the advantage X-linkage provides to the accumulation of sexually antagonistic alleles is weak. Overall, we found the consequences of X-linkage on gene sequences and expression to be similar across *Timema* species, showing the characteristics of the X chromosome are surprisingly consistent over 30 million years of evolution.

## Introduction

One of the most common forms of genomic variation between individuals within species stems from sex chromosomes. Sex chromosomes differ in copy number between males and females, which has a large effect on the evolutionary forces acting on genes located on them. Specifically, when males are the heterogametic sex (i.e., in XY or X0 systems), the genes on the X chromosome are present in only a single copy in males, while genes on the autosomes are present in two copies. This fundamental difference is expected to have a large effect on the evolutionary forces acting on genes located on the X chromosome, with consequences including a faster rate of sequence evolution, increased accumulation of sexually antagonistic alleles, and the evolution of dosage compensation mechanisms (Bachtrog *et al*., 2011; Wright *et al*., 2016; Lenormand & Roze, 2021). Such effects should apply across taxa, meaning we should observe common patterns of X chromosome evolution in diverse species. Despite this, studies of sex chromosome evolution have shown a great deal of variation among study systems, including differences in the influence selection and drift have on the content of the X and in the extent of dosage compensation (Bachtrog *et al*., 2011; Gu & Walters, 2017). Currently, it is difficult to understand the factors that govern this variation as only few taxa (typically with only a single or few representative species) have been studied. To elucidate these factors, studies of X chromosome evolution are thus needed from multiple species from a wide range of taxa (Palmer *et al*., 2019). Such studies will allow us to disentangling general sex chromosome-linked patterns from species-specific patterns, and allow us to develop a fuller understanding of sex chromosome evolution.

The rate of sequence evolution is expected to differ between the X chromosome and autosomes for two main reasons. Firstly, the X is hemizygous in males, meaning that recessive or partially recessive mutations in X-linked genes will be more exposed to selection than mutations in genes on the autosomes, allowing for beneficial mutations to be fixed and deleterious mutations to be purged more effectively (Charlesworth *et al*., 1987). On the other hand, as the X is only present in a single copy in males, its effective population size is expected to be smaller than that of the autosomes. This means that the effects of drift will also be stronger for genes on the X chromosome than on the autosomes, which will tend to reduce the fixation rate for advantageous mutations but increase it for deleterious mutations (Wright, 1931; Vicoso & Charlesworth, 2009). Together, these effects have been used as an explanation for the overall faster evolution of the X chromosome (the faster-X effect) seen in many species (Mank *et al*., 2010; Meisel & Connallon, 2013; Parsch & Ellegren, 2013; Charlesworth *et al*., 2018). In addition, the difference in X chromosome copy number between males and females is expected to facilitate the fixation of sexually antagonistic alleles (Rice, 1984; Gibson *et al*., 2002; Mullon *et al*., 2012). This is because the X chromosome spends two-thirds of its time in females, giving an advantage to dominant female-beneficial alleles on the X. In addition, recessive alleles on the X will be exposed to selection on the X in males, giving an advantage to male-beneficial alleles. These complex forces have the potential to shape how sexually antagonistic variation is distributed across the genome, which can in turn influence broad evolutionary processes such as speciation (Coyne & Orr, 2004; Payseur *et al*., 2018) and sexual conflict (Bachtrog *et al*., 2011; Mank *et al*., 2014; Wilkinson *et al*., 2015).

The fact that genes on the X are present in different copy numbers in males and females can also create a problem for gene expression, as for many genes expression is proportional to their copy number (Birchler & Veitia, 2012; Birchler, 2016). As such, species with differentiated sex chromosomes should have evolved dosage compensation mechanisms to equalise expression of the X chromosome in males and females (Ohno, 1967; Charlesworth, 1978, 1996). Note that such dosage compensation mechanisms can evolve either in response to sex chromosomes differentiation or alongside it (Lenormand *et al*., 2020). Dosage compensation has been demonstrated across a wide range of taxa (Disteche, 2012; Mank, 2013; Gu & Walters, 2017), including model species such as *Drosophila melanogaster* (Conrad & Akhtar, 2012) and *Caenorhabditis elegans* (Meyer, 2000) where this phenomenon has been studied in detail (Parkhurst & Meneely, 1994; Lucchesi, 1998; Meyer, 2000; Straub & Becker, 2011; Conrad & Akhtar, 2012). Despite this commonality, it has become increasingly clear that the extent to which genes on the X are dosage compensated varies among species. Several studied species show only partial or no dosage compensation (Mank, 2013). The extent of dosage compensation may also differ by tissue type, with reduced dosage compensation observed in the reproductive tracts of e.g. *C. elegans* (Kelly *et al*., 2002; Pirrotta, 2002) or *D. melanogaster* (Oliver, 2002; Meiklejohn *et al*., 2011; Mahadevaraju *et al*., 2021). However, it is not clear how widespread tissue-specific dosage compensation is, as work in non-model species often use whole-body samples for examining expression (Gu & Walters, 2017).

Here we expand our knowledge of the evolutionary characteristics of sex chromosomes by identifying and studying the X chromosome in *Timema* stick insects. Aspects of X chromosome evolution have been previously studied in several insect orders (Odonata (Chauhan *et al*., 2021), Hemiptera (Pal & Vicoso, 2015; Richard *et al*., 2017), Orthoptera (Rayner *et al*., 2021), Strepsiptera (Mahajan & Bachtrog, 2015), Coleoptera (Prince *et al*., 2010; Mahajan & Bachtrog, 2015), and Diptera (Bone & Kuroda, 1996; Marín *et al*., 1996; Deng *et al*., 2011; Nozawa *et al*., 2014; Jiang *et al*., 2015; Vicoso & Bachtrog, 2015; Rose *et al*., 2016)). However, to date no studies have examined X chromosome evolution in stick insects (Phasmatodea), an order which originated approximately 130 mya (Simon *et al*., 2019) and contains around 3100 extant species (Bradler & Buckley, 2018). First, we identified the X chromosome in five species of *Timema* that diverged approximately 30 mya (Riesch *et al*., 2017). *Timema* have an XX/X0 sex determination system (Schwander & Crespi, 2009). To determine if genes on the X chromosome show the predicted faster rate of sequence evolution than genes on the autosomes, we examined sequence evolution rates in each species. In addition, we tested if the X chromosome is enriched for sexually antagonistic alleles. This was done by examining if genes with sex-biased expression are enriched on the X chromosome, as the evolution of sex-biased expression is thought to be driven primarily by sexually antagonistic selection (Ellegren & Parsch, 2007; Innocenti & Morrow, 2010; Griffin *et al*., 2013). Finally, we examined if the genes on the X chromosome are dosage compensated by comparing male and female gene expression in three composite tissues (heads, legs, and reproductive tracts). Our study thus provides a detailed study of several key aspects of X chromosome evolution in a previously unstudied group, revealing that the characteristics of these X chromosomes are conserved over at least 30 million years of evolution.

## Materials and methods

### Sample collection and sequencing

We used a combination of available genomic data from females in addition to newly collected data for males for five sexually reproducing species of *Timema (T. bartmani, T. cristinae, T. californicum, T. podura,* and *T. poppensis).* Reads from five females per species were downloaded from NCBI (Bioproject accession number: PRJNA670663). We collected four males from each of the five species from natural populations in California, from the same (or a geographically very close) population as the available females (Table S8). DNA extractions were done on whole-body adult males using the Qiagen Mag Attract HMW DNA kit following the manufacturer instructions. Sequencing libraries were generated for each male using a TruSeq DNA nano prep kit (550bp insert size). Libraries were then sequenced using Illumina HiSeq 2500 at the Lausanne Genomic Technologies Facility.

### Using coverage to identify X-linked scaffolds

Reads were trimmed before mapping using Trimmomatic (v. 0.36) (Bolger *et al*., 2014) to remove adapter and low-quality sequences (options: ILLUMINACLIP:3:25:6 LEADING:9 TRAILING:9 SLIDINGWINDOW:4:15 MINLEN:90). Reads from each individual were mapped to their species’ reference genome (Jaron *et al*., 2022) (Bioproject accession number: PRJEB31411) using BWA-MEM (v. 0.7.15) (Li, 2013). Multi-mapping and poor quality alignments were filtered (removing reads with XA:Z or SA:Z tags or a mapq < 30). PCR duplicates were removed with Picard (v. 2.9.0) (http://broadinstitute.github.io/picard/). Coverage was then estimated for all scaffolds at least 1000 bp in length using BEDTools (v. 2.26.0) (Quinlan & Hall, 2010). Per base coverage distributions were inspected visually for each individual and libraries with extremely non-normal coverage distributions were excluded from further analysis (Fig. S1-S5).

To compare coverage between males and females, coverage was first summed for all male and all female libraries per scaffold (for scaffolds at least 1000 bp in length). Male and female coverage was then normalised by modal coverage to adjust for differences in overall coverage. X-linked scaffolds were then identified using the log2 ratio of male to female coverage. Autosomal scaffolds should have equal coverage in males and females (log2 ratio of male to female coverage ≈ 0) and X-linked scaffolds should have half the coverage in males as in females (log2 ratio of male to female coverage ≈ −1). Each species showed frequency peaks near these values (Fig. S6). X linked scaffolds were classified in two ways: a ‘liberal’ classification (following (Vicoso & Bachtrog, 2015)) whereby scaffolds were classified as X-linked if the log2 ratio of male to female coverage < autosomal peak - 0.5, and a ‘stringent’ classification (following (Pal & Vicoso, 2015)) whereby scaffolds were classified as X-linked if the log2 ratio of male to female coverage was within 0.1 of the value of the X-linked peak.

To compare X-linked scaffolds between species, we examined the overlap of one-to-one orthologs (previously identified in (Jaron *et al*., 2022)) on the X-linked scaffolds. To determine if the overlap of X-linked orthologs between species was greater than expected, we used the SuperExactTest package (v. 0.99.4) (Wang *et al*., 2015) in R (R Core Team, 2017). Additionally, we used a tblastn approach to identify which regions of the *Bacillus rossius* genome correspond to the 210 shared X linked orthologs in *Timema* (maximum e-value = 1 x 10^-20^, minimum query coverage = 50%).

### Heterozygosity, nucleotide diversity, effective population size

To calculate heterozygosity, reads were mapped to the reference genomes as described above. We then additionally performed indel realignment with GATK (v. 3.7) (Van der Auwera *et al*., 2013). A maximum-likelihood estimate of the number of heterozygous sites per scaffold was then calculated using AngsD (v. 0.921) (Korneliussen *et al*., 2014) (options: -doSaf 1 -gl 1 -minQ 20 -minMapQ 40 -fold 1 -doCounts 1, with a minimum depth of 5 and a maximum depth of twice the median genome coverage once sites with 0 coverage were excluded) for each sample (for all scaffolds at least 1000 bp in length). The proportion of heterozygous sites was then calculated for each scaffold and the median proportion of heterozygous sites was weighted by the number of covered sites on a scaffold. We used a weighted median for scaffolds as shorter contigs are typically enriched for repeats and therefore less likely to give reliable estimates of heterozygosity. A similar approach was used to calculate pairwise nucleotide diversity for each species, but using only female samples (options: -doSaf 1 -gl 1 - minQ 20 -minMapQ 40 -fold 1 -doCounts 1 with a minimum depth of 5 and a maximum depth of twice the median genome coverage once sites with 0 coverage were excluded). The number of individuals (-nind) and the minimum number of individuals a site must be present in (-minind) were both set to the number of samples analysed. Effective population size (Ne) was estimated for the X and autosomes using the estimates of nucleotide diversity, assuming that nucleotide diversity is equal to 4Neμ (μ = mutation rate). Estimates of the mutation rate were taken from *Drosophila melanogaster* (2.8 x 10^-09^) (Keightley *et al*., 2014) and *Heliconius melpomene* (2.9 x 10^-09^) (Keightley *et al*., 2015).

### Excluding male achiasmy

An absence of recombination in males could influence the genetic diversity of the X and autosomes. Previous work in different phasmid species has shown that males do recombine (White, 1976; Marescalchi *et al*., 1986), however, this has not been shown in *Timema.* To examine this, we re-surveyed images of Giemsa-stained chromosome preparations from (Schwander & Crespi, 2009) which were originally generated for karyotyping purposes. Specifically we looked for the presence and position of chiasmata on pre-metaphase plates obtained from male testes.

### Selection analyses

We took values of dN/dS (number of non-synonymous substitutions per non-synonymous site / number of synonymous substitutions per synonymous site) for each species branch from Jaron et al (2022). Briefly, branch-site models with rate variation at the DNA level (Davydov et al., 2019) were run using the Godon software (https://bitbucket.org/Davydov/godon/, version 2020-02-17, option BSG --ncat 4) for each gene with an ortholog found in at least three of the species of *Timema* used here. Godon estimates the proportion of sites evolving under purifying selection (p0), neutrality (p1), and positive selection. We used only sites evolving under purifying or neutrality to calculate dN/dS. To test for differences in dN/dS between the X and autosomes, we used Wilcoxon tests with p-values adjusted for multiple comparisons using Benjamini and Hochberg’s algorithm (Benjamini & Hochberg, 1995). GC was calculated using the seqinR package (v.4.2-8) (Charif & Lobry, 2007) in R. We examined the occurrence of positive selection with two approaches. First, as in Jaron et al (2022), we used the branchsite models as described above to identify branches with evidence for positively selected sites. However, such branch-site models may lack power to detect positive selection so in addition we also used GODON to perform site-based model comparisons. Specifically, we compared the fit of model M8 (which allows for positively selected sites) and M8a (which does not allow for any positively selected sites) (Yang *et al*., 2000), also with rate variation at the DNA level (Davydov et al., 2019). The two models were compared using a log-likelihood ratio test for each gene. p-values were corrected for multiple tests using Benjamini and Hochberg’s algorithm. We then used a Fisher’s exact test to determine if positively selected genes were overrepresented on the X.

### Gene expression analyses

RNA-seq reads from three composite tissues (heads, legs and reproductive tracts) for males and females (3 replicates per sex) for each of the five species are publically available (Parker *et al*., 2019a; b) (Bioproject accession number: PRJNA392384). Adapter sequences were removed using Cutadapt (v. 2.3) (Martin, 2011) before quality trimming reads with Trimmomatic (v. 0.36) (Bolger *et al*., 2014) (options: LEADING:10 TRAILING:10 SLIDINGWINDOW:4:20 MINLEN:80). Trimmed reads were then mapped to reference genomes using STAR (v. 2.6.0c, default options). HTSeq v.0.9.1 (Anders *et al*., 2015) was used to count the number of reads uniquely mapped to the exons of each gene, with the following options (htseq-count --order=name --type=exon --idattr=gene_id -- stranded=reverse). Expression analyses were performed using the Bioconductor package EdgeR (v 3.32.1) (Robinson *et al.*, 2010), and done separately for each species and tissue. Normalisation factors for each library were computed using the TMM method and were used to calculate normalised expression levels (either FPKM (Fragments Per Kilobase of transcript per Million mapped reads) or TPM (Transcripts Per Million mapped reads)). For the main analyses, genes with low expression (less than 2 FPKM (or TPM) in 2 or more libraries per sex) were excluded. This filtering step was used to exclude any sex-specifically expressed genes as our goal was to examine how the expression of genes differs in males and females. This decision could influence our results if sex-limited gene expression was extensive on the X chromosome. To investigate this we repeated our analysis with the inclusion of sex-limited genes and found similar results to the main analyses (Fig. S7 - S8). To examine dosage compensation, we used the log2 ratio of mean male expression level to female expression level and used Wilcoxon tests (adjusted for multiple testing using Benjamini and Hochberg’s algorithm (Benjamini & Hochberg, 1995)) to determine if this ratio differed between genes on the autosomal and X-linked scaffolds.

To determine the significance of sex on gene expression, we fit a generalised linear model (GLM) with a negative binomial distribution with sex as an explanatory variable and used a GLM likelihood ratio test to determine the significance for each gene. P-values were then corrected for multiple tests using Benjamini and Hochberg’s algorithm (Benjamini & Hochberg, 1995). In the main analysis sex-biased genes were then defined as genes that showed difference in expression between males and females with a false-discovery rate (FDR) < 0.05 (we also repeated analyses with the additional condition that genes must show a greater than two fold difference in expression to ensure our results are robust to the effects of sex-biased allometry (Montgomery & Mank, 2016)). Note that all genes not classified as sex-biased were classified as unbiased genes. We then examined if male- or female-biased genes were under-or over-represented on autosomal or X-linked scaffolds using Fisher’s exact tests (adjusted for multiple-testing using Benjamini and Hochberg’s algorithm (Benjamini & Hochberg, 1995)). To examine species by sex interactions in gene expression, we used a similar similar GLM approach as above, with sex, species, and species by sex interaction as explanatory variables for genes with an ortholog in each species. Genes with an FDR < 0.1 for the interaction term were considered to have significant species by sex interaction. To determine if sex-biased genes on the X were underrepresented for species by sex interactions, we used a Fisher’s exact test.

It is possible that dosage compensation may vary along the X chromosome. Unfortunately, our current reference genomes are too fragmented to investigate this question. Previous work by Nosil et al. (Nosil *et al*., 2018) however produced a more contiguous genome assembly of one of our study species, *T. cristinae,* which has been further constructed into a linkage map. To use this synteny information for each of our species, we anchored the scaffolds from each of our genome assemblies to the Nosil et al. (Nosil *et al*., 2018) reference genome (BioProject Accession PRJNA417530) using MUMmer (version 4.0.0beta2) (Marçais *et al*., 2018) with parameter --mum. The alignments were processed by other tools within the package: show-coords with parameters -THrcl to generate tab-delimited alignment files and dnadiff to generate 1-to-1 alignments. We used only uniquely anchored scaffolds for which we were able to map at least 10k nucleotides to the Nosil et al. (2018) reference genome. Nosil et al (2018) indicated that linkage group 13 was the X chromosome ((Nosil *et al*., 2018)). To determine if this was correct, we repeated our coverage analyses on the Nosil et al. (2018) assembly. From this, we found that most scaffolds that make up linkage group 13 did not show reduced coverage in males (Fig. S9). In addition, several scaffolds from other linkage groups did show reduced coverage in males (Fig. S9). In order to use as much of the synteny information as possible we “cleaned” the Nosil et al. (2018) assembly by removing X-linked scaffolds from autosomal linkage groups 1-12. These scaffolds were then assigned to a new, unordered collection of scaffolds from the X chromosome, together with X-linked scaffolds from linkage group 13 and from those not assigned to any linkage group in Nosil et al. (2018). Scaffolds from linkage group 13 that were not X-linked were assigned to linkage group NA. We classified scaffolds in the Nosil et al. (2018) assembly as X-linked if most of a scaffold was covered by aligned scaffolds from our assembly assigned to the X rather than autosomes (i.e. if aligned scaffolds assigned as X covered more than twice as many bases as those assigned as autosomal for a particular scaffold, it was classed as X-linked).

## Results

### Identifying X-linked scaffolds

We used a coverage approach to identify X-linked scaffolds in our previous genome assemblies (Jaron *et al*., 2022). Genomic reads from four males and five females from each species were mapped onto the corresponding reference genome. After filtering low-quality alignments and non-uniquely mapping reads (see Methods) the median coverage per sample ranged from 11x to 31x (18.5x on average, see Table S1). Visual inspection of coverage distributions (Figs S1-S5) found that while most libraries had either one (in females) or two (in males) coverage peaks, three libraries (*T. podura* (H56, Fig. S4), *T. poppensis* (ReSeq Ps08 and Reseq_Ps12, Fig. S5)) did not show a clear coverage peak, and were excluded from all further analyses.

To identify X-linked scaffolds, we used the log2 ratio of (normalised) coverage of males to females. As males have only a single copy of the X and females have two, X linked scaffolds should have twice as much coverage in females as males (log2 male:female coverage ≈ −1). Autosomes are expected to have the same coverage in both sexes (log2 male:female coverage ≈ 0). Considering all scaffolds with a log2 ratio of male to female coverage < autosomal peak −0.5 to be X-linked, we classified between 12 and 14 % of each genome as X-linked (Table 1, Fig. S6), which fits well with the X chromosome size observed in karyotypes (Schwander & Crespi, 2009). This approach may mean some autosomal contigs may be misclassified as X-linked, thus we also repeated all our analyses with a more stringent classification scheme (scaffolds with a log2 ratio of male to female coverage within 0.1 of the X linked peak) (Table S2, Fig. S6). Using the more stringent classification scheme produced very similar results (not shown). Of note, most differences between the classification schemes are for short scaffolds (1000-4999 bp), which represent ~20% of the genome assemblies. When these are excluded the two classification schemes classify almost the same set of scaffolds as X-linked (Fig. S10, Table S3). Finally, we also examined the heterozygosity of the X in males. As expected, the heterozygosity of the X is close to zero and much lower than on the autosomes, corroborating our X-linked scaffold assignments (Fig. S11).

**Table 1.** Number of genes and scaffolds classified as X-linked or autosomal

| Species | Min scaffold length | % genome X-linked | N Autosomal genes | N X-linked genes | N genes not classified |
| --- | --- | --- | --- | --- | --- |
| *T. bartmani* | 1000 | 12.11 | 12804 | 1244 | 18 |
| *T. cristinae* | 1000 | 12.21 | 12542 | 1316 | 24 |
| *T. californicum* | 1000 | 12.22 | 13207 | 1344 | 12 |
| *T. podura* | 1000 | 13.75 | 15069 | 1457 | 3 |
| *T. poppensis* | 1000 | 12.40 | 14115 | 1454 | 36 |

### The X chromosome is conserved across *Timema*

Comparing orthologs across the five different species shows X chromosome gene content is conserved, with >90% of X-linked orthologs shared between the 5 species (Fig. 1, Fig. S12). This overlap is much greater than expected by chance (FDR < 6.863 x 10^-316^). This suggests that the X chromosome is homologous in all five species, which last shared a common ancestor approximately 30 million years ago (Riesch *et al*., 2017). Additionally, we used species genome alignments to an independent *T. cristinae* genome assembly (Nosil *et al*., 2018) to assign scaffolds to linkage groups based on a single reference. By applying coverage analyses we were able to identify and correct the X linked scaffolds in the Nosil et al. 2018 assembly (see Methods). Using this corrected reference, we found that contigs aligned to X-linked scaffolds showed reduced coverage in males but not females in each species (Figs S13-S17), again indicating the X chromosome is the same in all species. Finally, using BLAST we found that the majority (72%) of the shared X-linked genes in *Timema* (for which we were able to obtain a significant hit) were also present on the X chromosome of *Bacillus rossius* (Fig. S18). The split between *Timema* and all other extant phasmids (the Euphasmatodea, which includes *Bacillus)* occurred approximately 120 mya (Simon *et al*., 2019), suggesting that the X chromosome in phasmids predates this split.

**Fig. 1.**
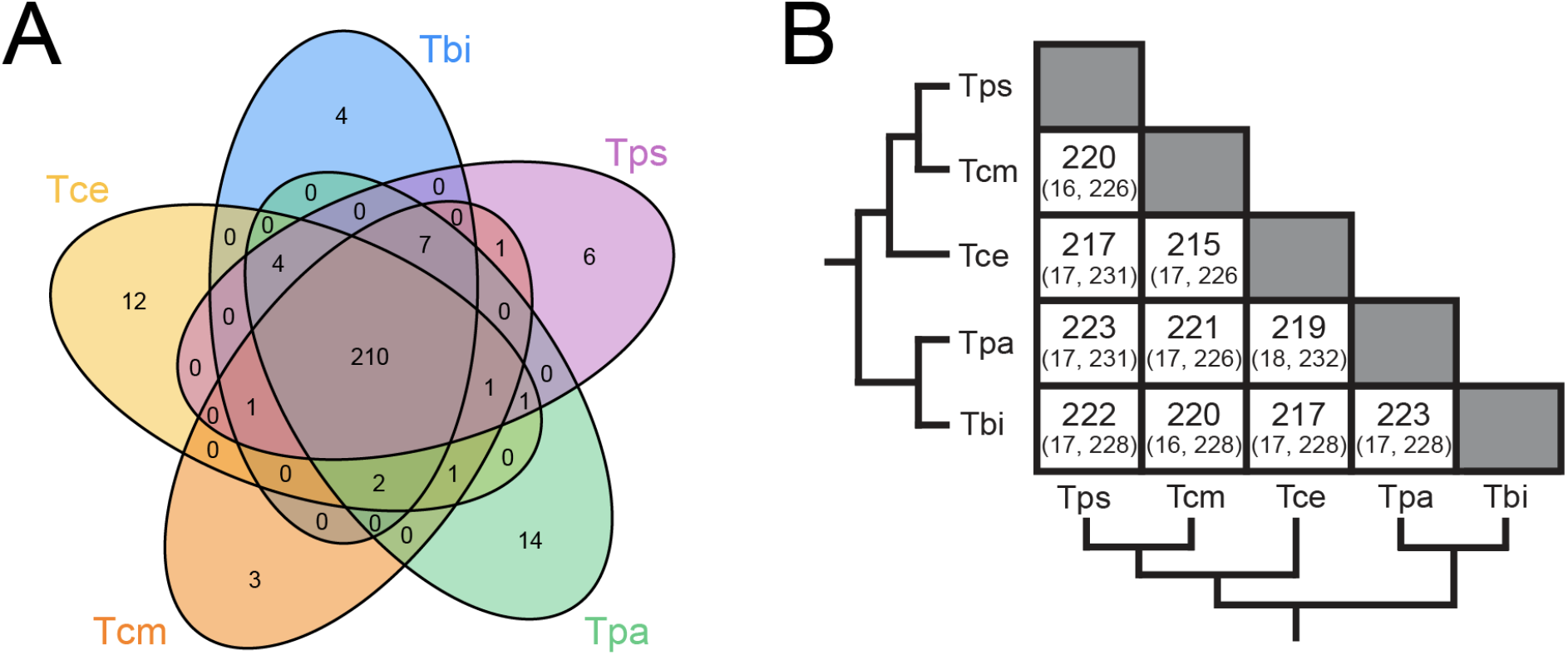
The X chromosome is conserved between *Timema* species. **A**. Venn-diagrams showing the number of shared X-linked orthologs between species. **B**. Number of shared orthologs (expected, maximum possible). The observed amount of overlap was much greater than expected in all comparisons (FDR < 6.863 x 10^-316^). Species names are abbreviated as Tbi = *T. bartmani,* Tce = *T. cristinae,* Tcm = *T. californicum,* Tps = *T. poppensis,* and Tpa = *T. podura.*

### The X chromosome has reduced genetic variation

We used female samples to examine two related measures of genetic variation: heterozygosity and nucleotide diversity (π). Only female samples were used, as males are hemizygous for the X, which would affect the estimation of heterozygosity on the X directly. It would also affect the estimation of nucleotide diversity indirectly, as X haplotypes containing recessive lethals on the X will be absent in male samples. We found that both heterozygosity and nucleotide diversity (π) were lower on X-linked than on autosomal scaffolds (Fig. S11, S19), as expected. In all species, the ratios of X to autosomes for both of these measures (heterozygosity = 0.16 to 0.62 (Fig. 2), π = 0.19 to 0.48 (Fig. 2)) were lower than the 0.75 that would be expected from the reduced effective population size of the X relative to the autosomes. This pattern was also seen when comparing the X to each of the autosomal linkage groups individually (Fig. S20, S21). Nucleotide diversity estimates were then used to estimate the effective population size of each species. From this, we estimated the autosomal effective population size in *Timema* to range from ~150,000 (*T. poppensis)* to ~2,000,000 (*T. podura)* (Table S4). Finally, we corroborated that male meiosis was chiasmate, which was revealed by the presence of at least one chiasma per autosome on pre-metaphase plates (Fig. S22).

**Fig. 2.**
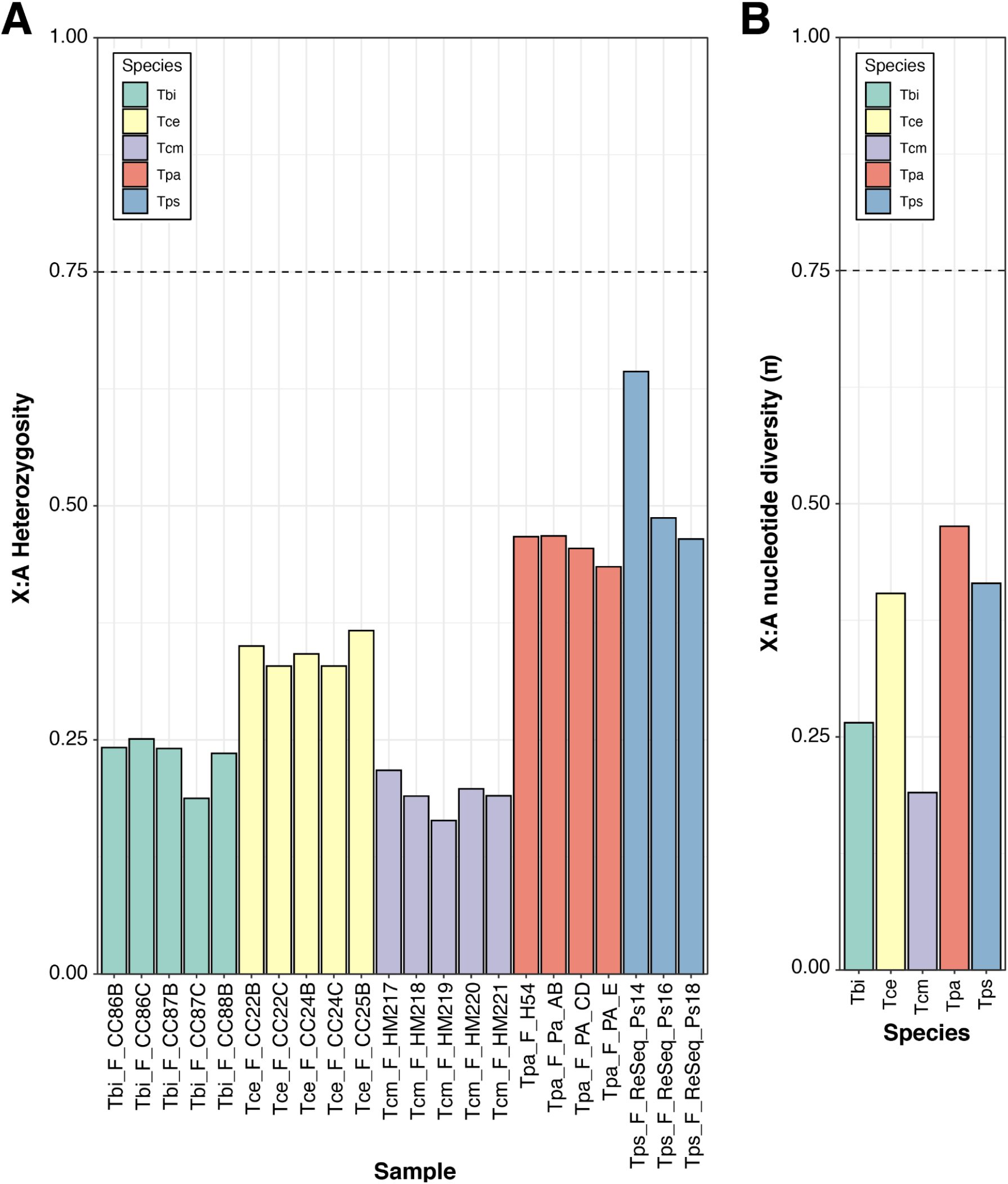
Genetic variation on the X chromosome is lower than expected. **A.** Ratio of heterozygosity on the X and autosomes in 3 to 5 females per species. **B.** Ratio of pairwise nucleotide diversity (π) on the X and autosomes. Species names are abbreviated as Tbi = *T. bartmani,* Tce = *T. cristinae,* Tcm = *T. californicum,* Tps = *T. poppensis,* and Tpa = *T. podura.* The dotted lines indicate the neutral expectation in a population with a balanced sex ratio.

### The X chromosome shows evidence for reduced purifying selection but no differences in positive selection

Genes on the X chromosome show an elevated dN/dS relative to genes on the autosomes in each species (Fig. 3A). This effect remains when accounting for differences in GC and mean expression level (permutation ANOVA p < 0.05 for all species). This increased ratio appears to be driven by reduced purifying selection, as X-linked genes are not enriched for positively selected genes (Fig. 3B, Table S9).

**Fig. 3.**
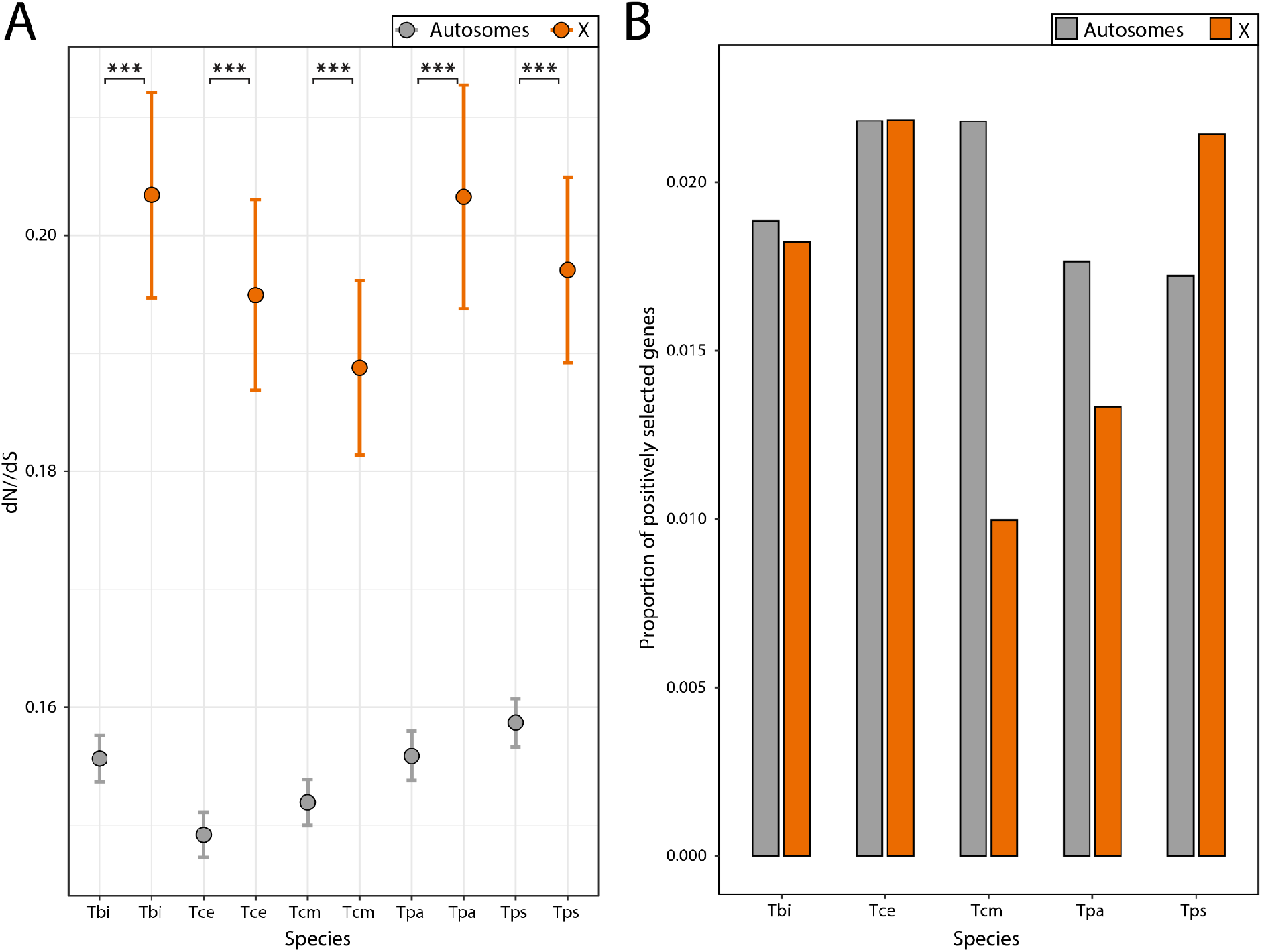
Sequence evolution on the X and autosomes. **A.** Average dN/dS across genes. Error bars indicate standard error. Asterisks indicate the significance (FDR) of Wilcoxon tests (***<0.001, **<0.01, *<0.05). **B.** Proportion of positively selected genes. Positively selected genes were not enriched on the X chromosome (Fisher’s exact test p value = 0.50). Species names are abbreviated as Tbi = *T. bartmani,* Tce = *T. cristinae,* Tcm = *T. californicum,* Tps = *T. poppensis,* and Tpa = *T. podura.*

Although dN/dS is elevated for genes on the X, overall divergence (branch length, dN + dS) is lower for genes on the X than genes on the autosomes (Fig. S23A). More specifically, dS is lower on the X for all species (Fig. S23B) and values of dN are only higher on the X for two species (*T. podura* and *T. poppensis*, Fig. S23C). The reduction in dS on the X is likely due to an overall lower mutation rate in the female than male germline, resulting in fewer mutations on the X due to the X’s more frequent transmission through females (Kirkpatrick & Hall, 2004; Ellegren, 2007). Taken together, this suggests that while the genes on the X are subject to reduced purifying selection, the overall divergence of X-linked genes is smaller.

### *Timema* show complete dosage compensation in heads and legs, but no dosage compensation in reproductive tracts

We examined if the X chromosome is dosage compensated in *Timema* by comparing gene expression in males and females in three different composite tissues (heads, legs, and reproductive tracts) for each of our five species. While for the main analysis we calculated expression values as FPKM, we also repeated our analyses using TPM. Analyses based on TPM showed very similar results and are provided as supplemental figures and tables. We examined the log2 ratio of male to female expression on the X and the autosomes. For the autosomes and a dosage compensated X, the log2 value should be approximately 0, whereas a non-dosage compensated X would have a value of approximately −1. We found that in heads and legs the ratio was close to 0 with only small differences between the X and the autosomes (Figs. 4, S24, S25), indicating almost-complete complete dosage compensation in these tissues. By contrast, in the reproductive tracts the ratio of male to female expression for genes on the X is close to −1 (Figs. 4, S24, S25), indicating an absence of dosage compensation in this tissue. This observation was also seen when comparing the X to each of the autosomal linkage groups individually (Figs. S26, S27). An alternative, mutually non-exclusive possibility is that the greatly reduced expression observed in genes on the X in male reproductive tracts is due to a large enrichment of female-biased genes and depletion of male-biased genes. Although we cannot formally exclude this possibility, three lines of evidence indicate that lack of dosage compensation in reproductive tracts is the best explanation for our findings. Firstly, a lack of dosage compensation is expected to result in a two-fold reduction of expression, meaning X linked genes in males should show a major peak with a two-fold reduction in expression (Mank & Ellegren, 2009; Vicoso *et al*., 2013; Pal & Vicoso, 2015), which is what we observe (Fig. 5). Secondly, if the X chromosomes facilitates the accumulation of sex-biased genes, we should be able to observe this effect in all tissues (Jaquiéry *et al*., 2021). While we find a large enrichment of female-biased genes and a depletion of male-biased genes on the X in the reproductive tracts, this is not found in the other tissues, with only a slight enrichment of female-biased genes in the heads of two species and a depletion of male-biased genes in one species (Fig. 6, Fig. S28, Table S5, Table S6). This suggests that selection for the enrichment of female-biased genes and the depletion of male-biased genes on the X is weak, and thus unlikely to generate the large effect sizes we see in the reproductive tracts. Finally, a reduction in expression of the male X due to a lack of dosage compensation is expected to be consistent across species, since they share the same X chromosome (see above). By contrast, sex biased expression independently of dosage is more likely to be species-specific given the very fast turnover of sex biased genes observed between closely related species in several taxa (Zhang *et al*., 2007; Harrison *et al*., 2015) including *Timema* (Parker *et al*., 2019b). To distinguish between these patterns, we tested if sex-biased genes on the X in the reproductive tracts are underrepresented for genes with species by sex interactions. We found that genes on the X are underrepresented for genes showing species by sex interactions in the reproductive tracts (p = 0.010), and that this is not the case for the heads (p = 0.938) or legs (p = 0.795). This shows that sex-differences in expression on the X are more consistent between species in the reproductive tracts than in the heads and legs (also see Fig. S29), and further supports a lack of dosage compensation in the reproductive tracts rather than a large enrichment of female-biased genes on the X.

**Fig. 4.**
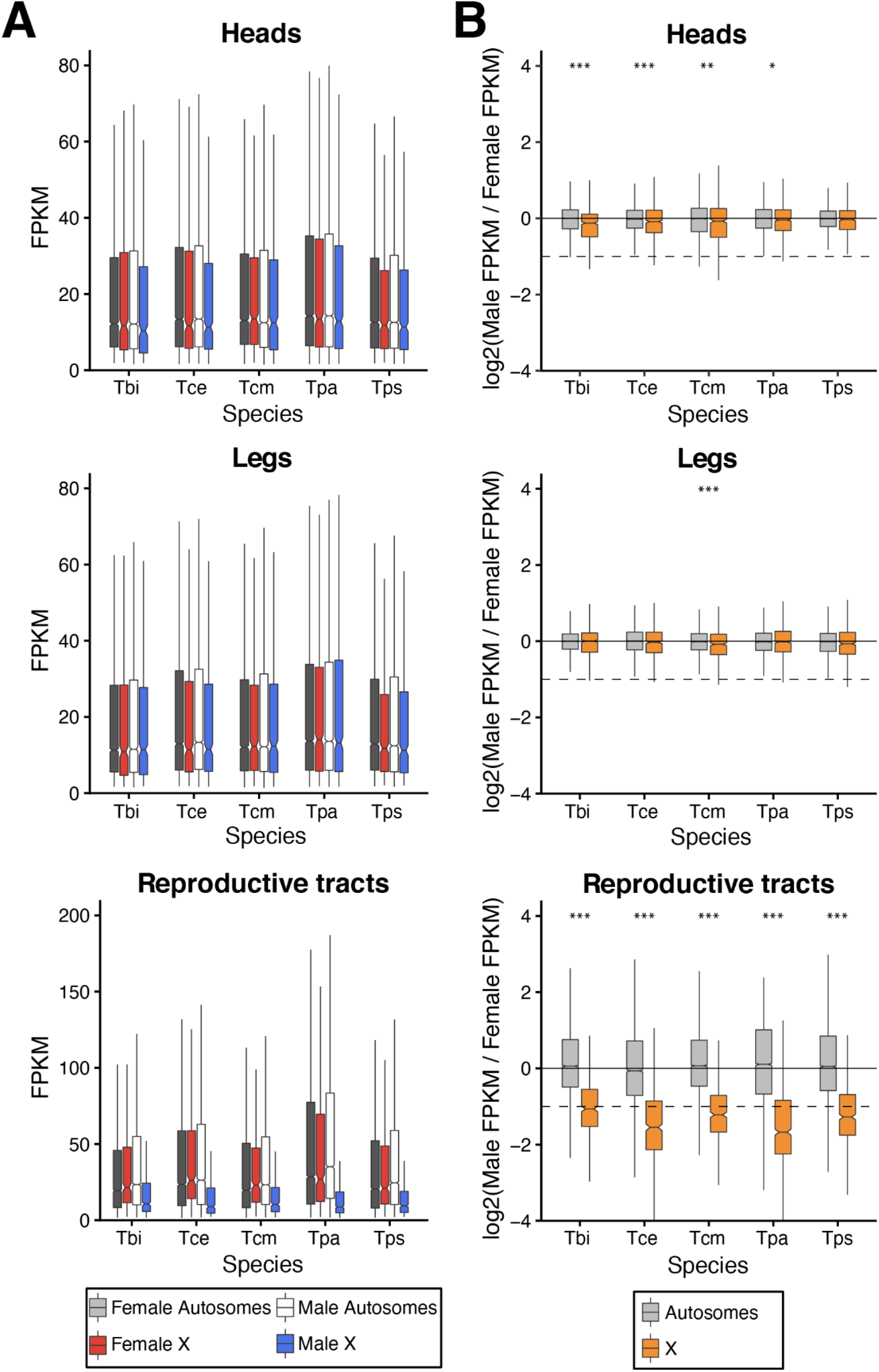
Gene expression on the X and autosomes in heads, legs, and reproductive tracts. **A.** Average expression levels in males and females on the X and autosomes **B.** Log2 of male to female expression ratio for the X and autosomes. Dashed lines represent a twofold reduction in expression in males (as expected if there was no dosage compensation). Species names are abbreviated as Tbi = *T. bartmani*, Tce = *T. cristinae*, Tcm = *T. californicum*, Tps = *T. poppensis,* and Tpa = *T. podura.*

**Fig. 5.**
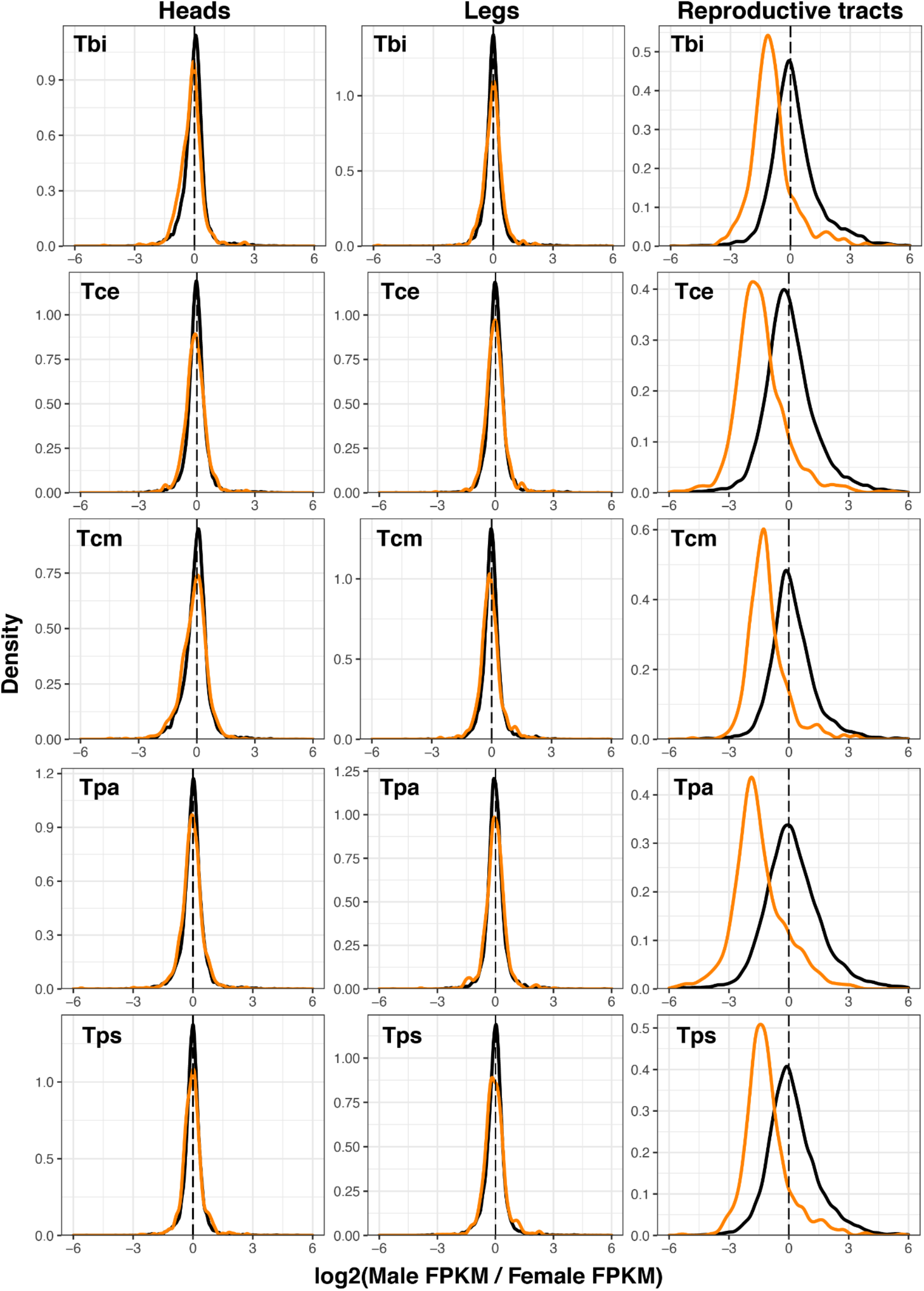
Ratio of male and female gene expression on the X (orange) and autosomes (black) in heads, legs, and reproductive tracts. Species names are abbreviated as Tbi = *T. bartmani,* Tce = *T. cristinae,* Tcm = *T. californicum,* Tps = *T. poppensis,* and Tpa = *T. podura*.

**Fig. 6.**
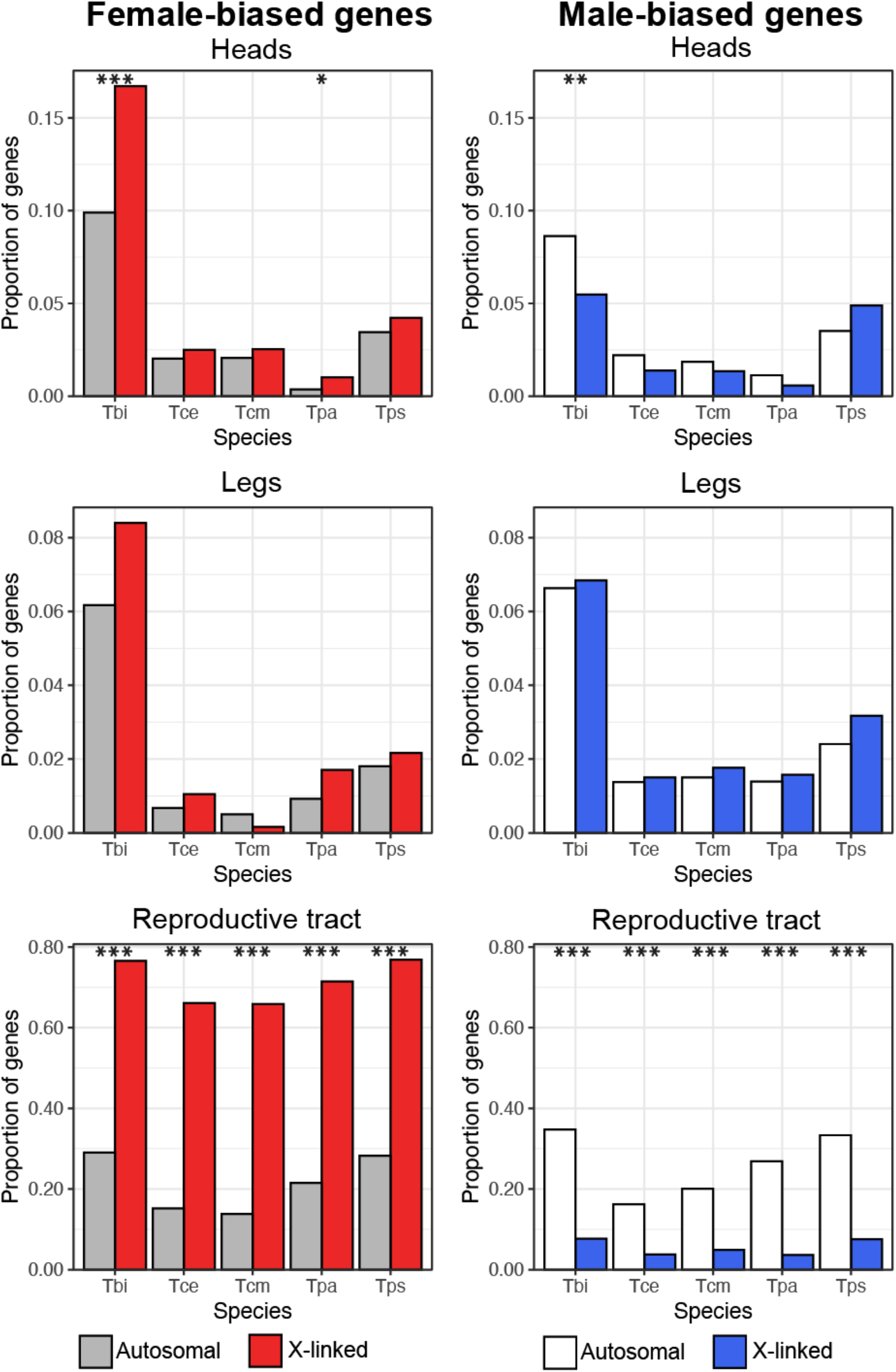
Proportion of female- and male-biased genes on the X and autosomes in reproductive tract, head and leg samples. Note the scale changes between tissue-types. Asterisks indicate the significance level (FDR) of Fisher’s exact tests (***<0.001, **<0.01, *<0.05). Species names are abbreviated as Tbi = *T. bartmani,* Tce = *T. cristinae,* Tcm = *T. californicum,* Tps = *T. poppensis,* and Tpa = *T. podura.*

By examining the expression of the X and autosomes in the different tissues, we can infer the type of dosage compensation. In the dosage compensated tissues (heads and legs) genes on the autosomes and X have similar overall expression levels in both males and females, and the overall expression of the X is similar to that of the autosomes (Fig. 4, Fig. S24, Table S7). By contrast, in the reproductive tracts, where dosage compensation is lacking, expression of genes on the X is much lower in males than in females. This difference seems to be driven by changes in male expression, as X-linked gene expression in females remains similar to the autosomes (Fig. 4, Fig. S24, Table S7). This supports a mechanism of dosage compensation by hyper-transcription of the X in males, a mechanism common among other insects (Gu & Walters, 2017).

Although we find almost complete dosage compensation, it is possible that its extent could vary along the X chromosome (Mullon *et al*., 2015). To address this, we examined the extent of dosage along the longest X-linked scaffolds in the genome, and found no evidence of variation along the X chromosome (Fig. S30 - S34, see also Fig. S35 - S46 for expression variation along the autosomes).

### Sex-biased genes are not enriched on the X chromosome

Sexually antagonistic mutations are expected to fix more easily on the X chromosome than the autosomes. As the evolution of sex-biased gene expression is thought to be primarily driven by sexually antagonistic selection, it is expected that the X chromosome will be a hotspot for sex-biased gene expression (Rice, 1984; Gibson *et al*., 2002; Ellegren & Parsch, 2007; Innocenti & Morrow, 2010; Griffin *et al*., 2013). Contrary to this expectation, we find most sex-biased genes on the autosomes (% of sex-biased genes on the X ranges from 8.3 - 14.3, Table S5), and we find little evidence for enrichment of sex-biased genes on the X in the head and leg tissues (Fig. 6, Table S5, Table S6). Note that almost all genes on the X appear to be sex-biased in the reproductive tracts, but this effect is largely due to a lack of dosage compensation in this tissue (see above). In addition, while we do find an enrichment of female-biased genes in the heads of two species (*T. bartmani* and *T. podura,* Fig. 6) and a depletion of male-biased genes in *T. bartmani,* the effect sizes are small and the effect becomes insignificant when considering only sex-biased genes with at least a twofold difference in expression between males and females (Fig. S28). These findings suggest that the selective pressures to accumulate female-biased genes and reduce male-biased genes on the X are weak and/or that there are constraints to the gene content or expression levels on the *Timema* X chromosome.

## Discussion

The difference in copy numbers of the X chromosome in males and females is expected to have profound effects on its evolution. In particular, the X chromosome is predicted to evolve at a faster rate due to a combination of hemizygous selection and increased drift (Wright, 1931; Charlesworth *et al*., 1987; Vicoso & Charlesworth, 2009), to accumulate sexually antagonistic alleles (Rice, 1984; Gibson *et al*., 2002), and to evolve dosage compensation mechanisms (Disteche, 2012; Mank, 2013; Gu & Walters, 2017; Lenormand *et al.*, 2020). Support for these predictions has been mixed from the taxa examined so far, yet the factors responsible for the variation among different taxa are poorly understood (Bachtrog *et al*., 2011; Gu & Walters, 2017). In this study, we examine these predictions across five species of *Timema* stick insects. Overall, we find evidence for relaxed selection and complete dosage compensation in somatic tissues, but little evidence for the accumulation of sexually antagonistic alleles (in the form of sex-biased genes) on the X. Patterns of X chromosome evolution were generally consistent across *Timema* species, suggesting that the factors influencing sex chromosome evolution in this group are also largely the same.

Sex chromosome conservation is highly variable between taxa, with extensive turnover between species in some groups, e.g. beetles (Coleoptera) (Blackmon & Demuth, 2014), flies (Diptera) (Vicoso & Bachtrog, 2015), or frogs (Ranidae) (Jeffries *et al*., 2018), and conservation for over a hundred million years in others, e.g. Eutherian mammals (Lahn & Page, 1999; Cortez *et al*., 2014; Marshall Graves, 2015), moths and butterflies (Lepidoptera) (Fraïsse *et al*., 2017), or birds (Shetty *et al*., 1999; Xu & Zhou, 2020). The factors influencing turnover rate are complex and interacting (Vicoso, 2019), however, one key factor is the level of differentiation between the X and Y chromosomes, with more differentiated chromosomes less likely to turnover (Pokorná & Kratochvíl, 2009; Vicoso, 2019). Our finding that the X chromosome in *Timema* is old (likely conserved for over 120 million years) supports this idea. XX/X0 sex determination systems as found in *Timema* (Schwander & Crespi, 2009) are thought to derive from XX/XY systems with highly differentiated X and Y chromosomes and represent the end point of the gradual loss of gene content from the Y (Bergero & Charlesworth, 2009). As such, XX/X0 systems could be considered the most extreme example of sex chromosome differentiation possible, which may mean that XX/X0 systems are particularly unlikely to turnover (but see (Blackmon & Demuth, 2014)).

Gene sequence evolution on the X chromosome is expected to be faster than that on the autosomes due to a combination of increased drift and hemizygous selection (Charlesworth *et al*., 1987; Vicoso & Charlesworth, 2009). While this prediction should apply universally, empirical support is mixed (Mank *et al*., 2010; Meisel & Connallon, 2013; Charlesworth *et al*., 2018; Pinharanda *et al*., 2019; Whittle *et al*., 2020). The cause of this variation is unclear, in particular, because typically only single lineages are examined. In single lineages it is difficult to disentangle the influence of X-linkage from lineage-specific effects such as recent bottlenecks, differences in operational sex ratio, or population size. By examining the influence of X-linkage in multiple *Timema* species covering a span of 30 million years of divergence, we can assess how consistent the effects of X linkage are, allowing for a comprehensive assessment of X chromosome evolution in this genus. Using this approach, we found consistent evidence for a relaxed selection (as measured by dN/dS), known as the “faster X effect”, in all species, and that this effect is primarily driven by reduced purifying selection on the X. Interestingly, while dN/dS was higher for genes on the X, the overall amount of divergence was lower. This result is likely driven by a lower net mutation rate on the X relative to autosomes as a result of a lower mutation rate in the female than male germline (Kirkpatrick & Hall, 2004; Ellegren, 2007).

The reduction of purifying selection on the X relative to the autosomes is expected to be largest when the effective population size of the X (Ne_x_) is much smaller than that of the autosomes (Ne_A_) (Vicoso & Charlesworth, 2009). The neutral expectation in a population with a balanced sex ratio is that the ratio of Ne_x_ to Ne_A_ will be approximately 0.75 (Wright, 1969; Hartl & Clark, 2006), however, demographic and selective processes can have a large effect on this ratio (Nunney, 1993; Caballero, 1995; Charlesworth, 2001). In *Timema*, Ne_x_ / Ne_A_ values are much smaller than 0.75 (0.19 to 0.48, estimated from nucleotide diversity), meaning the effective population size of the X is much smaller than that of the autosomes. Departures from the expected 0.75 nucleotide diversity ratio are common across animals and are typically thought to be a consequence of sex-biased demography (Mank *et al*., 2010). One common cause for such departures is that in many systems reproductive skew is stronger for males than females, meaning fewer males contribute to the next generation than females. In *Timema,* females mate multiply and skew offspring towards particular males (Arbuthnott *et al*., 2015), suggesting that in *Timema* reproductive skew is indeed greater in males than females. In male heterogametic species (such as *Timema*), such a reproductive skew should disproportionately reduce diversity on the autosomes, and result in an X to autosome nucleotide diversity ratio > 0.75. Here however, we observe the opposite pattern, a reduction in the X to autosome nucleotide diversity ratio. As such, the reduced diversity on the X we observe is unlikely to be due to sex-specific variation in reproductive success. There are many potential alternative demographic and selective explanations for this finding, however the most likely (discussed below) are that the X has a lower recombination rate than the autosomes, and/or that *Timema* undergo frequent population bottlenecks. Since we show that crossing over occurs in both sexes, the effective recombination rate for the X is likely lower than for the autosomes (as recombination between different copies of the X can only occur in females). Low recombination rates intensify the consequences of selective sweeps and background selection on genetic diversity and could thus contribute to the reduced genetic diversity on the X we observe (Betancourt *et al*., 2004; Charlesworth, 2012, 2013; Wilson Sayres, 2018). Population bottlenecks could also contribute to the reduced genetic diversity on the X. Although bottlenecks will reduce genetic diversity for both the X and the autosomes, it is expected that they will disproportionately reduce genetic diversity on the X (Pool & Nielsen, 2007). This factor may be particularly relevant as *Timema* live in fire-prone habitats where fires may lead to frequent population bottlenecks.

Independently of the mechanisms responsible for the strongly reduced effective population size of the X in *Timema*, it supports our interpretation that the faster-X effect is driven by less effective purifying selection. Most previous studies of the faster-X effect have focused on species with Ne_x_ / Ne_A_ values ≥ 0.75 (Mank *et al*., 2010), making direct comparisons with our study difficult. The exceptions to this are studies on birds and *Heliconius* butterflies which have values of Ne_z_ / Ne_A_ in a similar range as *Timema.* Note that both these taxa have heterogametic females (ZW), where low Ne_z_ / Ne_A_ ratios are believed to stem from strong reproductive skew among males (Vicoso & Charlesworth, 2009; Mank *et al.*, 2010). In birds, a faster-Z (faster-X) effect is commonly observed and appears to be driven primarily by less effective purifying selection (Mank *et al*., 2010). In contrast, in *Heliconius* evidence for a faster-Z effect is weaker and is thought to be driven by increased levels of adaptive evolution on the Z, with no evidence for reduced purifying selection on the Z (Pinharanda *et al*., 2019). The cause of this difference is unclear. However, it has been suggested that it may be due to the overall higher effective population size in *Heliconius* (Ne ≈ 2,000,000 (Keightley *et al*., 2015)) compared to birds (Ne = 200,000 - 600,000 (Primmer *et al*., 2002; Axelsson *et al*., 2004; Jennings & Edwards, 2005; Backström *et al*., 2008)), meaning that selection can be efficient in *Heliconius* even with the relatively reduced effective population size of the sex chromosome (Mank *et al*., 2010; Pinharanda *et al*., 2019). This interpretation conflicts with our findings in *Timema,* as the (autosomal) effective population size varies greatly between species (from ~150,000 in *T. poppensis* to ~2,000,000 in *T. podura*), yet we observe a similar reduction in purifying selection on the X in all species. The reason for this variation between study systems is thus unclear, and highlights the need for future studies across diverse taxa. It should be noted that the values of Ne we estimate for *Timema* should be considered as rough approximations as our calculations of Ne assume the mutation rate in *Timema* is similar to that in *D. melanogaster* and *H. melpomene.* Since these are distantly related species the mutation rate in *Timema* could differ markedly, which would mean our estimates of Ne would be systematically under- or over-estimated. While this complicates direct comparisons with other taxa, it is unlikely to influence our comparisons among *Timema* species for which only the relative differences in Ne are important.

The X chromosome has long been predicted to be a hotspot for sexually antagonistic variation (Rice, 1984; Gibson *et al*., 2002). The evolution of sex-biased gene expression is thought to be driven by sexually antagonistic selection, and thus sex-biased gene expression should be overrepresented on the X chromosome ((Ellegren & Parsch, 2007; Innocenti & Morrow, 2010; Griffin *et al.*, 2013) but see (Hitchcock & Gardner, 2020; Ruzicka & Connallon, 2020)). In *Timema,* we find very little support for this prediction with only a small enrichment of sex-biased genes on the X in one of our five species. In combination with studies in *Drosophila melanogaster* (Ruzicka *et al*., 2019), *Callosobruchus maculatus* (Sayadi *et al*., 2019), and *Ischnura elegans* (Chauhan *et al*., 2021) which also found little or no enrichment of sexually antagonistic alleles or sex-biased gene expression on the sex chromosomes, our study suggests that the advantage of accumulating sexually antagonistic alleles on the X be may be smaller than often assumed. The reasons for this are unclear. However, it is likely that the advantage X-linkage gives to sexually antagonistic alleles is balanced by other forces such as epistatic interactions (Arnqvist *et al*., 2014) or sex-specific dominance (Fry, 2010). Both forces favour the accumulation of sexually antagonistic alleles on the autosomes. Additionally, it should be noted that our approach to use sex-biased genes as a proxy for sexually antagonistic variation may be underpowered as it will miss any genes that are evolving under sexual conflict but have not (yet) evolved sex-biased gene expression. Despite this, several studies do show the expected enrichment of sexually antagonistic alleles or sex-biased gene expression on the X: in *Tribolium castaneum* (Whittle *et al*., 2020), Diptera (Innocenti & Morrow, 2010; Vicoso & Bachtrog, 2015), Hemiptera (Pal & Vicoso, 2015), and nematodes (Albritton *et al*., 2014). A key challenge for future work will thus be to integrate studies that quantify multiple factors that influence the accumulation of sexually antagonistic alleles (e.g. epistatic interactions, sex-specific dominance, reproductive skew, etc.) to understand how the variation between studies and taxa is produced.

In many species dosage compensation is thought to be important for ameliorating the costs of misexpression of genes on the X chromosome (Marín *et al*., 2000). Although common, there is a great deal of variation in the extent to which genes are dosage compensated. Here we find almost complete dosage compensation in the somatic tissues of all five *Timema* species, as reported for other species with X0 systems (e.g. Nematodes (Meyer, 2000) and crickets (Rayner *et al*., 2021)). Dosage compensation is expected for X0 systems as they are thought to arise from XY systems where the Y chromosome has degraded to the point it can be lost without a large decrease in fitness. By this stage, most genes on the X should already be haplo-sufficient in males. In addition, the evolution of dosage compensation could itself hasten the loss of the Y chromosome, as genes that have functional copies on both the X and the Y will be misexpressed if chromosome-wide dosage compensation evolves (Vicoso & Bachtrog, 2009; Lenormand & Roze, 2021).

In contrast to the somatic tissues, male reproductive tracts displayed a lack of dosage compensation. Reduced expression of X chromosome in male reproductive tissue has been observed in a number of species including mammals (Khil *et al*., 2004; Disteche, 2012; Sangrithi & Turner, 2018), *C. elegans* (Kelly *et al*., 2002; Pirrotta, 2002) and insects such as *D. melanogaster* (Oliver, 2002; Meiklejohn *et al*., 2011; Mahadevaraju *et al*., 2021) or *Teleogryllus oceanicus* (Rayner *et al*., 2021). Whether reduced X expression in male reproductive tracts generally stems from a lack of dosage compensation is, however, not clear. Indeed, reduced X expression can be caused by several non-mutually exclusive mechanisms, including the accumulation of female-biased genes on the X (Mank, 2009), the movement of male-beneficial genes to the autosomes (Vibranovski *et al*., 2009), or the inactivation of X chromosomes in the germ line (Lee, 2005; Vibranovski, 2014). In *Timema,* we suggest an absence of dosage compensation is the most likely explanation as expression of X-linked genes in the male reproductive tracts show a major peak of genes with expression approximately half of that observed in females, similar to that observed in species that lack dosage compensation (Mank & Ellegren, 2009; Vicoso *et al*., 2013). It should be noted, however, that the expression reduction we see on the male X in *Timema* is actually slightly less than the half we would expect from a lack of dosage compensation alone. This suggests that other factors, described above, may also have an influence. Disentangling the contribution of each of these factors will require further work. Our work, however, clearly shows that expression in reproductive tissues behaves differently than in somatic tissues, highlighting the importance of studying these tissues separately particularly when assessing the extent of dosage compensation (Gu & Walters, 2017).

Dosage compensation in *Timema* somatic tissues appears to be achieved by the upregulation of the X in males (type I, (Gu & Walters, 2017)) as indicated by similar expression levels of genes on the X and the autosomes in both sexes. This interpretation assumes that the ancestral state of X chromosome expression (before it was a sex chromosome) was similar to the other autosomes. This seems likely as expression across all *Timema* autosomes is similar. However, assessing the ancestral state of X expression is largely impossible as the *Timema* X chromosome appears to be very old (~120 million years). As a consequence, there are likely no species available for comparison with conserved genome organisation but with X chromosomes that are homologous to different *Timema* autosomes. Similar to *Timema,* upregulation of the X chromosome in males also appears to be the mechanism for dosage compensation in all other XX/XY or XX/X0 insect systems yet studied, even when dosage compensation is incomplete (Gu & Walters, 2017). With our study, this amounts to information from seven different insect orders (Odonata (Chauhan *et al*., 2021), Phasmatodea (this study), Hemiptera (Pal & Vicoso, 2015; Richard *et al*., 2017), Orthoptera (Rayner *et al*., 2021), Strepsiptera (Mahajan & Bachtrog, 2015), Coleoptera (Prince *et al*., 2010; Mahajan & Bachtrog, 2015), and Diptera (Bone & Kuroda, 1996; Marín *et al*., 1996; Deng *et al*., 2011; Nozawa *et al.*, 2014; Jiang *et al.*, 2015; Vicoso & Bachtrog, 2015; Rose *et al.*, 2016)) suggesting that this mechanism may be universal for insect species with heterogametic males.

## Conclusions

How consistent the consequences of sex linkage are for gene sequence and expression evolution across taxa remains an open question. Here we examine several key aspects of sex chromosome evolution in a previously neglected group, phasmids. Overall, we find evidence for several predicted consequences, including complete dosage compensation of the X (in somatic tissues) and a faster rate of evolution of X-linked than autosomal genes. By contrast, we find little evidence that sex linkage facilitates the accumulation of sexually antagonistic alleles. While our results were consistent across different *Timema* species, they also show key differences from studies in other taxa, highlighting the importance of studying sex chromosome evolution across a diverse set of species to distinguish general patterns caused by sex linkage from species-specific idiosyncrasies.Data and code availability Raw sequence reads have been deposited in NCBI’s sequence read archive under the bioproject: PRJNA725673 (Table S8). Scripts for the analyses in this paper are available at: https://github.com/DarrenJParker/Timema_sex_chr_evol_code and will be archived at Zenodo upon acceptance. Data was processed to generate plots and statistics using R v4.0.3 and Python v.3.7.3 unless otherwise stated.

## Acknowledgements

We would like to thank Chloe Larose, Kirsten Jalvingh, and Bart Zijlstra for their assistance in the field. We thank Sébastien Moretti for his assistance with running Godon. We also thank Matthew Hahn for useful comments. This project was supported by several grants from the Swiss Science Foundation: PP00P3_170627 and 31003A_182495 to TS as well as CRSII3_160723 to TS, MRR and Nicolas Galtier. We would like to thank the former Vital-IT platform of the SIB Swiss Institute of Bioinformatics (SIB) and the DCSR for maintaining the computer infrastructure at the University of Lausanne.

## Notes

### Competing Interest Statement

The authors have declared no competing interest.

### Summary of Updates

R1.

