## Supplemental tables and figures for "X chromosomes show relaxed selection and complete somatic dosage compensation across *Timema* stick insect species"

This PDF file includes:

Figs. S1 to S46
Tables S1 to S9

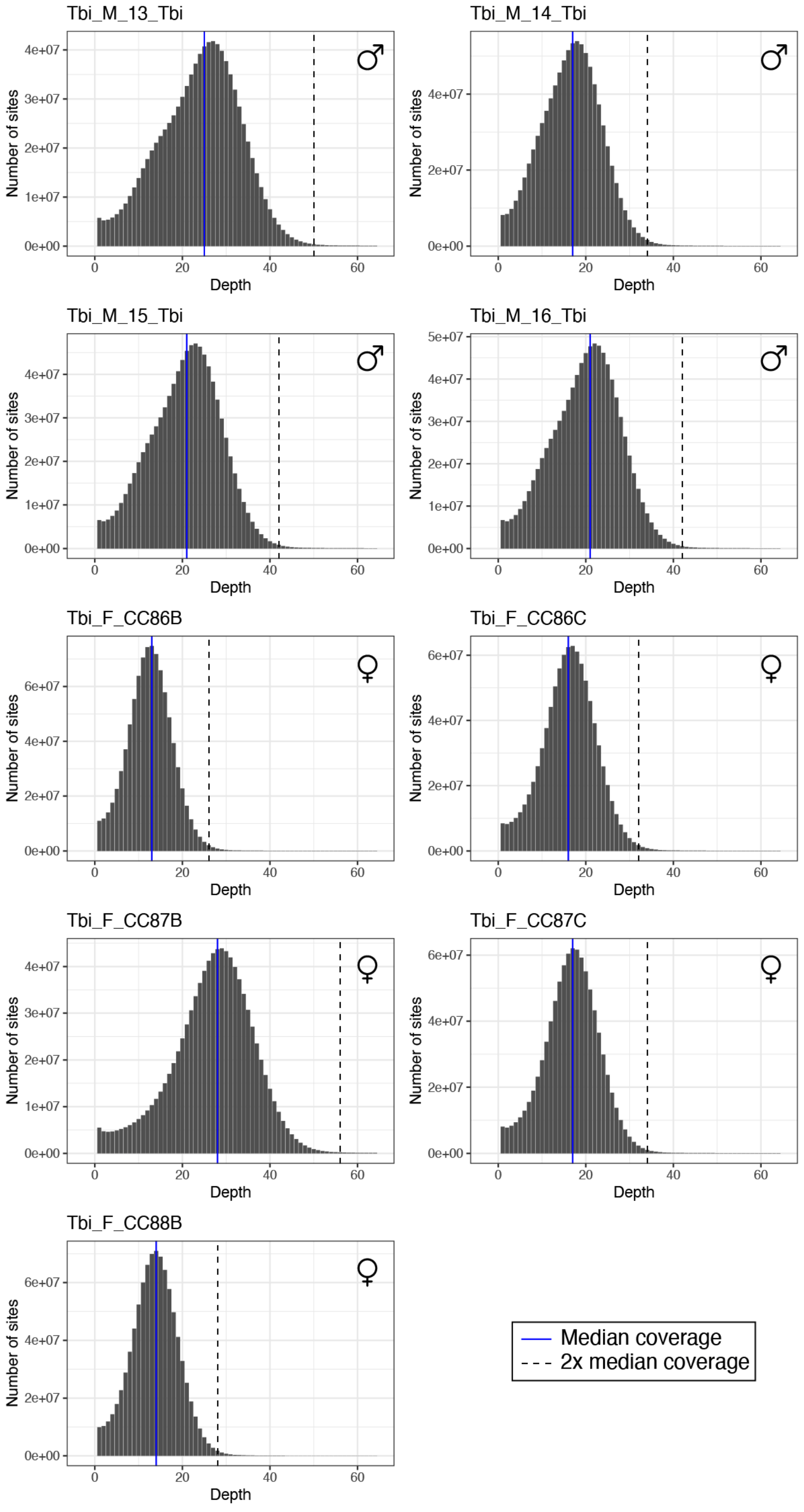


**Fig. S1 | Per base coverage in *T. bartmani.*** Blue solid lines represent the median coverage once sites with 0 coverage were excluded. Black dashed lines represent two-times the median coverage.

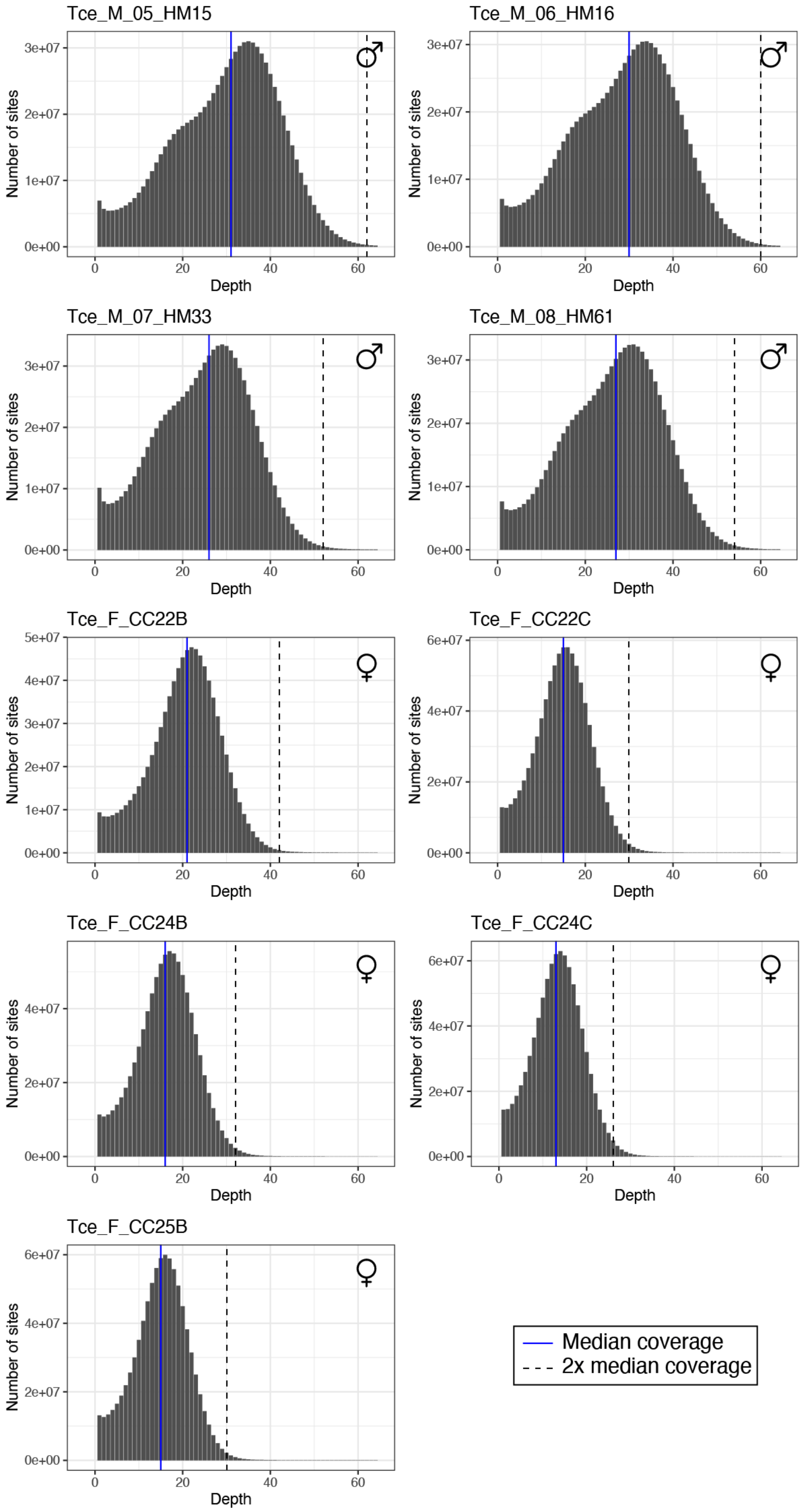


**Fig. S2 | Per base coverage in *T. cristinae.*** Blue solid lines represent the median coverage once sites with 0 coverage were excluded. Black dashed lines represent two-times the median coverage.

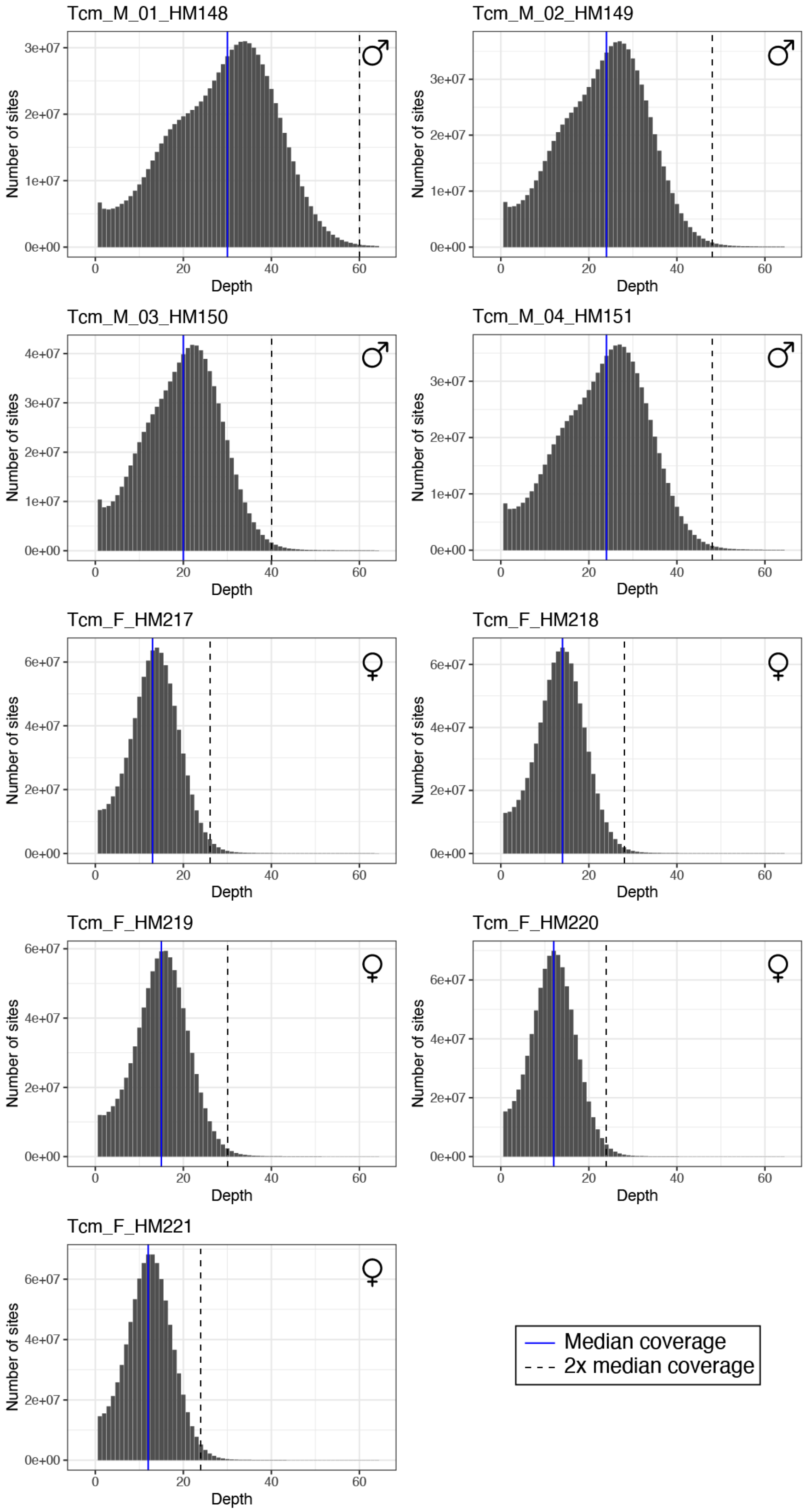


**Fig. S3 | Per base coverage in *T. californicum.*** Blue solid lines represent the median coverage once sites with 0 coverage were excluded. Black dashed lines represent two-times the median coverage.

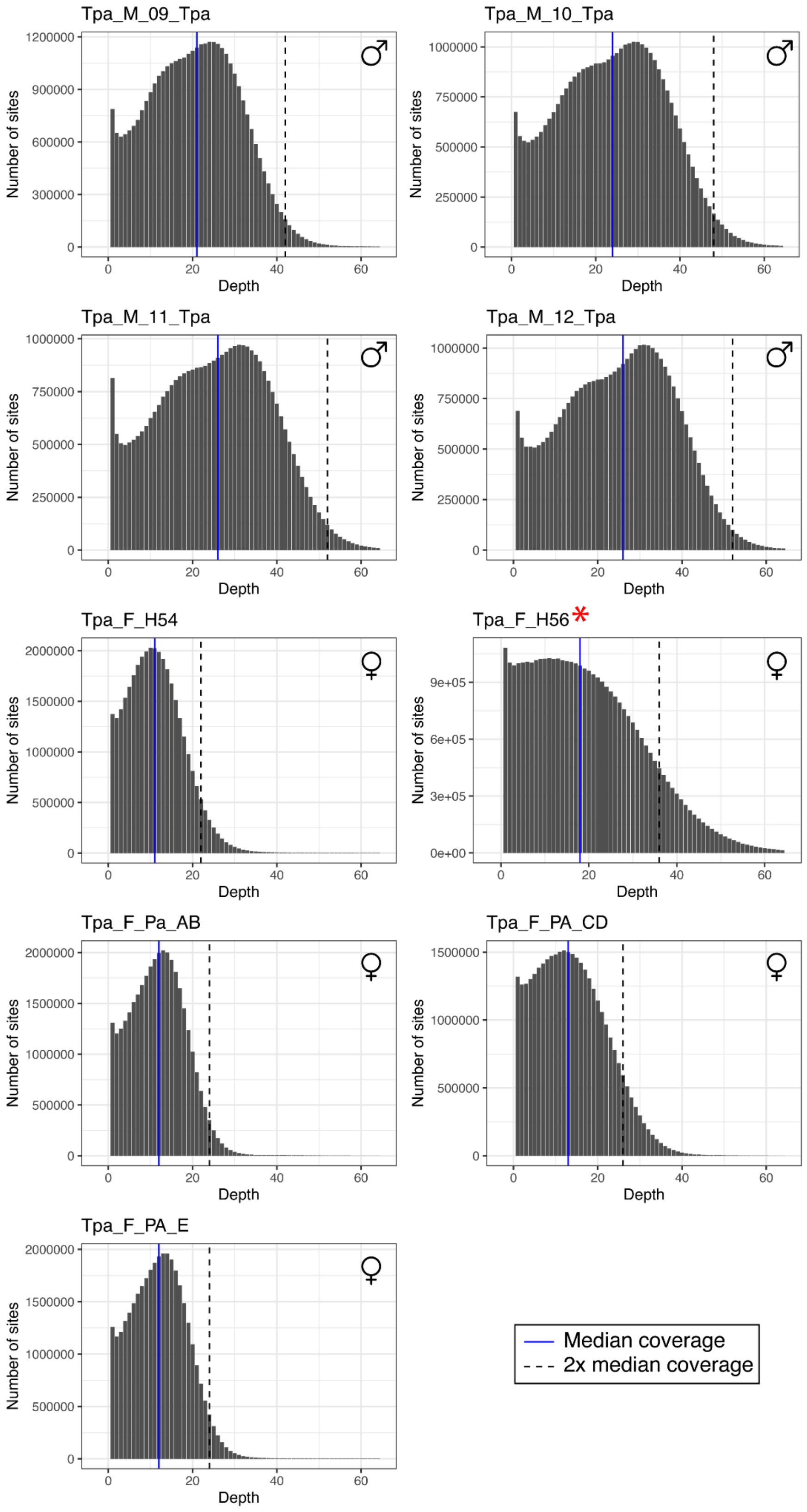


**Fig. S4 | Per base coverage in *T. podura.*** Blue solid lines represent the median coverage once sites with 0 coverage were excluded. Black dashed lines represent two-times the median coverage. Note sample H56 (indicated with a red asterisk) was excluded from all subsequent analyses.

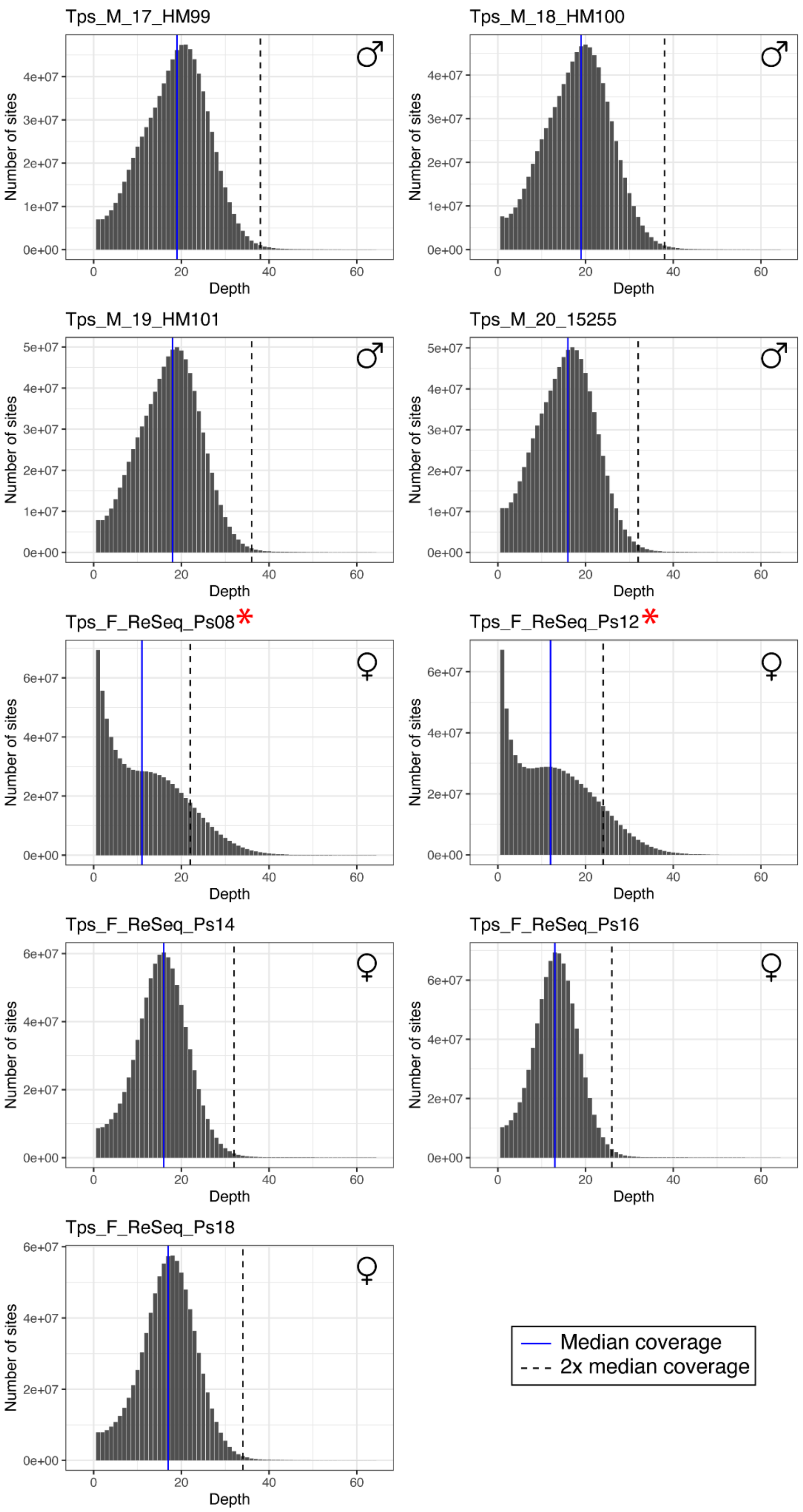


**Fig. S5 | Per base coverage in *T. poppensis*.** Blue solid lines represent the median coverage once sites with 0 coverage were excluded. Black dashed lines represent two-times the median coverage. Note samples Ps08 and Ps12 (indicated with red asterisks) were excluded from all subsequent analyses.

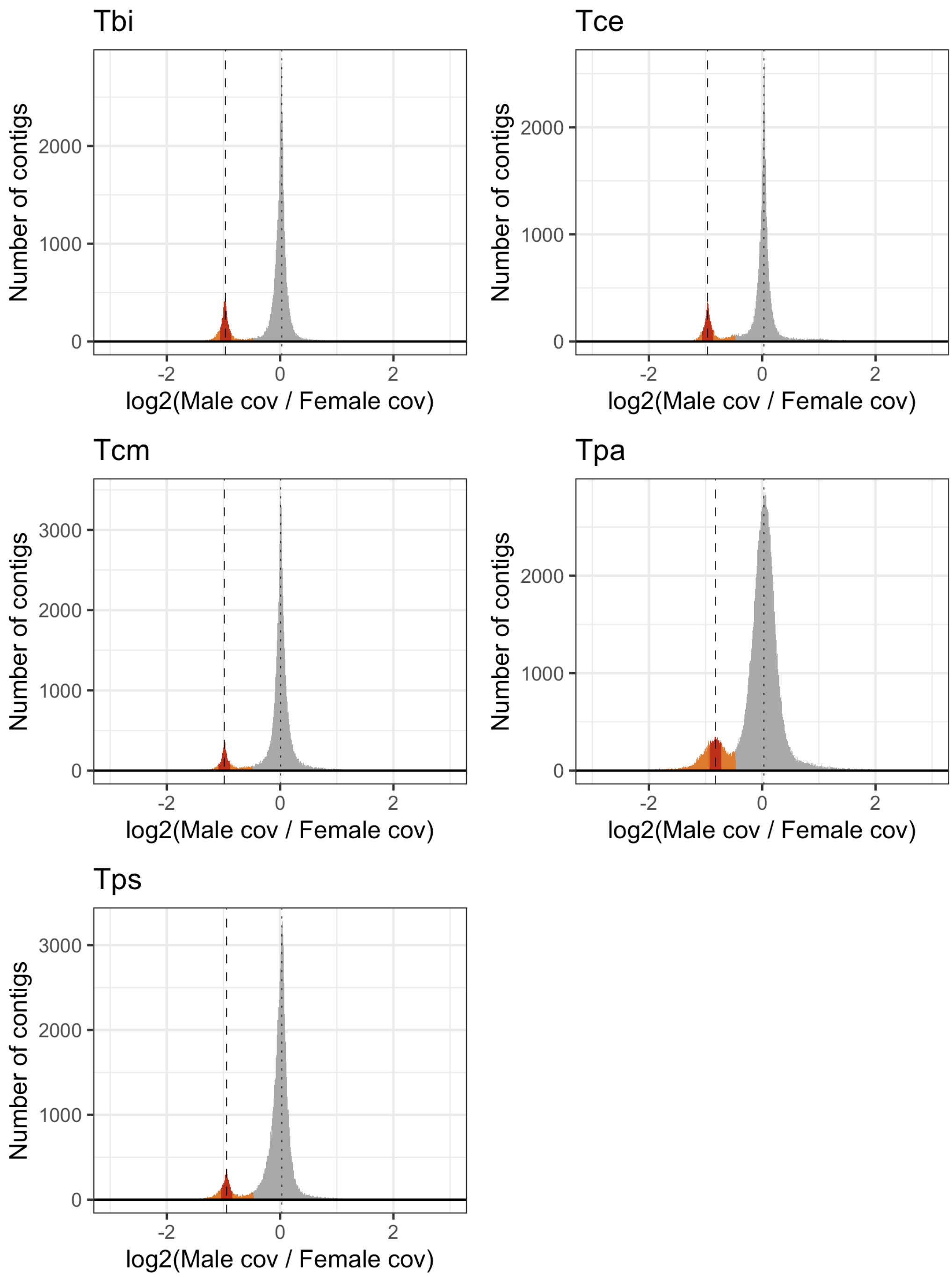


**Fig. S6 | Frequency distribution of $log_2$ ratio of male to female coverage for all contigs >= 1000 bp.** Autosomal contigs (grey) should have equal coverage in males and females ($log_2$ ratio of male to female coverage = 0). X-linked contigs should have half the coverage in males than in females ($log_2$ ratio of male to female ratio coverage = -1). Dotted lines indicate the distribution peaks. X-linked scaffolds were classed in two ways; Liberal: contigs with a $log_2$ ratio of male to female coverage < Autosomal peak - 0.5 (orange and red) and Stringent: contigs with a $log_2$ ratio of male to female coverage within 0.1 of the X linked peak (red). Species names are abbreviated as Tbi = *T. bartmani*, Tce = *T. cristinae*, Tcm = *T. californicum*, Tps = *T. poppensis*, and Tpa = *T. podura*.

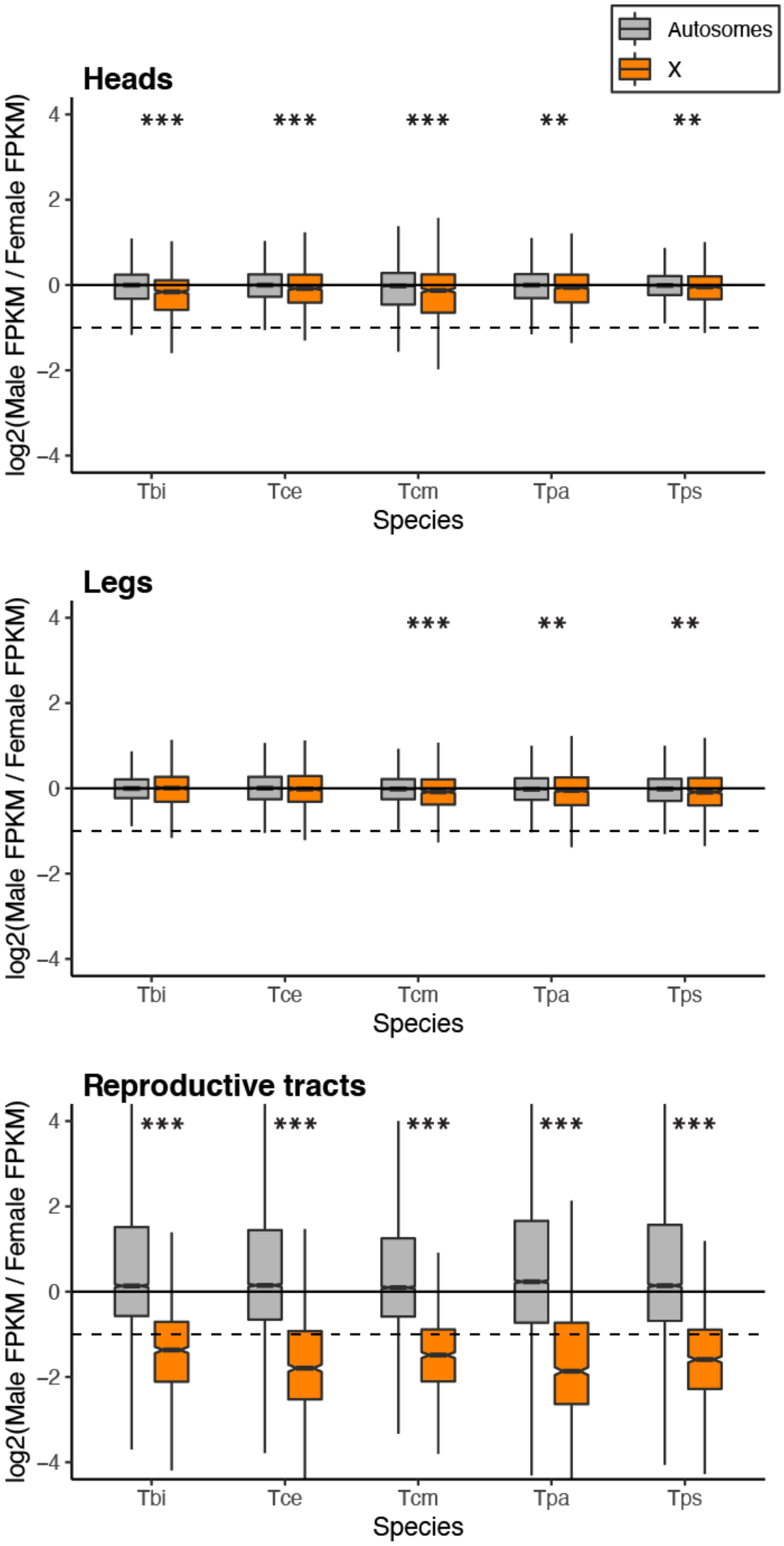


**Fig. S7 | Log2 ratio of male to female expression for the X and autosomes when sex-specific genes are retained.** Dashed line represents a two-fold reduction in expression in males (as expected if there was no dosage compensation). Species names are abbreviated as Tbi = *T. bartmani*, Tce = *T. cristinae*, Tcm = *T. californicum*, Tps = *T. poppensis*, and Tpa = *T. podura*.

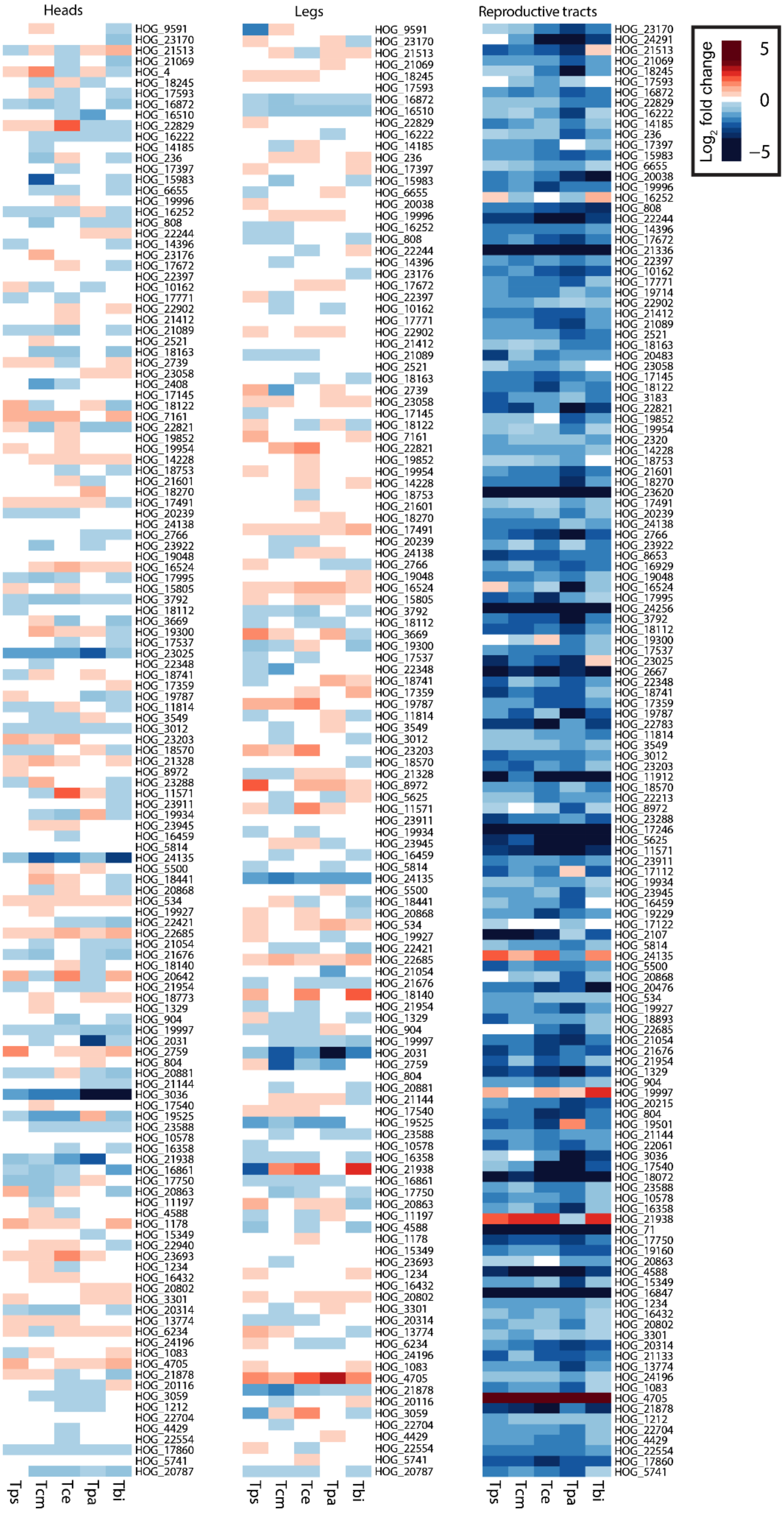


**Fig. S8 | Heatmap of gene expression of orthologs on the X in the heads, legs and reproductive tracts when sex-specific genes are retained.** Species names are abbreviated as Tbi = *T. bartmani*, Tce = *T. cristinae*, Tcm = *T. californicum*, Tps = *T. poppensis*, and Tpa = *T. podura*.

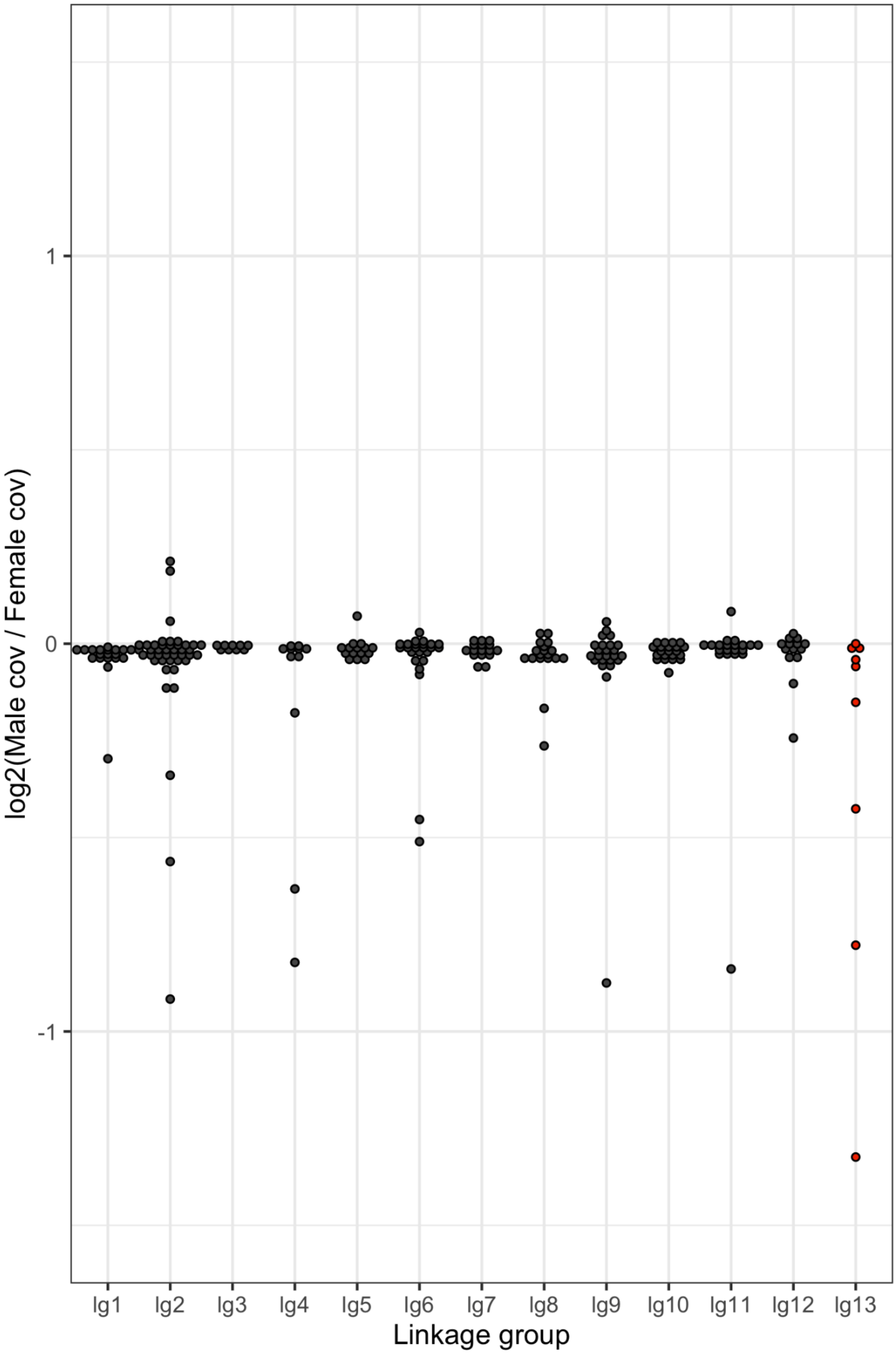


**Fig. S9 | Log$_2$ ratio of male to female coverage of scaffolds assigned to linkage groups in** (Nosil *et al.*, 2018)**.** The hypothesized X chromosome, LG13, is shown in red.

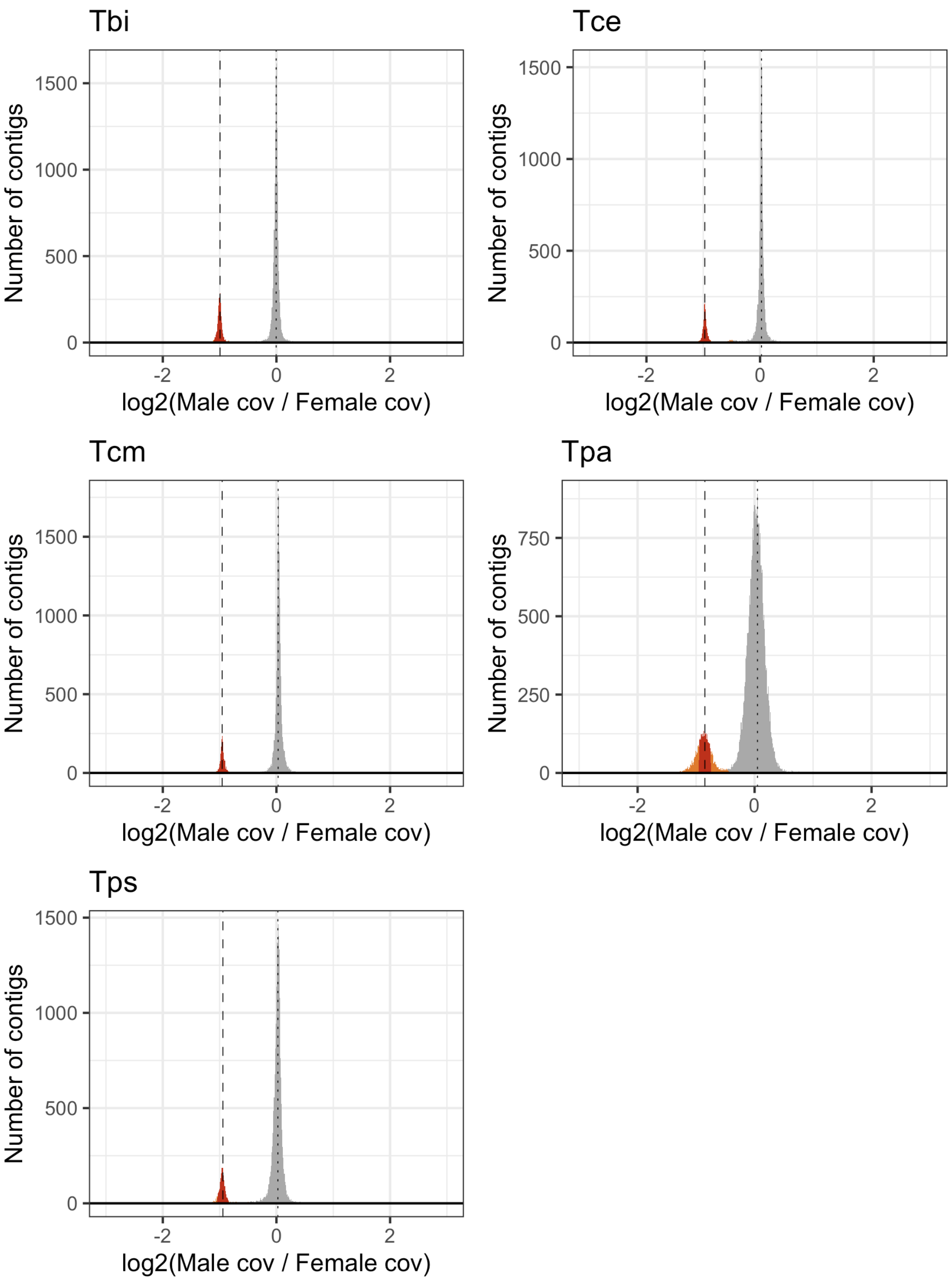


**Fig. S10 | Frequency distribution of $log_2$ ratio of male to female coverage for all contigs >= 5000 bp in length.** Autosomal contigs (grey) should have equal coverage in males and females ($log_2$ ratio of male to female coverage = 0). X-linked contigs should have half the coverage in males than in females ($log_2$ ratio of male to female ratio coverage = -1). Dotted lines indicate the distribution peaks. X-linked scaffolds were classed in two ways; Liberal: contigs with a $log_2$ ratio of male to female coverage < Autosomal peak - 0.5 (orange and red) and Stringent: contigs with a $log_2$ ratio of male to female coverage within 0.1 of the X linked peak (red). Species names are abbreviated as Tbi = *T. bartmani*, Tce = *T. cristinae*, Tcm = *T. californicum*, Tps = *T. poppensis*, and Tpa = *T. podura*.

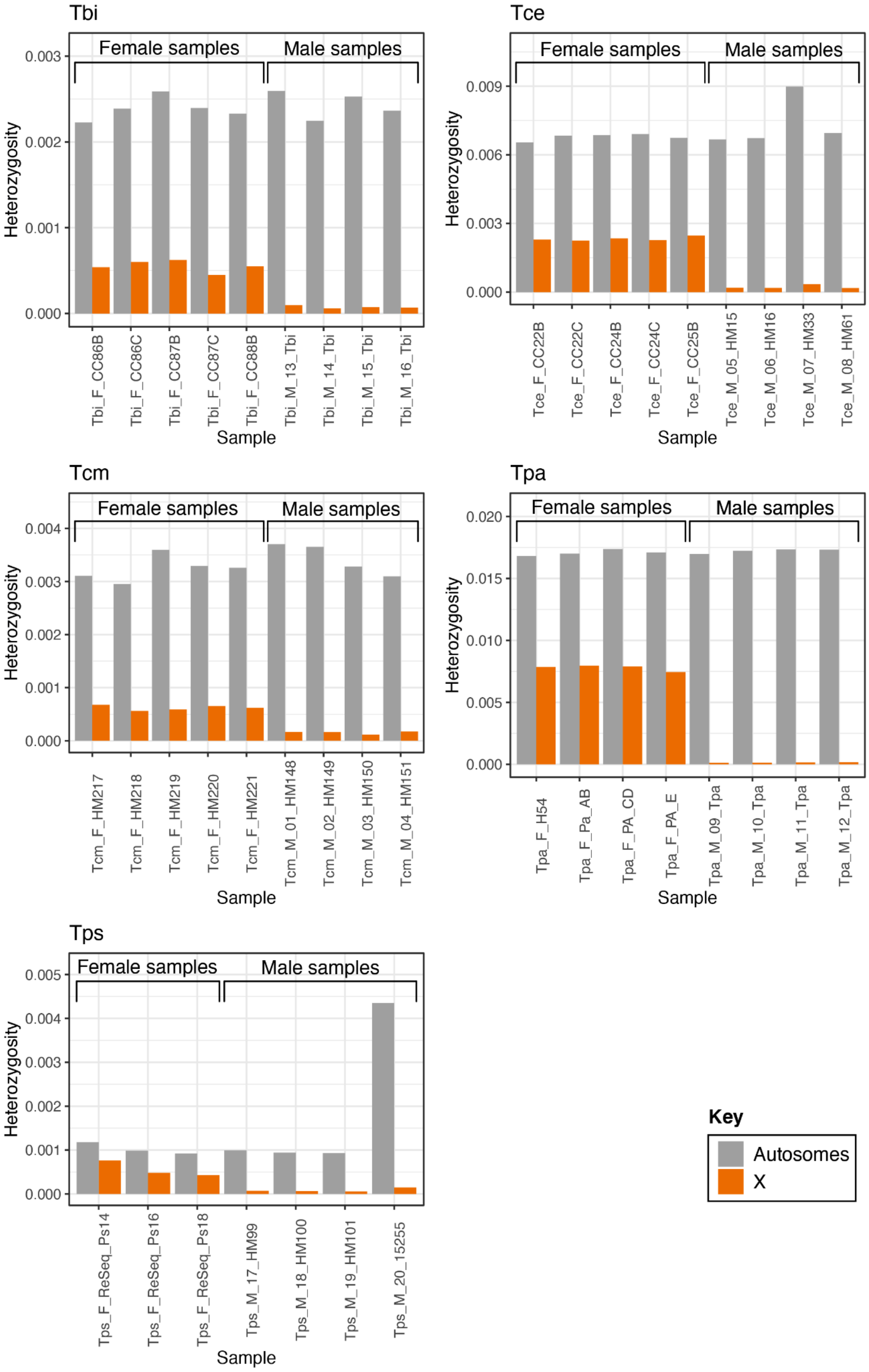


**Fig. S11 | Heterozygosity on the X and autosomes per sample.** Heterozygosity values displayed are the median heterozygosity of scaffolds weighted by scaffold length. Species names are abbreviated as Tbi = *T. bartmani*, Tce = *T. cristinae*, Tcm = *T. californicum*, Tps = *T. poppensis*, and Tpa = *T. podura*.

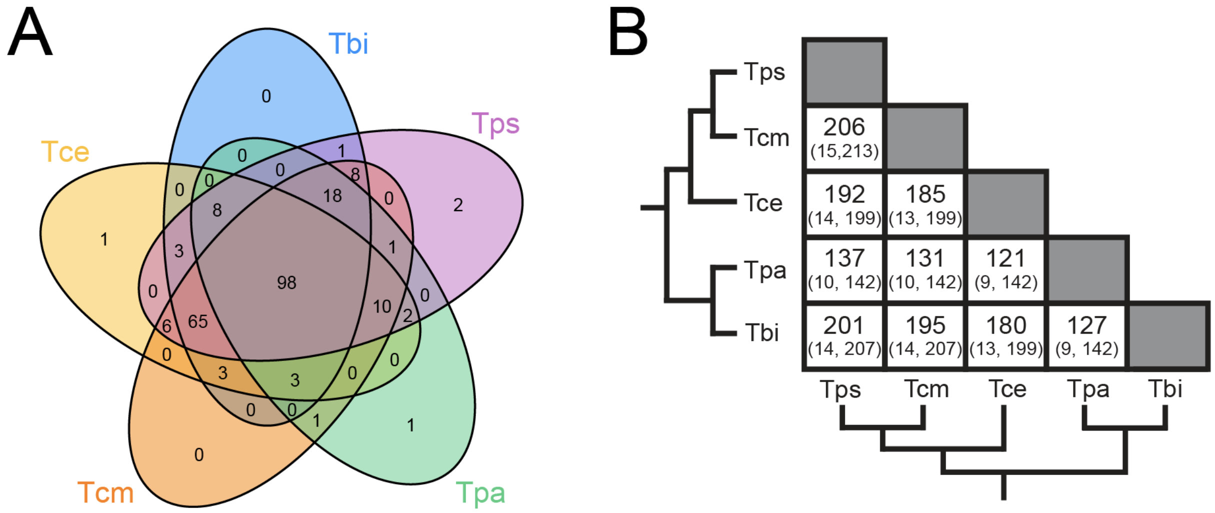


**Fig. S12 |** A) Venn-diagrams showing the number of shared X-linked orthologs between species when using the stringent coverage classification. B) Number of shared orthologs (expected, maximum possible) when using the stringent classification. The observed amount of overlap was much greater than expected in all comparisons (FDR < $3.425 \times 10^{-314}$). Species names are abbreviated as Tbi = *T. bartmani*, Tce = *T. cristinae*, Tcm = *T. californicum*, Tps = *T. poppensis*, and Tpa = *T. podura*.

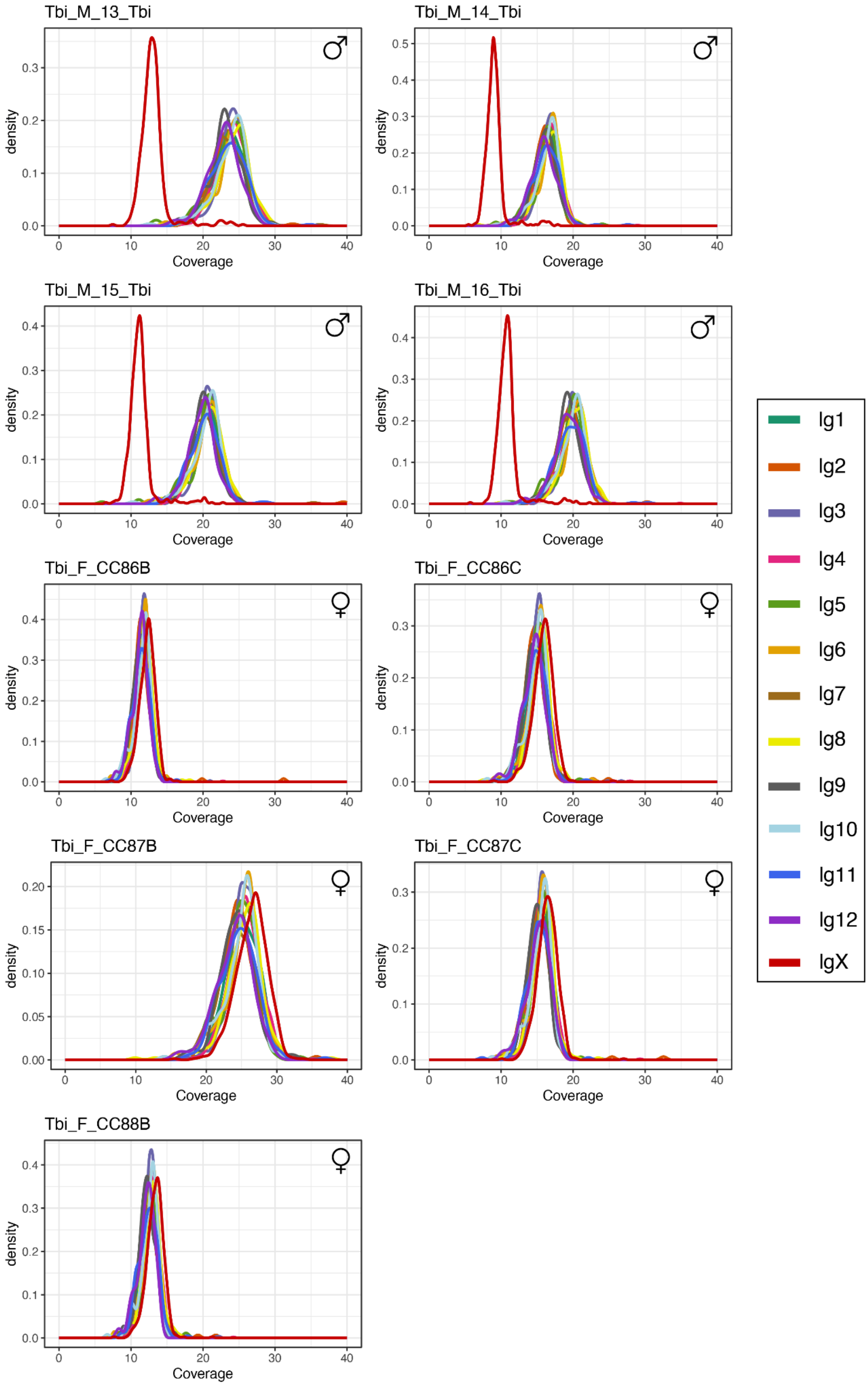


**Fig. S13 | Coverage of contigs assigned to linkage groups in *T. bartmani*.**

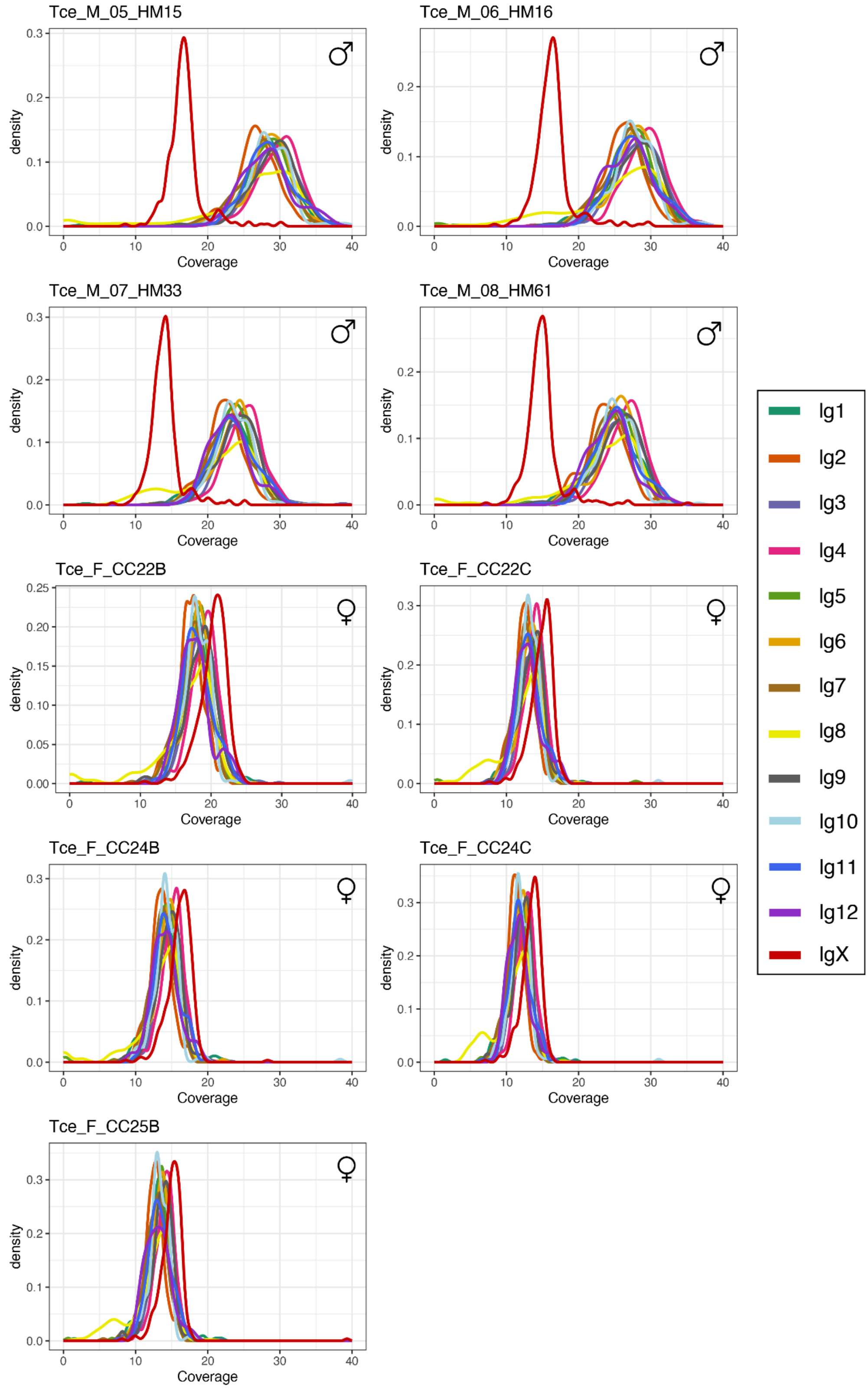


**Fig. S14 | Coverage of contigs assigned to linkage groups in *T. cristinae.***

107

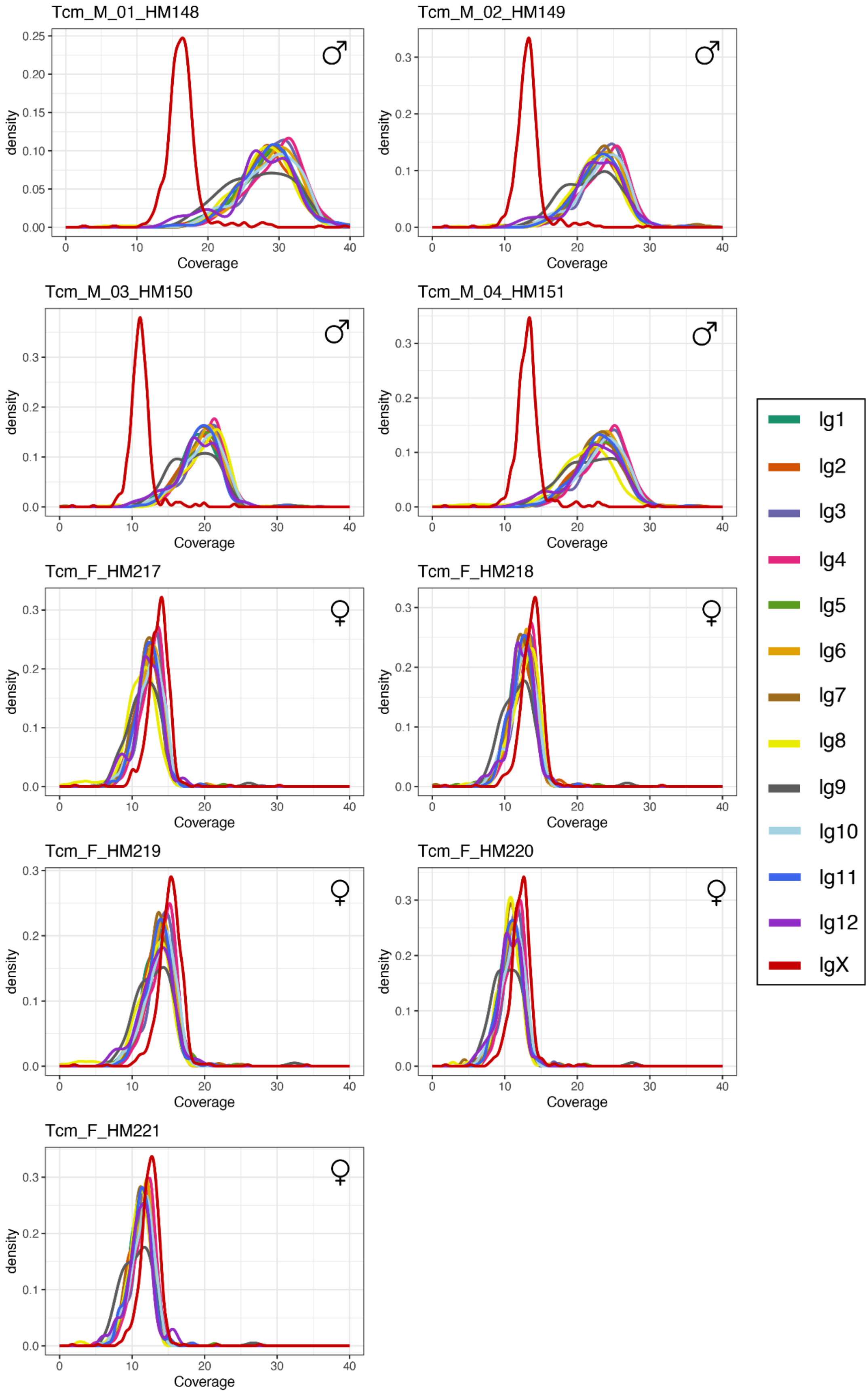


108 **Fig. S15 | Coverage of contigs assigned to linkage groups in *T. californicum.***

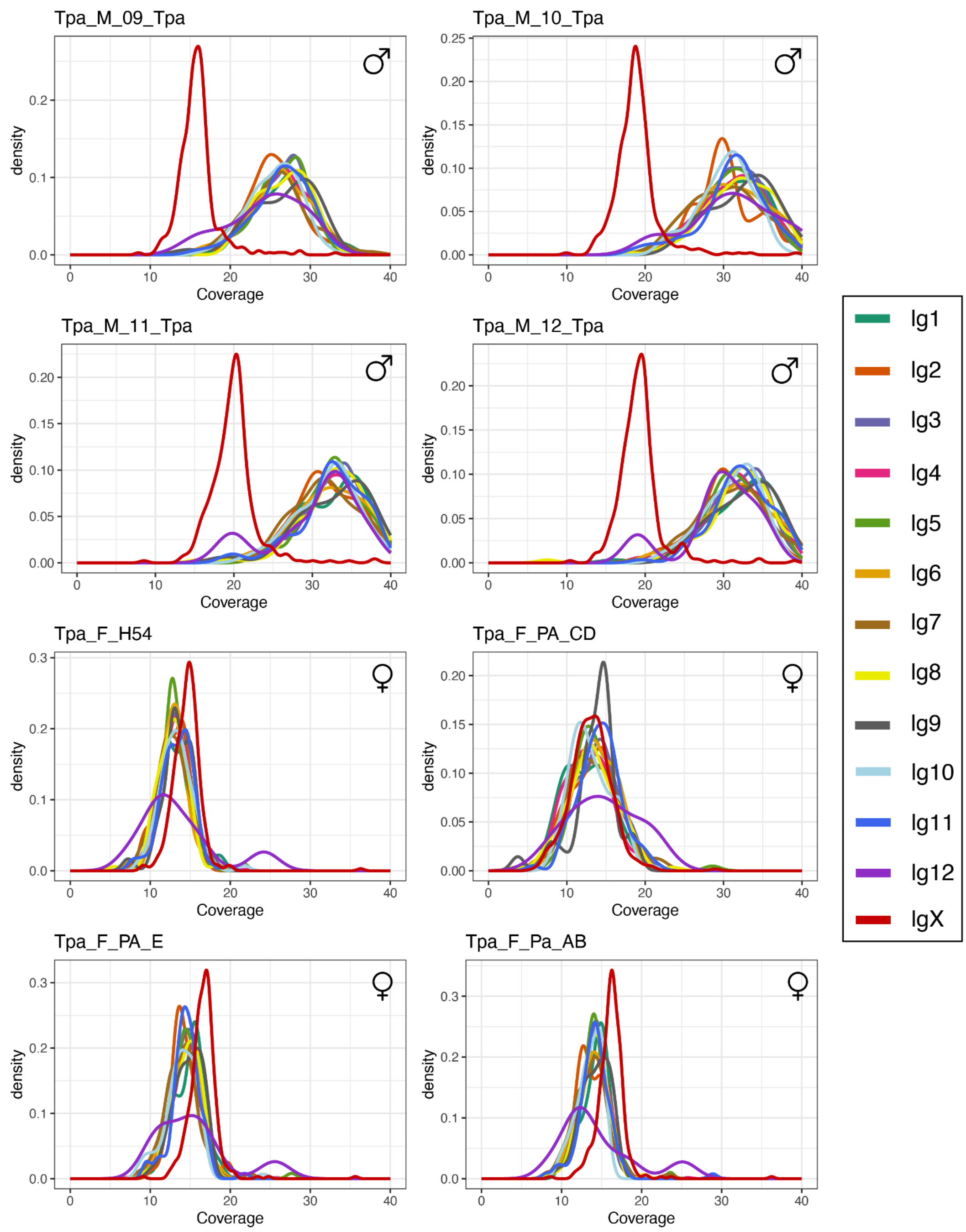


**Fig. S16 | Coverage of contigs assigned to linkage groups in *T. podura.***

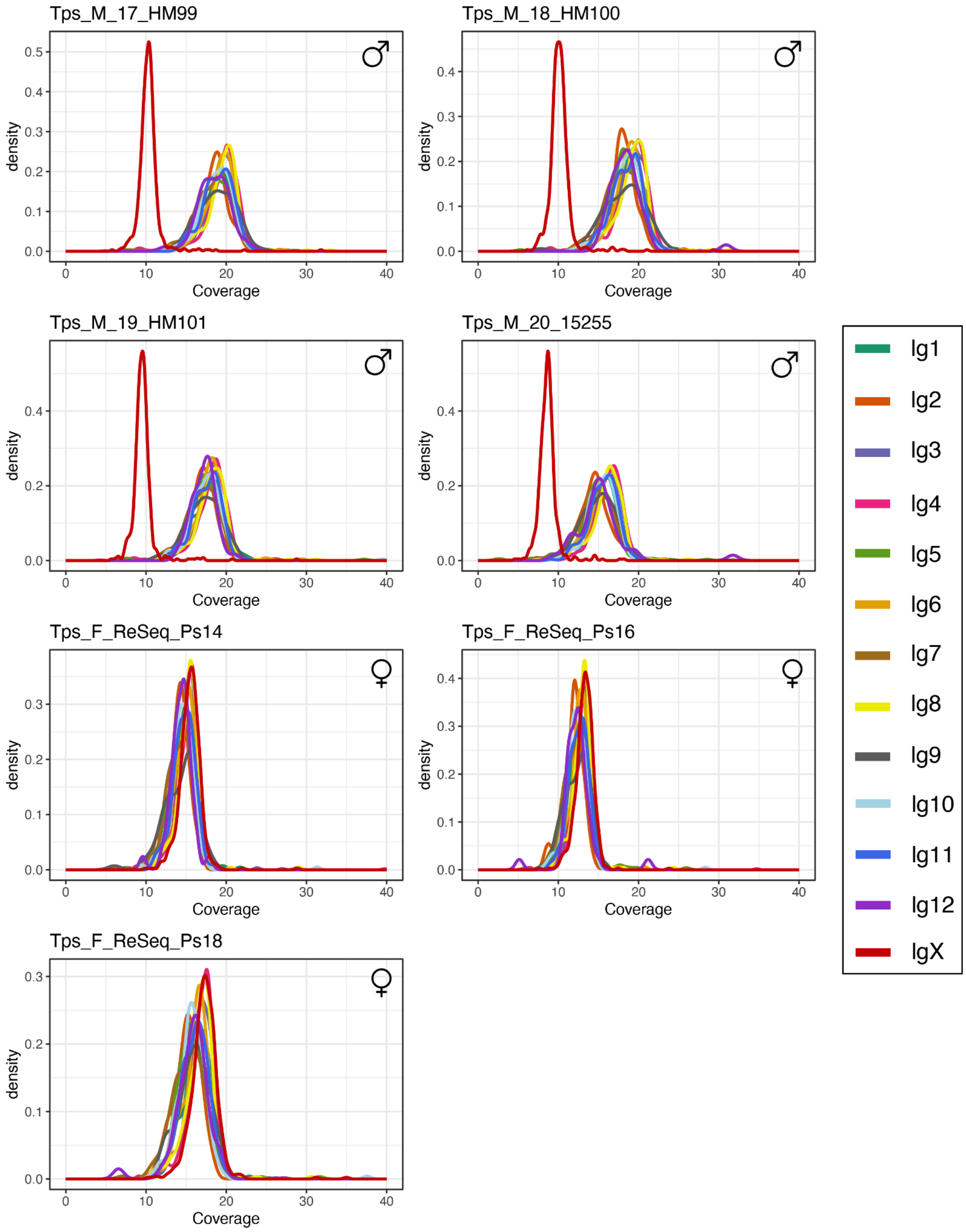


**Fig. S17 | Coverage of contigs assigned to linkage groups in *T. poppensis.***

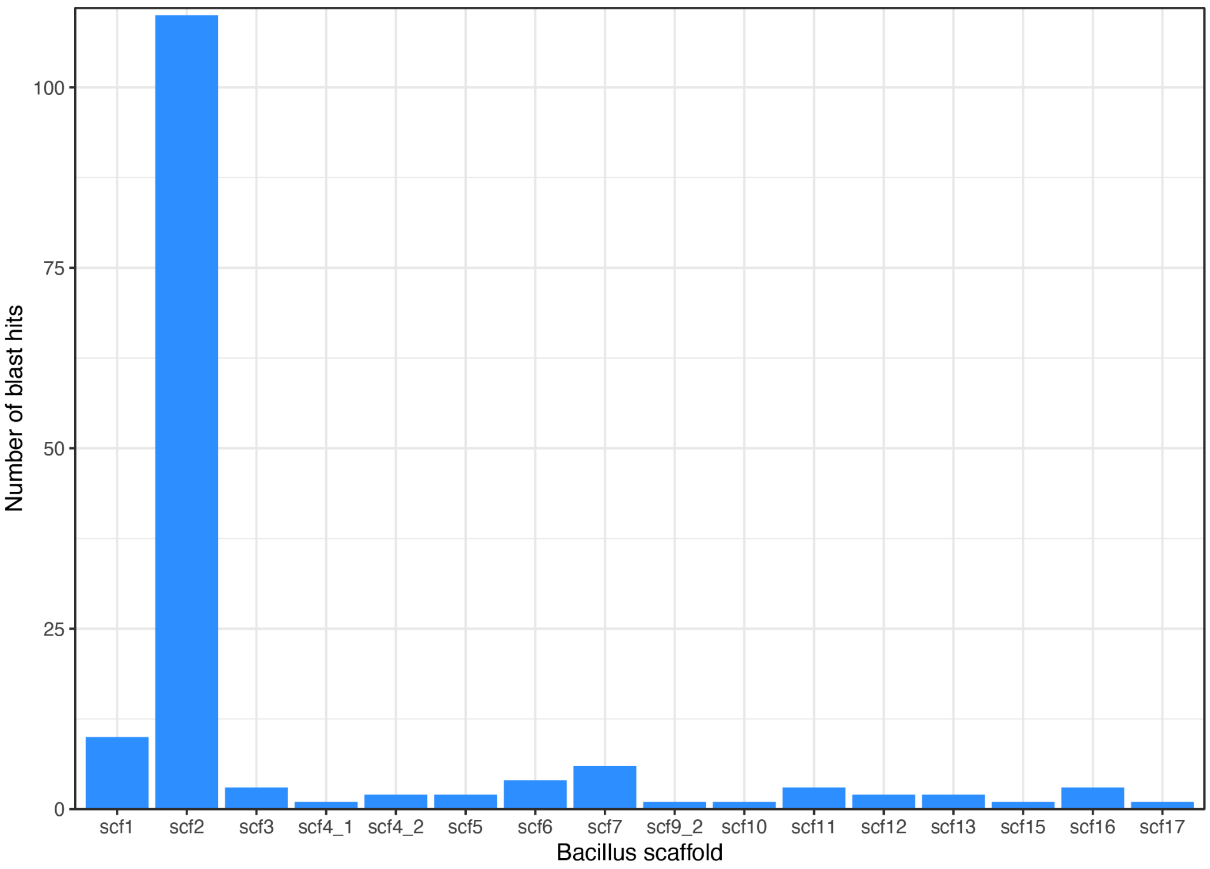


**Fig. S18 | Number of blast hits of *Timema* X orthologs to *Bacillus rossius* scaffolds. Scaffold 2 represents the X chromosome in this species.**

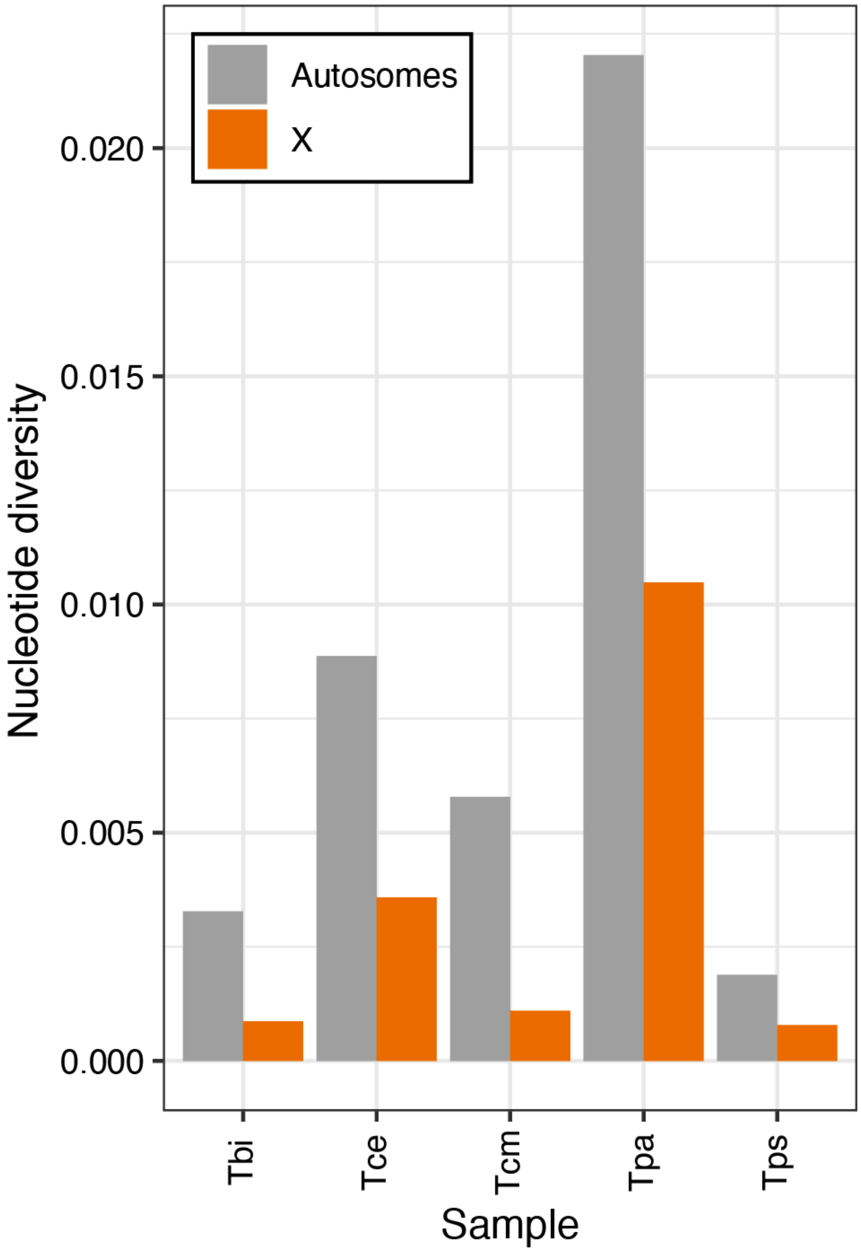


**Fig. S19 | Pairwise nucleotide diversity (π) on the X and autosomes.** Values displayed are the median nucleotide diversity of scaffolds weighted by scaffold length. Species names are abbreviated as Tbi = *T. bartmani*, Tce = *T. cristinae*, Tcm = *T. californicum*, Tps = *T. poppensis*, and Tpa = *T. podura*.

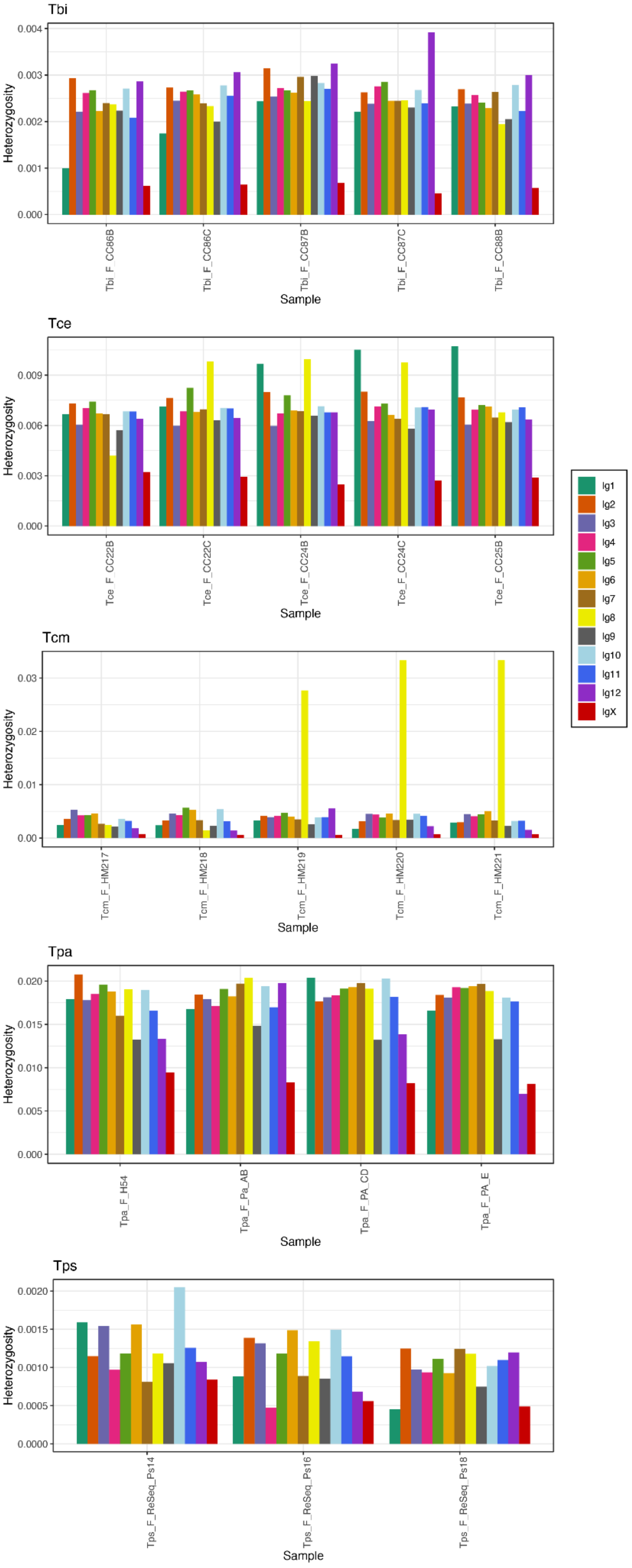


**Fig. S20 | Heterozygosity per linkage group per sample.** Heterozygosity values displayed are the median heterozygosity of scaffolds weighted by scaffold length. Note the high heterozygosity for some individuals on linkage group 8 in *T. californicum* is likely due to these individuals being heterozygous for different haplotypes associated with differences in colouration (Nosil *et al.*, 2018; Jaron *et al.*, 2022). Species names are abbreviated as Tbi = *T. bartmani*, Tce = *T. cristinae*, Tcm = *T. californicum*, Tps = *T. poppensis*, and Tpa = *T. podura*.

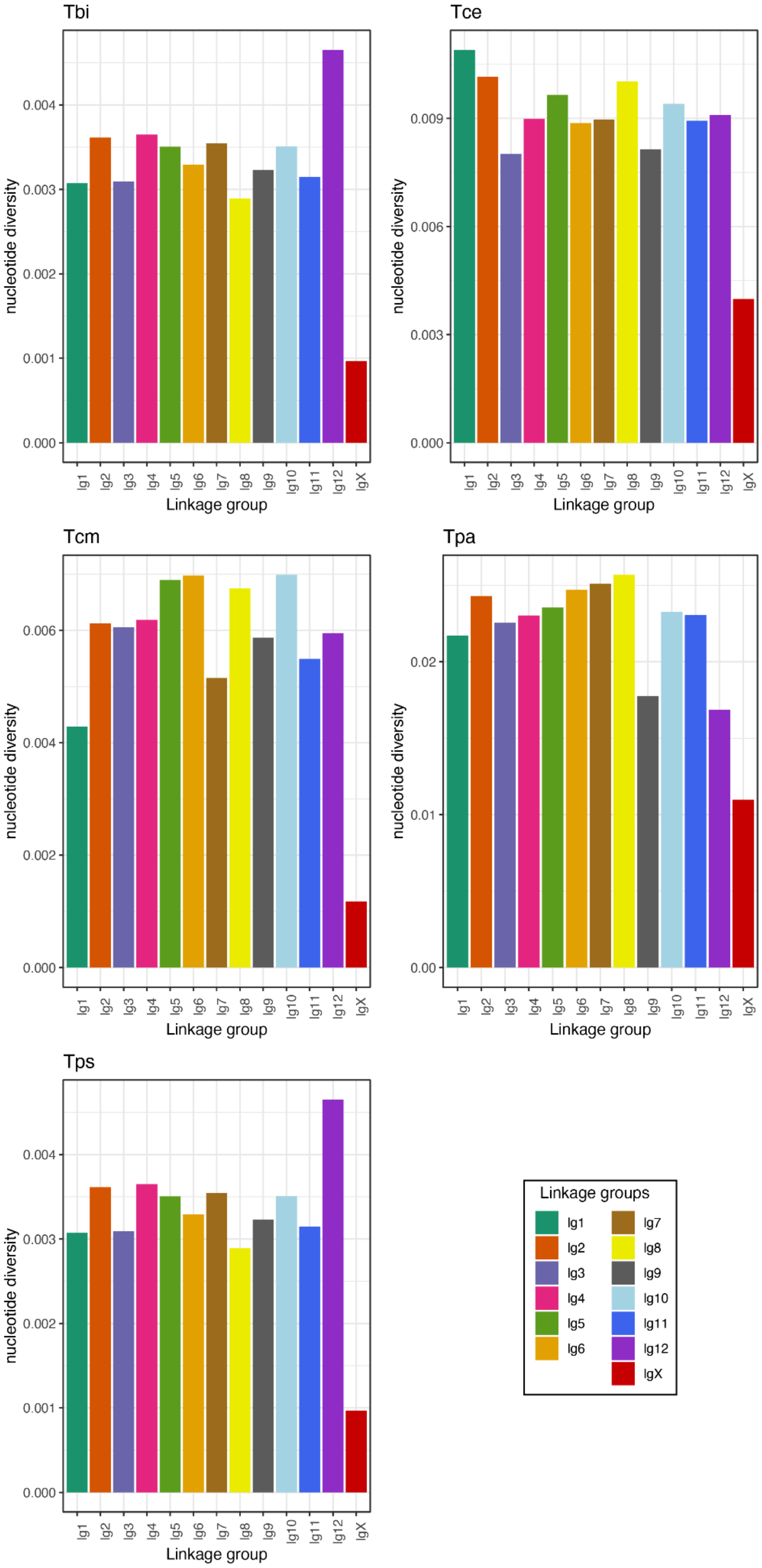


**Fig. S21 | Pairwise nucleotide diversity (π) on each linkage group.** Values displayed are the median nucleotide diversity of scaffolds weighted by scaffold length. Species names are abbreviated as Tbi = *T. bartmani*, Tce = *T. cristinae*, Tcm = *T. californicum*, Tps = *T. poppensis*, and Tpa = *T. podura*.

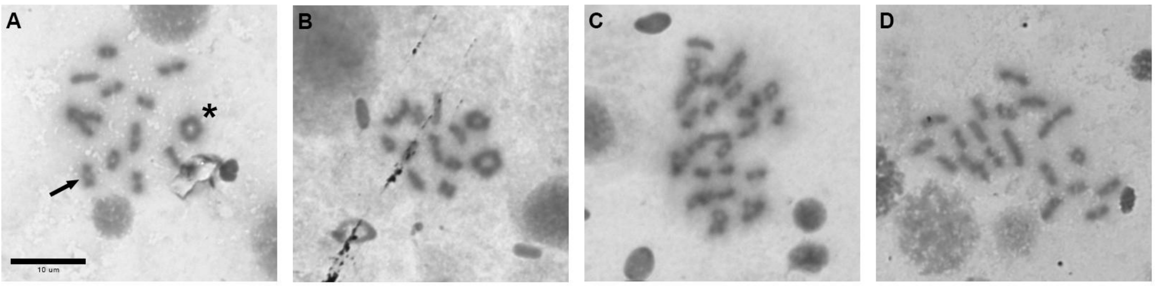


**Fig. S22 | Chiasmate meiosis in *Timema* males.** Most chromosomes are acrocentric and with a single, distally positioned chiasma (as in the chromosome pair highlighted by an arrow in panel A). Few chromosome pairs have two chiasmata, which are generally located more near to the centromere and telomere (star in panel A). **A**. *T. poppensis,* **B**. *T. californicum,* **C**. *T. cristinae,* **D**. *T. podura.*

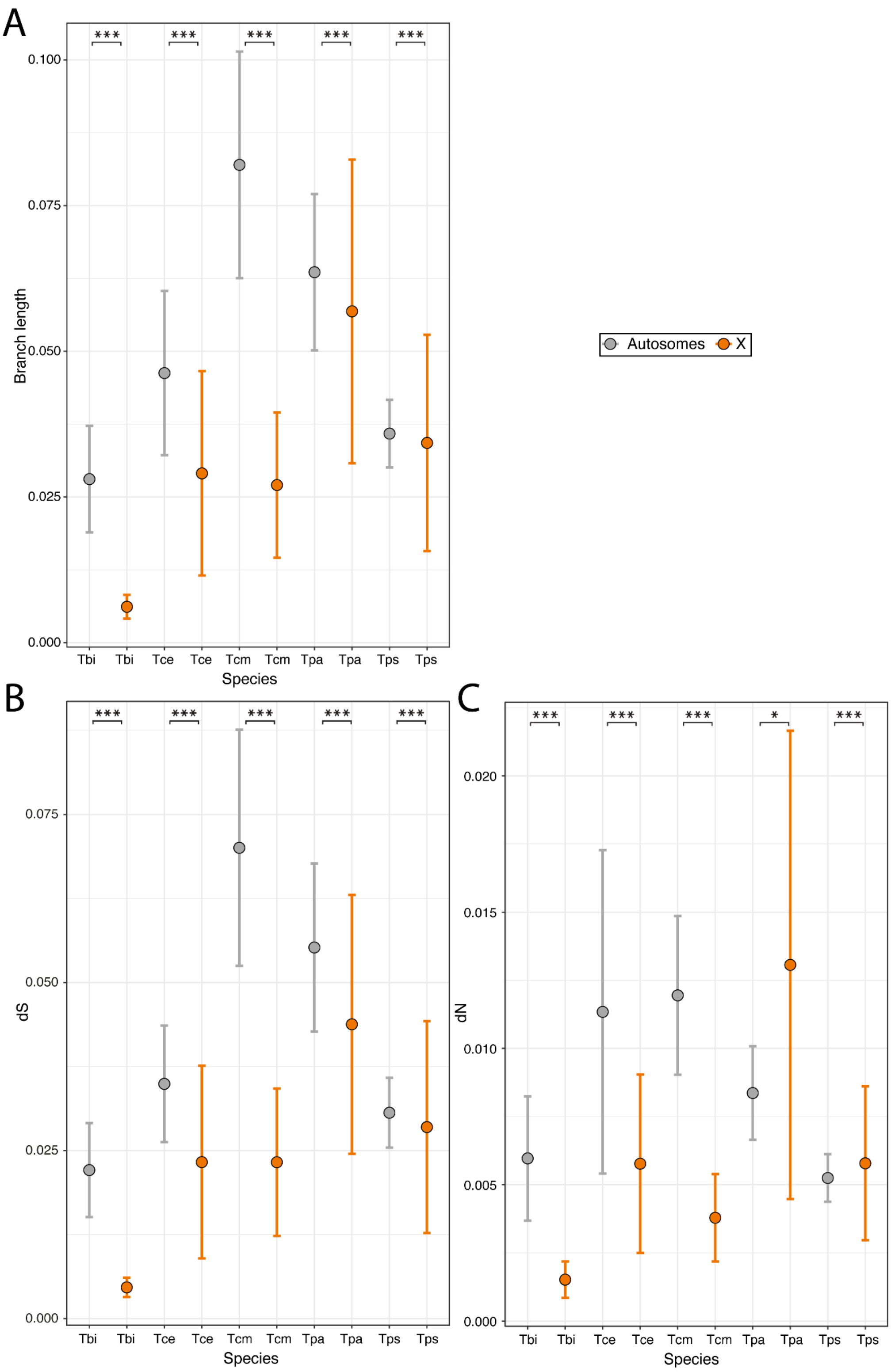


**Fig. S23 | Sequence evolution on the X and autosomes. A.** Average branch lengths across genes. **B.** Average dS values across genes**. C.** Average dN values across genes**.** Error bars indicate standard error. Asterisks indicate the significance (FDR) of Wilcoxon tests (***<0.001, **<0.01, *<0.05).

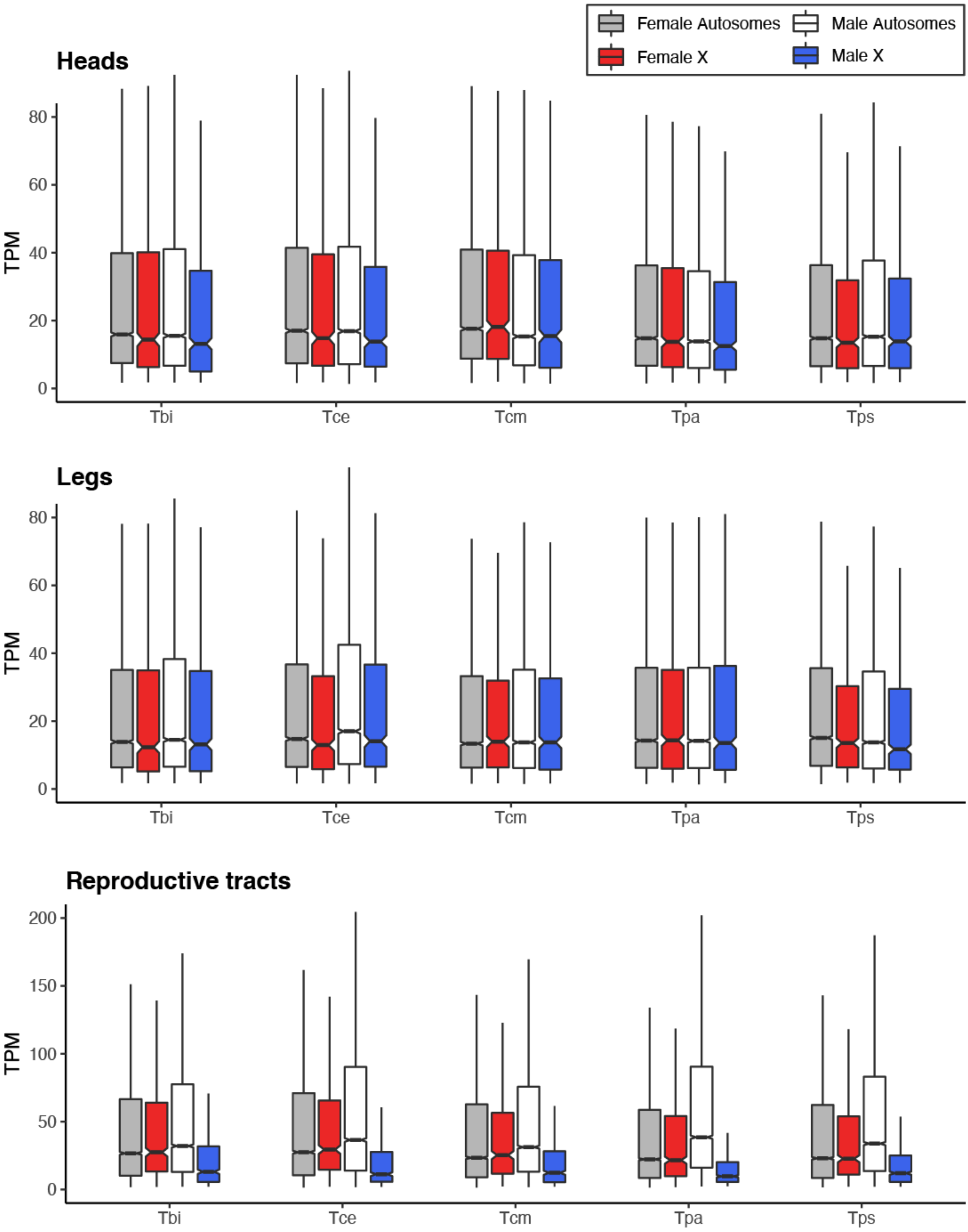


**Fig. S24 | Average expression levels in males and females on the X and autosomes using TPM.** Species names are abbreviated as Tbi = *T. bartmani*, Tce = *T. cristinae*, Tcm = *T. californicum*, Tps = *T. poppensis*, and Tpa = *T. podura*.

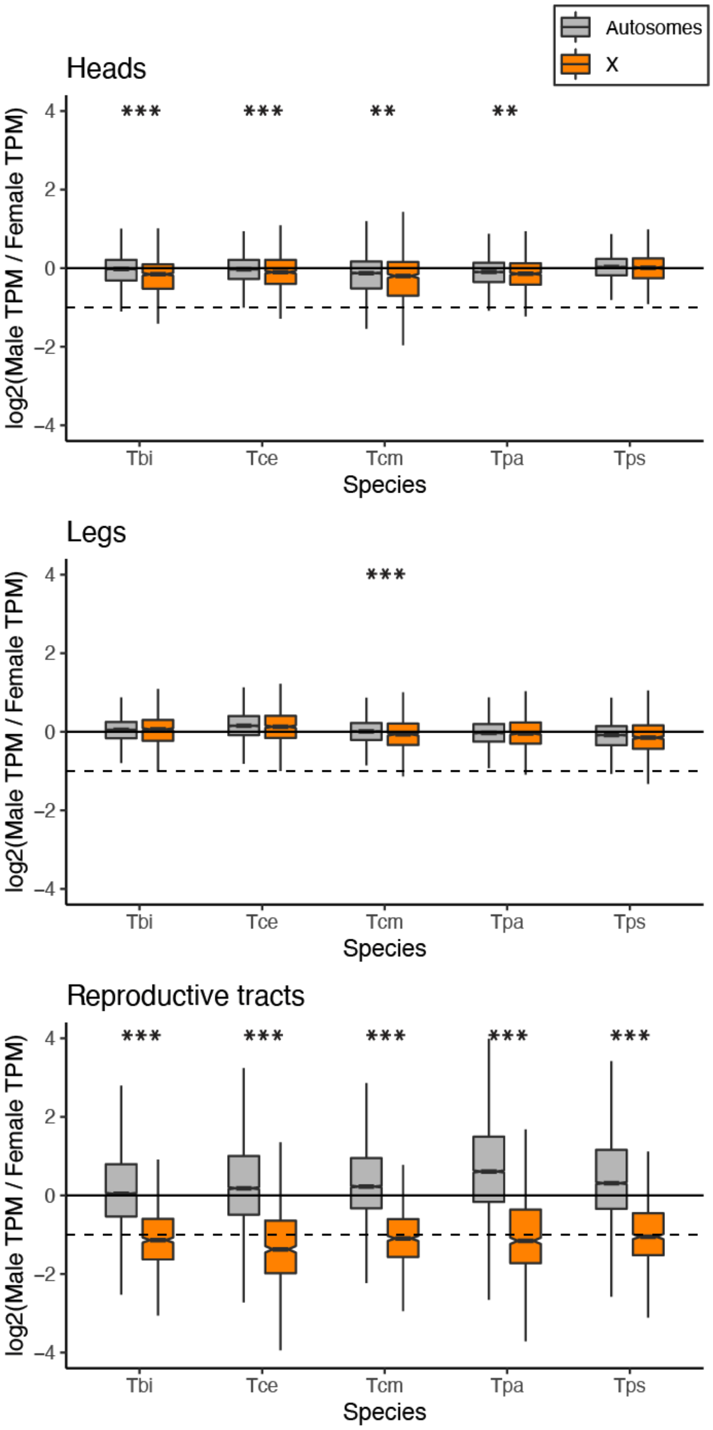


**Fig. S25 | Log$_2$ of male to female ratio of expression (TPM) for the X and autosomes.** Dashed line represents a two-fold reduction in expression in males (as expected if there was no dosage compensation). Species names are abbreviated as Tbi = *T. bartmani*, Tce = *T. cristinae*, Tcm = *T. californicum*, Tps = *T. poppensis*, and Tpa = *T. podura*.

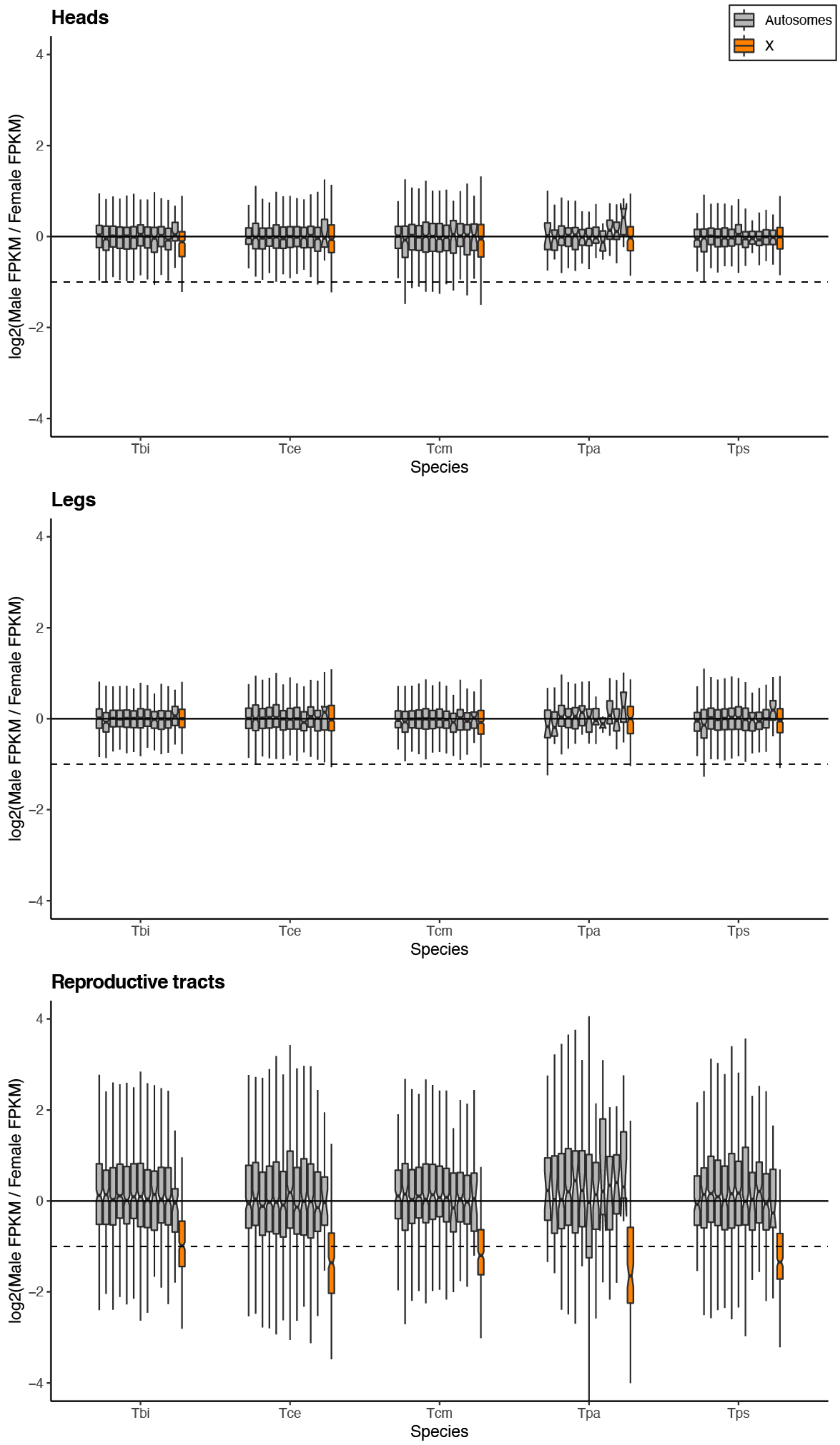


**Fig. S26 | Log2 of male to female ratio of expression (FPKM) for different linkage groups.** Dashed line represents a two-fold reduction in expression in males (as expected if there was no dosage compensation). Species names are abbreviated as Tbi = *T. bartmani*, Tce = *T. cristinae*, Tcm = *T. californicum*, Tps = *T. poppensis*, and Tpa = *T. podura*.

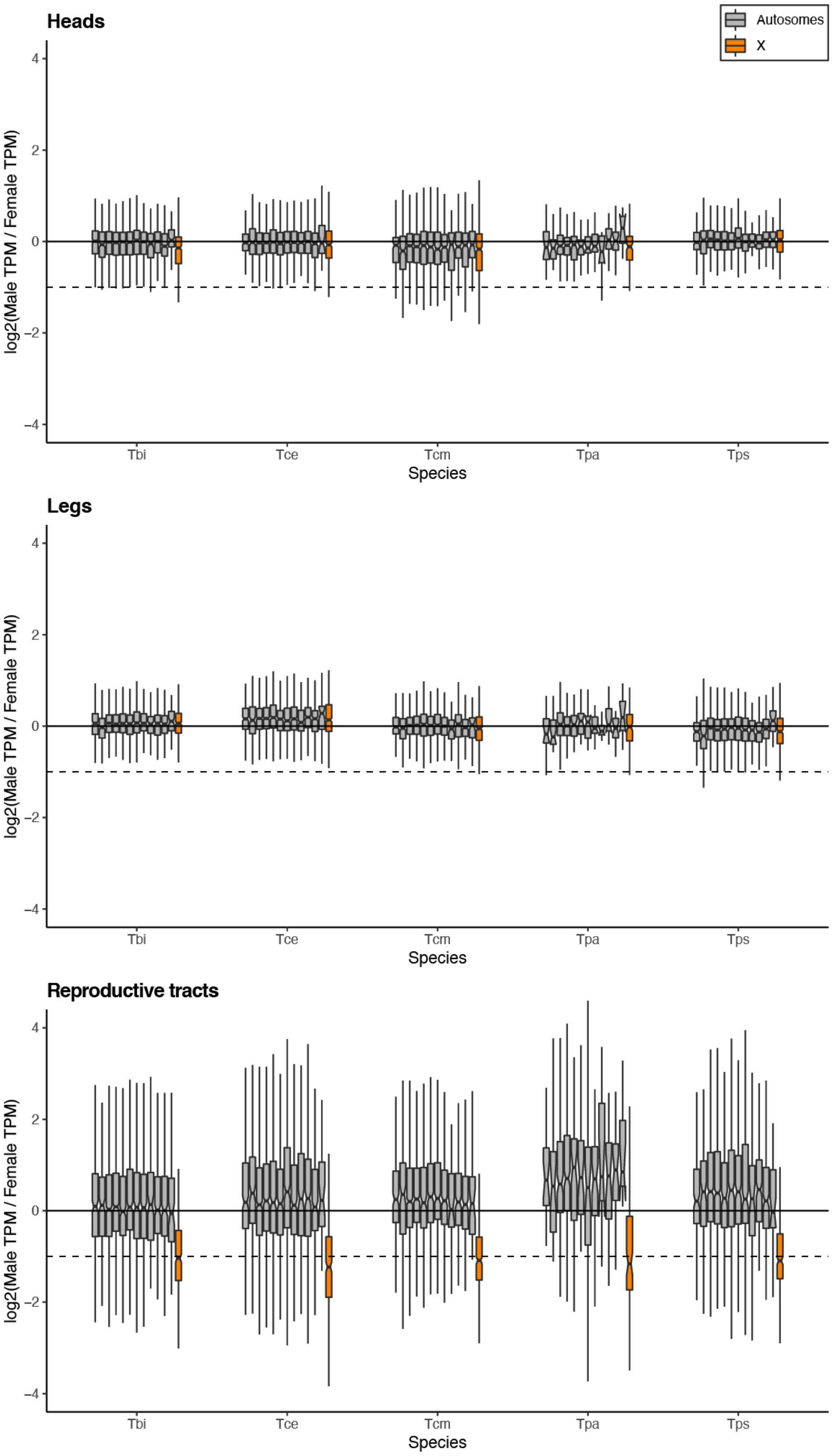


**Fig. S27 | Log2 of male to female ratio of expression (TPM) for different linkage groups.** Dashed line represents a two-fold reduction in expression in males (as expected if there was no dosage compensation). Species names are abbreviated as Tbi = *T. bartmani*, Tce = *T. cristinae*, Tcm = *T. californicum*, Tps = *T. poppensis*, and Tpa = *T. podura*.

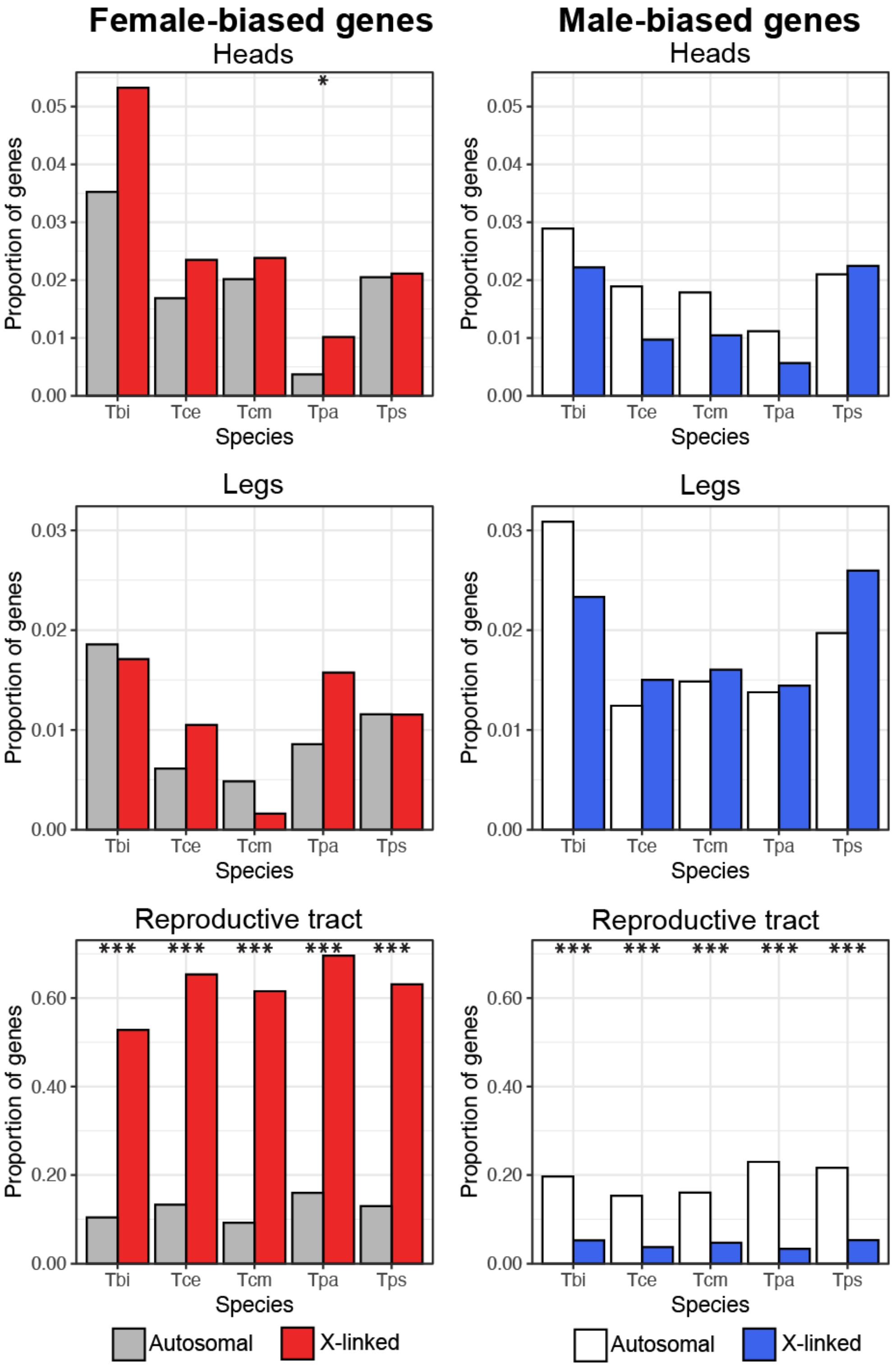


**Fig. S28 | Proportion of female- and male- biased genes on the X and autosomes in reproductive tract, head and leg samples when considering only sex-biased genes with at least a twofold difference in expression between males and females.** Note the scale changes between tissue-types. FDR: * <0.05, ** < 0.01, *** < 0.001.

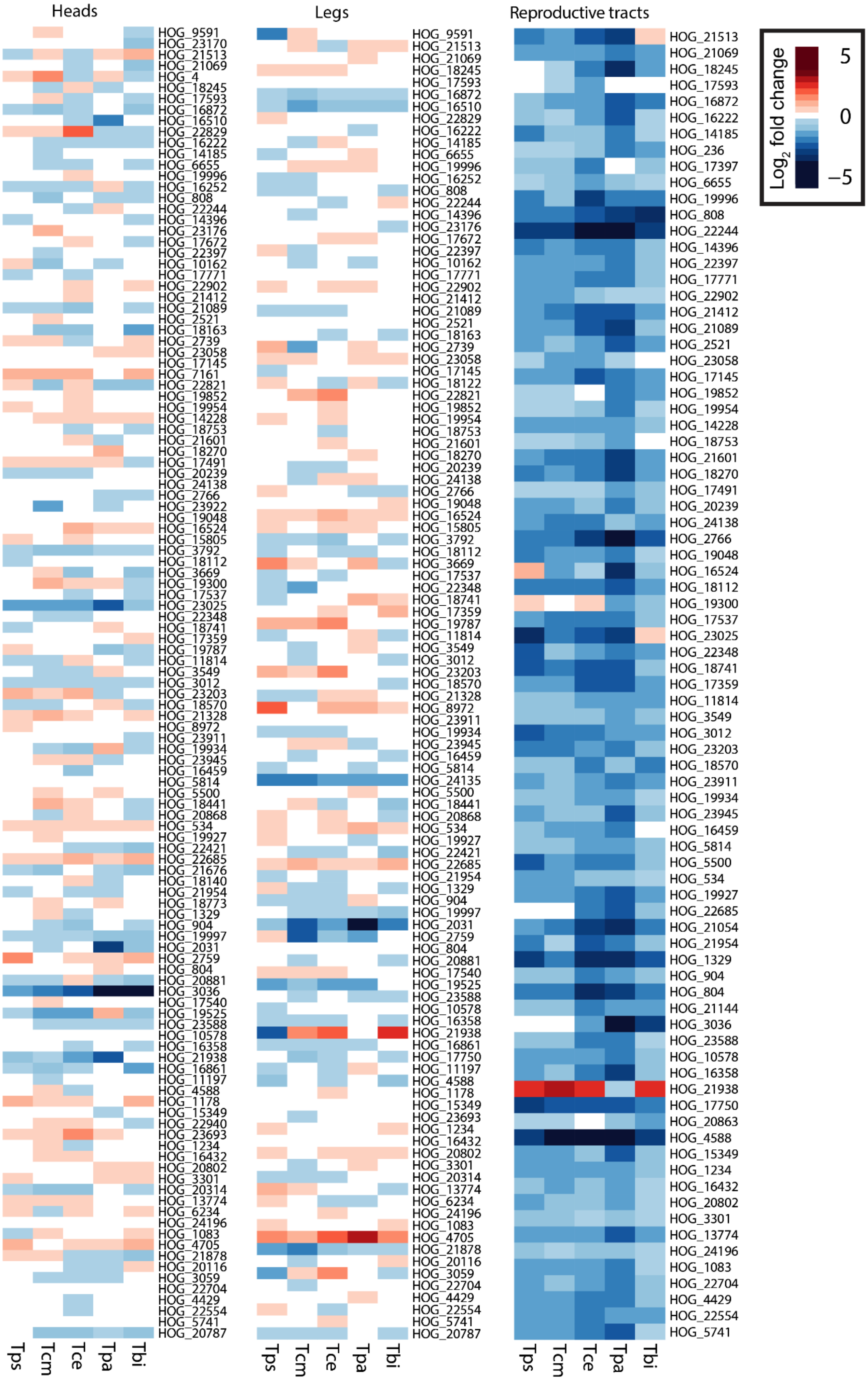


**Fig. S29 | Heatmap of gene expression of orthologs on the X in the heads, legs and reproductive tracts.** Species names are abbreviated as Tbi = *T. bartmani*, Tce = *T. cristinae*, Tcm = *T. californicum*, Tps = *T. poppensis*, and Tpa = *T. podura*.

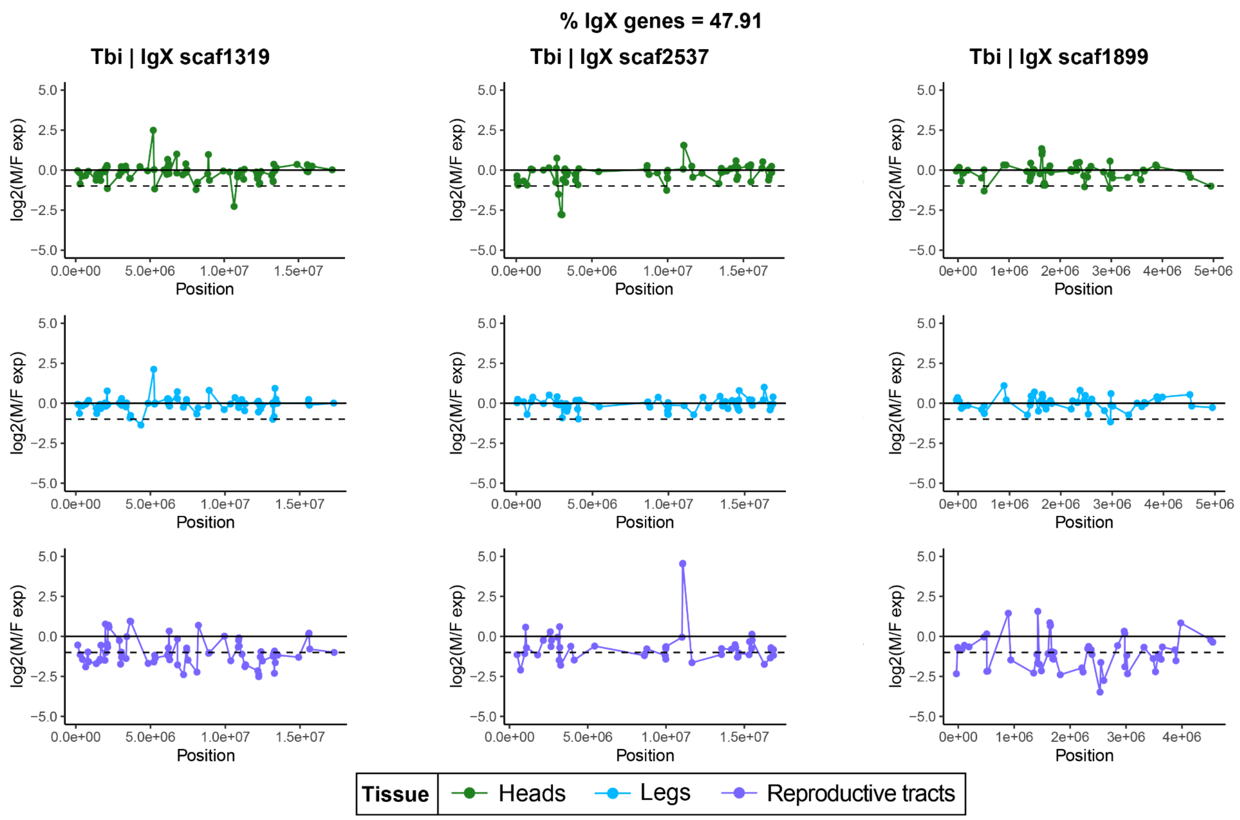


**Fig. S30 | Log₂ ratio of male to female expression along the three largest X-linked scaffolds in heads (green, top), legs (blue, middle) and reproductive tracts (purple, bottom) in *T. bartmani*.**

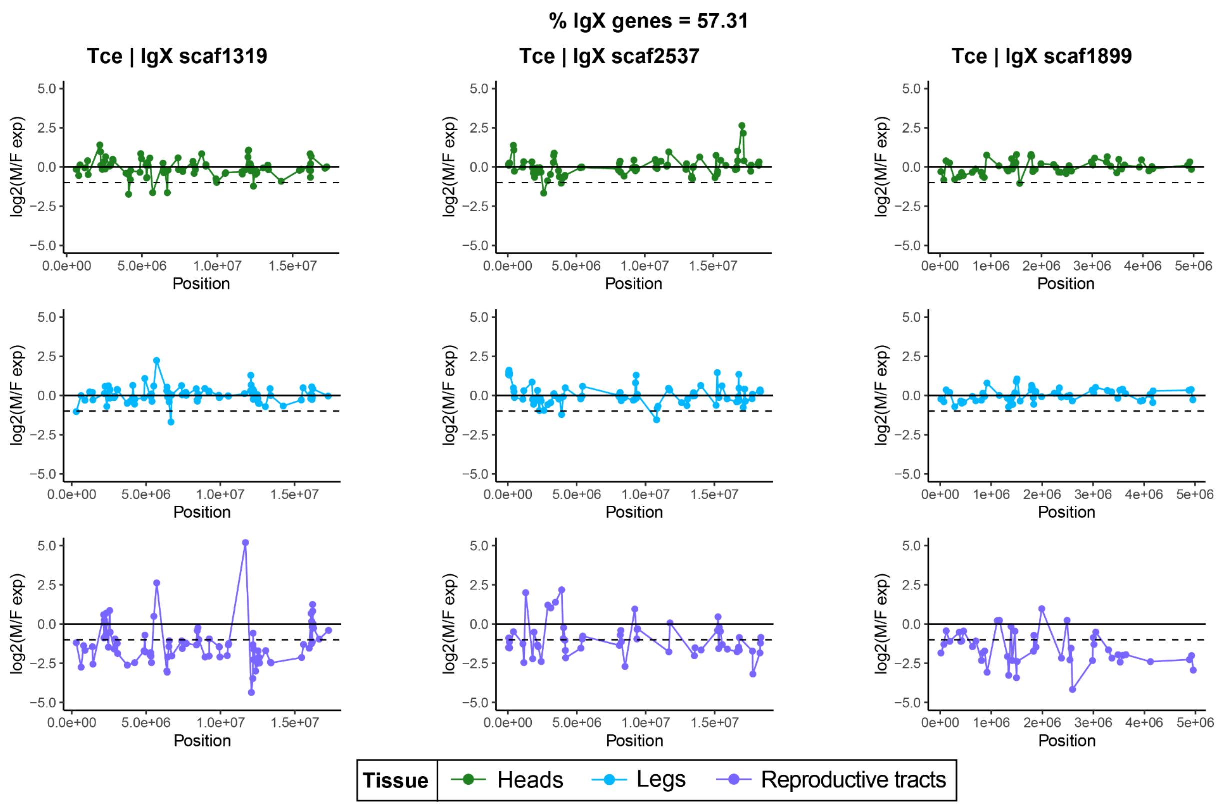


**Fig. S31 | $Log_2$ ratio of male to female expression along the three largest X-linked scaffolds in heads (green, top), legs (blue, middle) and reproductive tracts (purple, bottom) in *T. cristinae.***

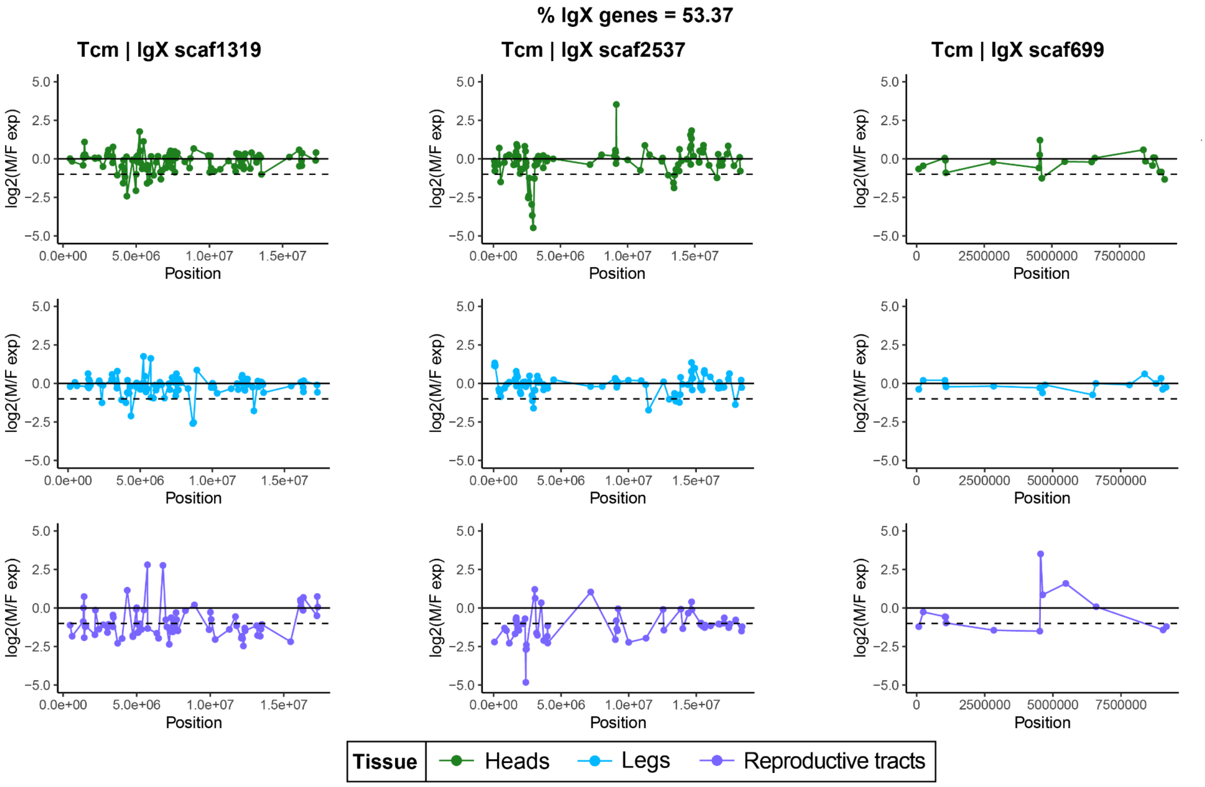


**Fig. S32 | Log$_2$ ratio of male to female expression along the three largest X-linked scaffolds in heads (green, top), legs (blue, middle) and reproductive tracts (purple, bottom) in *T. californicum.***

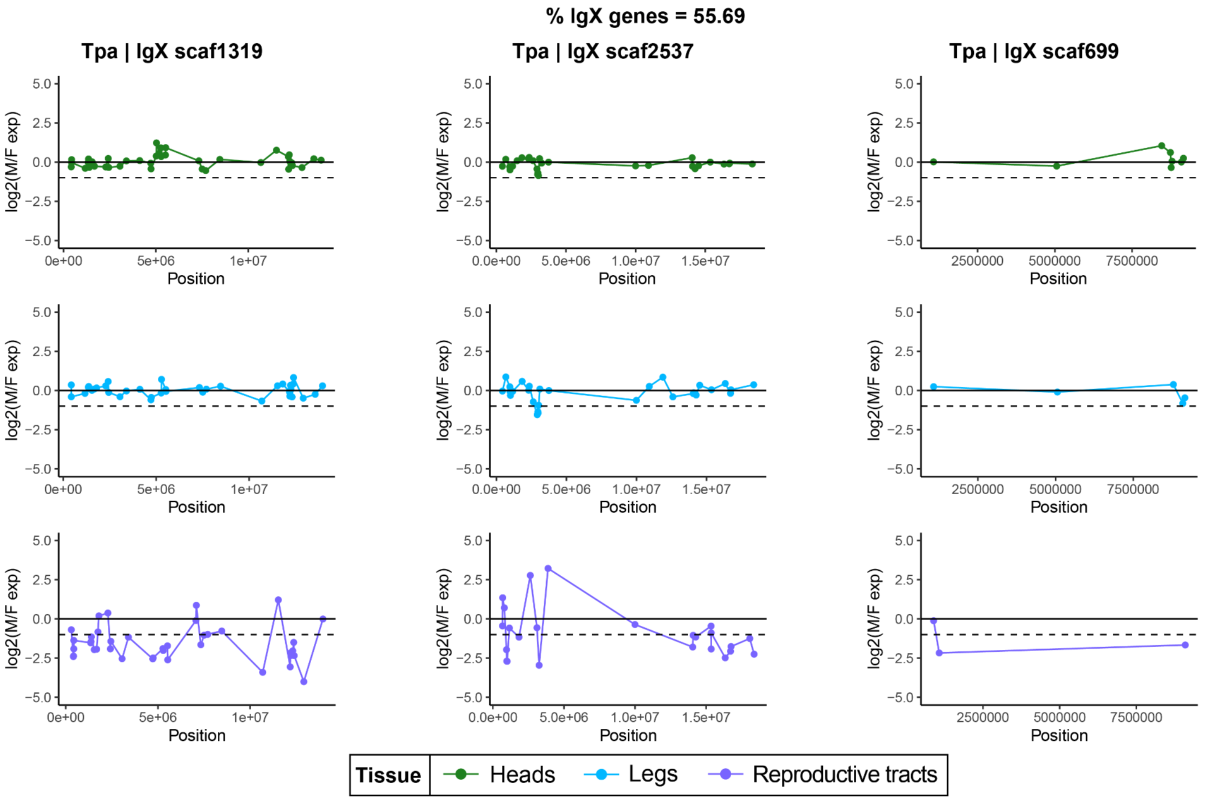


**Fig. S33 | Log$_2$ ratio of male to female expression along the three largest X-linked scaffolds in heads (green, top), legs (blue, middle) and reproductive tracts (purple, bottom) in *T. podura.***

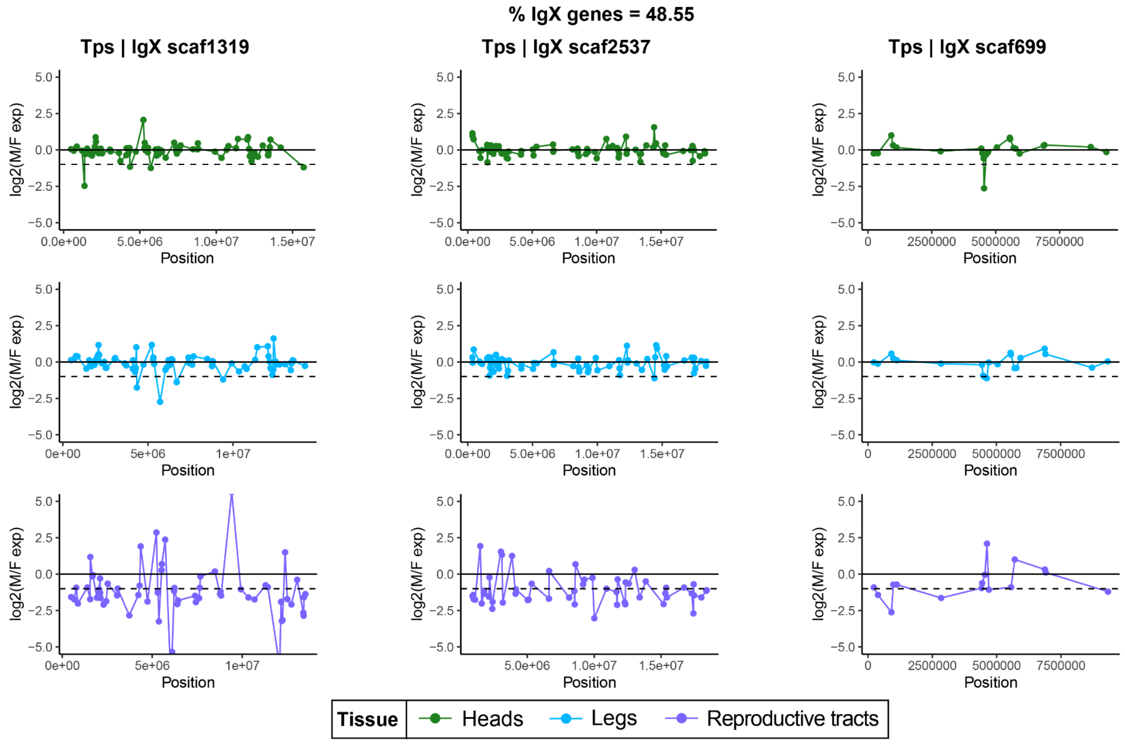


**Fig. S34 | Log$_2$ ratio of male to female expression along the three largest X-linked scaffolds in heads (green, top), legs (blue, middle) and reproductive tracts (purple, bottom) in *T. poppensis.***

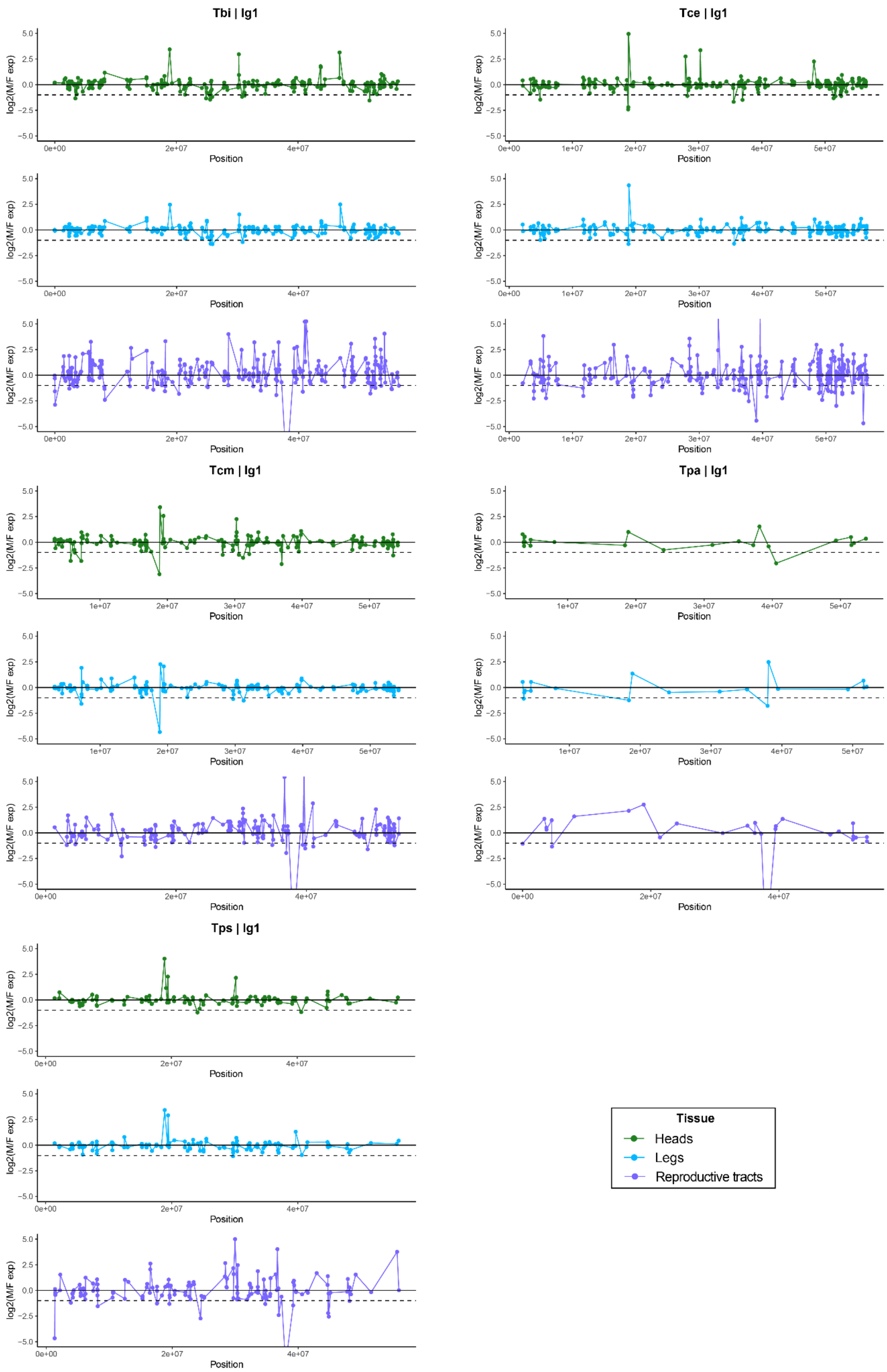


**Fig. S35 | Log$_2$ ratio of male to female expression along linkage group 1 in heads (green, top), legs (blue, middle) and reproductive tracts (purple, bottom).** Species names are abbreviated as Tbi = *T. bartmani*, Tce = *T. cristinae*, Tcm = *T. californicum*, Tps = *T. poppensis*, and Tpa = *T. podura*.

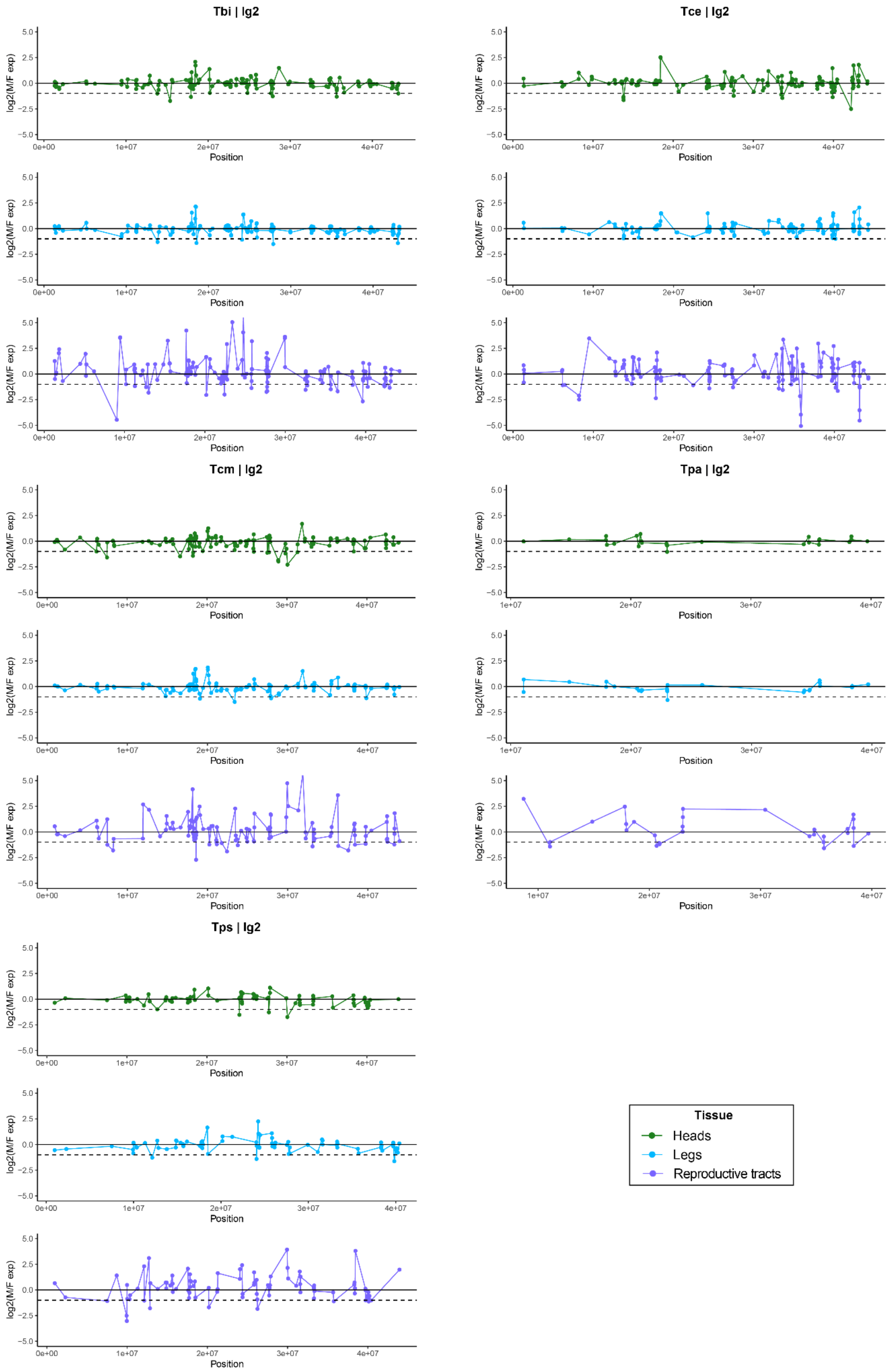


**Fig. S36 | Log$_2$ ratio of male to female expression along linkage group 2 in heads (green, top), legs (blue, middle) and reproductive tracts (purple, bottom).** Species names are abbreviated as Tbi = *T. bartmani*, Tce = *T. cristinae*, Tcm = *T. californicum*, Tps = *T. poppensis*, and Tpa = *T. podura*.

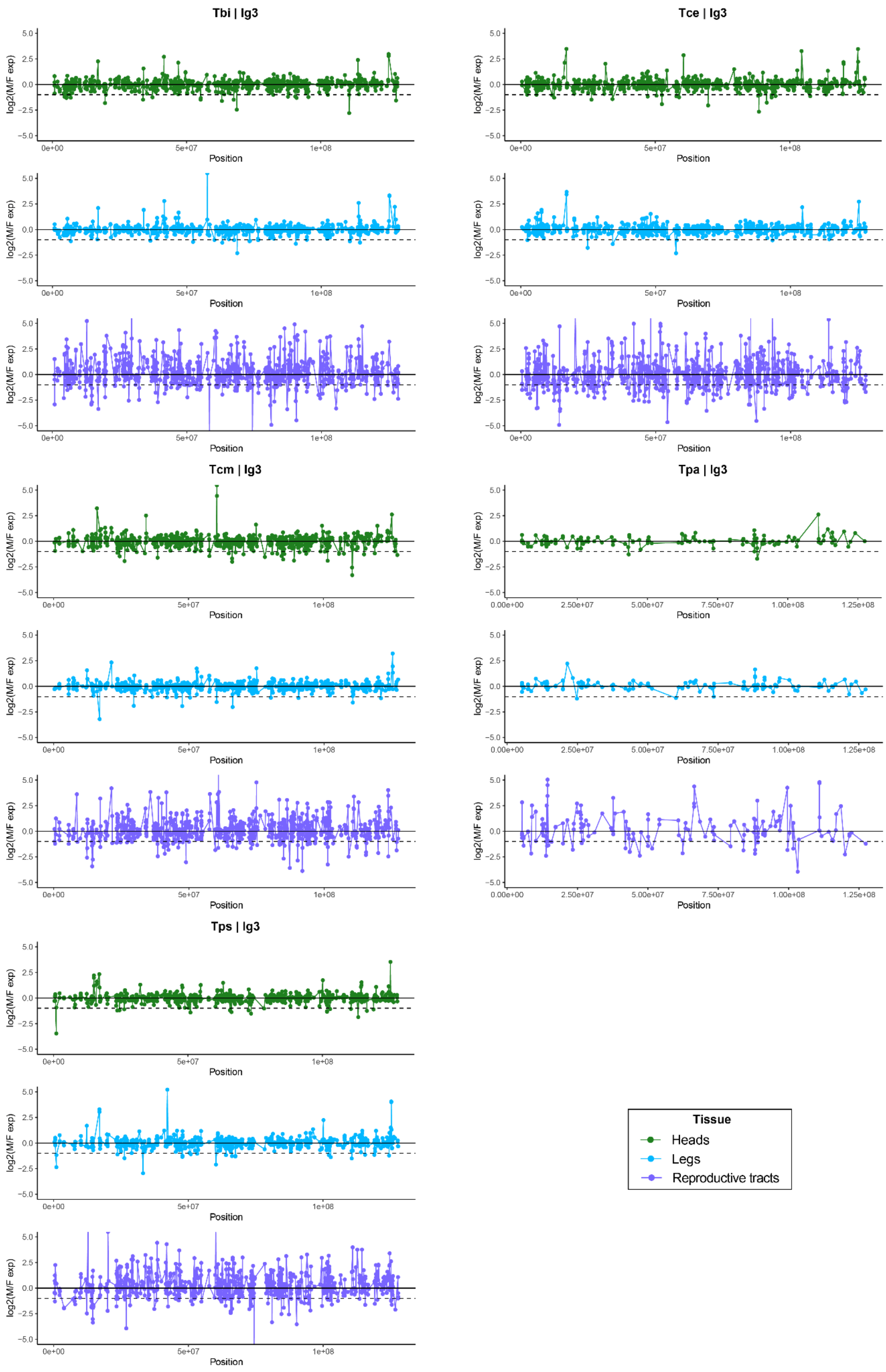


**Fig. S37 | Log₂ ratio of male to female expression along linkage group 3 in heads (green, top), legs (blue, middle) and reproductive tracts (purple, bottom).** Species names are abbreviated as Tbi = *T. bartmani*, Tce = *T. cristinae*, Tcm = *T. californicum*, Tps = *T. poppensis*, and Tpa = *T. podura*.

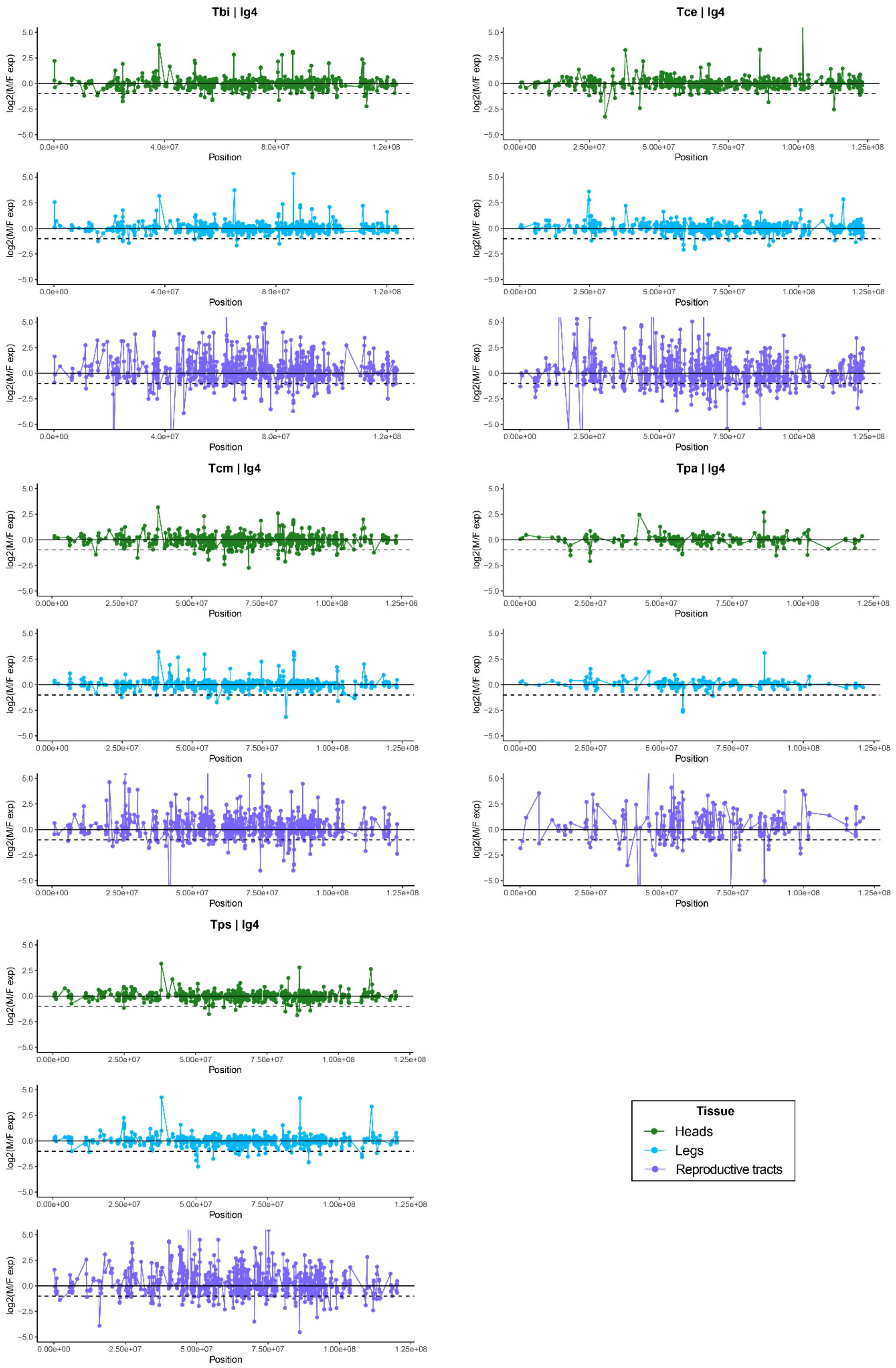


**Fig. S38 | $Log_2$ ratio of male to female expression along linkage group 4 in heads (green, top), legs (blue, middle) and reproductive tracts (purple, bottom).** Species names are abbreviated as Tbi = *T. bartmani*, Tce = *T. cristinae*, Tcm = *T. californicum*, Tps = *T. poppensis*, and Tpa = *T. podura*.

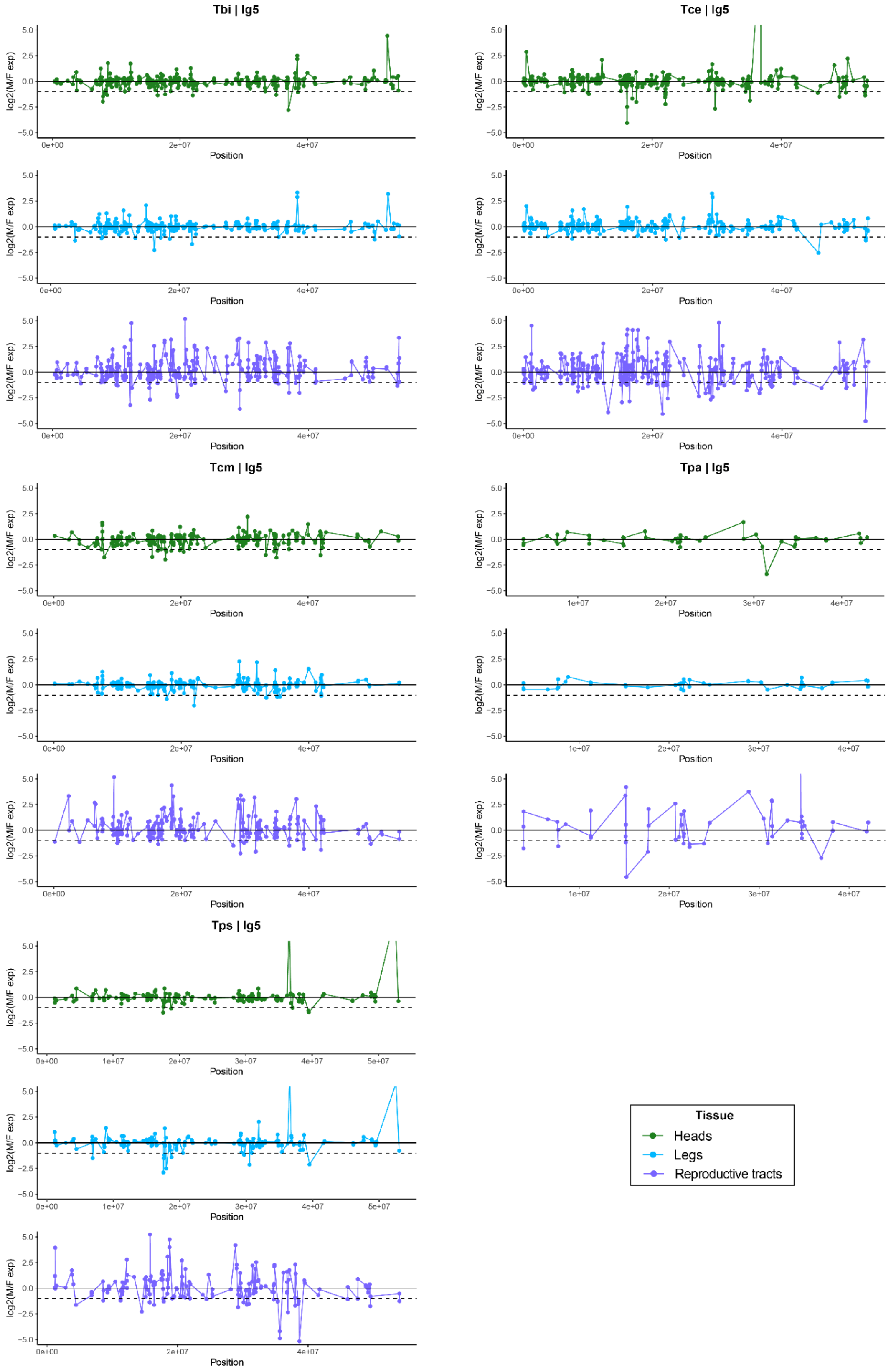


**Fig. S39 | Log$_2$ ratio of male to female expression along linkage group 5 in heads (green, top), legs (blue, middle) and reproductive tracts (purple, bottom).** Species names are abbreviated as Tbi = *T. bartmani*, Tce = *T. cristinae*, Tcm = *T. californicum*, Tps = *T. poppensis*, and Tpa = *T. podura*.

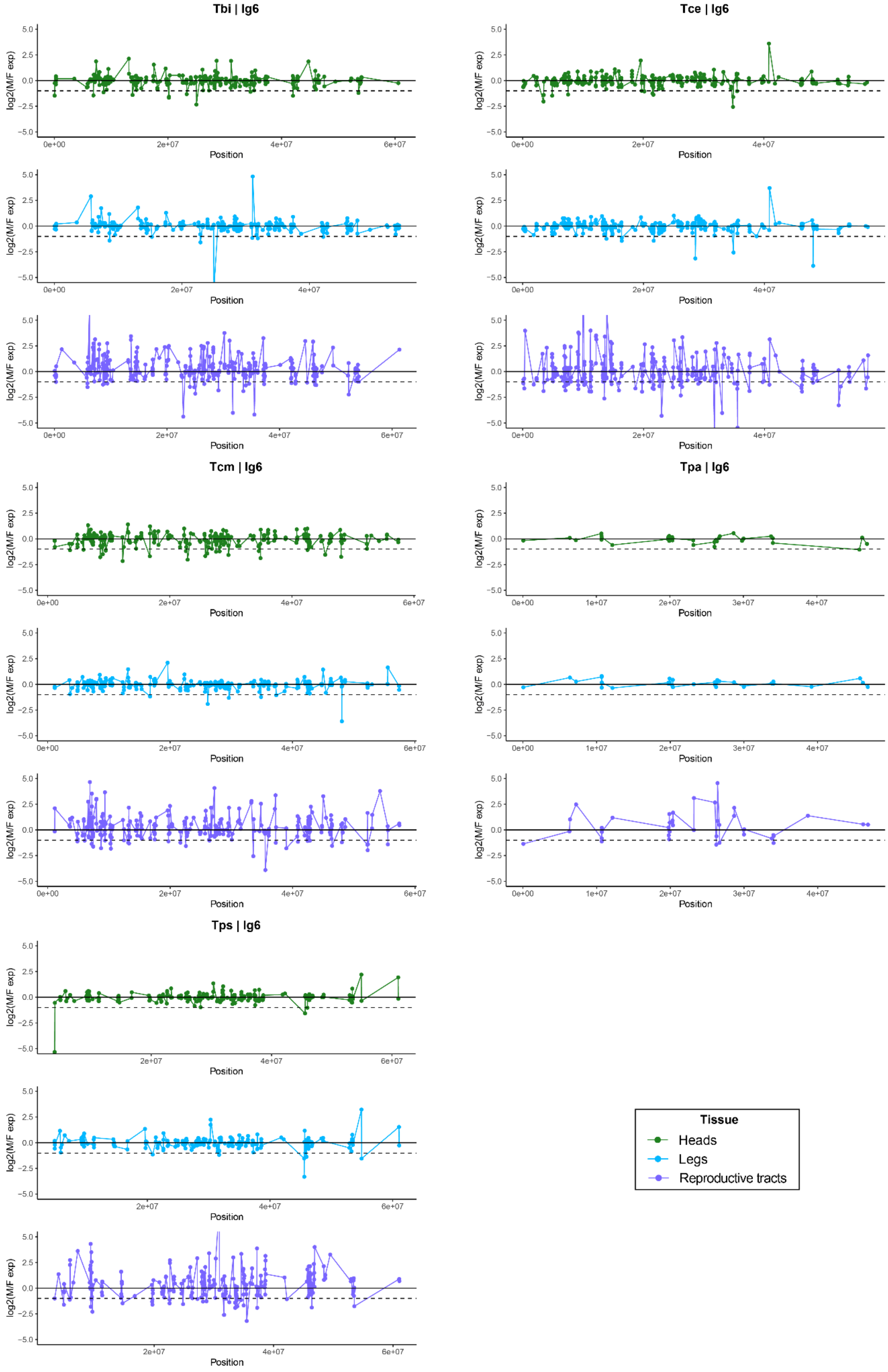


**Fig. S40 | Log$_2$ ratio of male to female expression along linkage group 6 in heads (green, top), legs (blue, middle) and reproductive tracts (purple, bottom).** Species names are abbreviated as Tbi = *T. bartmani*, Tce = *T. cristinae*, Tcm = *T. californicum*, Tps = *T. poppensis*, and Tpa = *T. podura*.

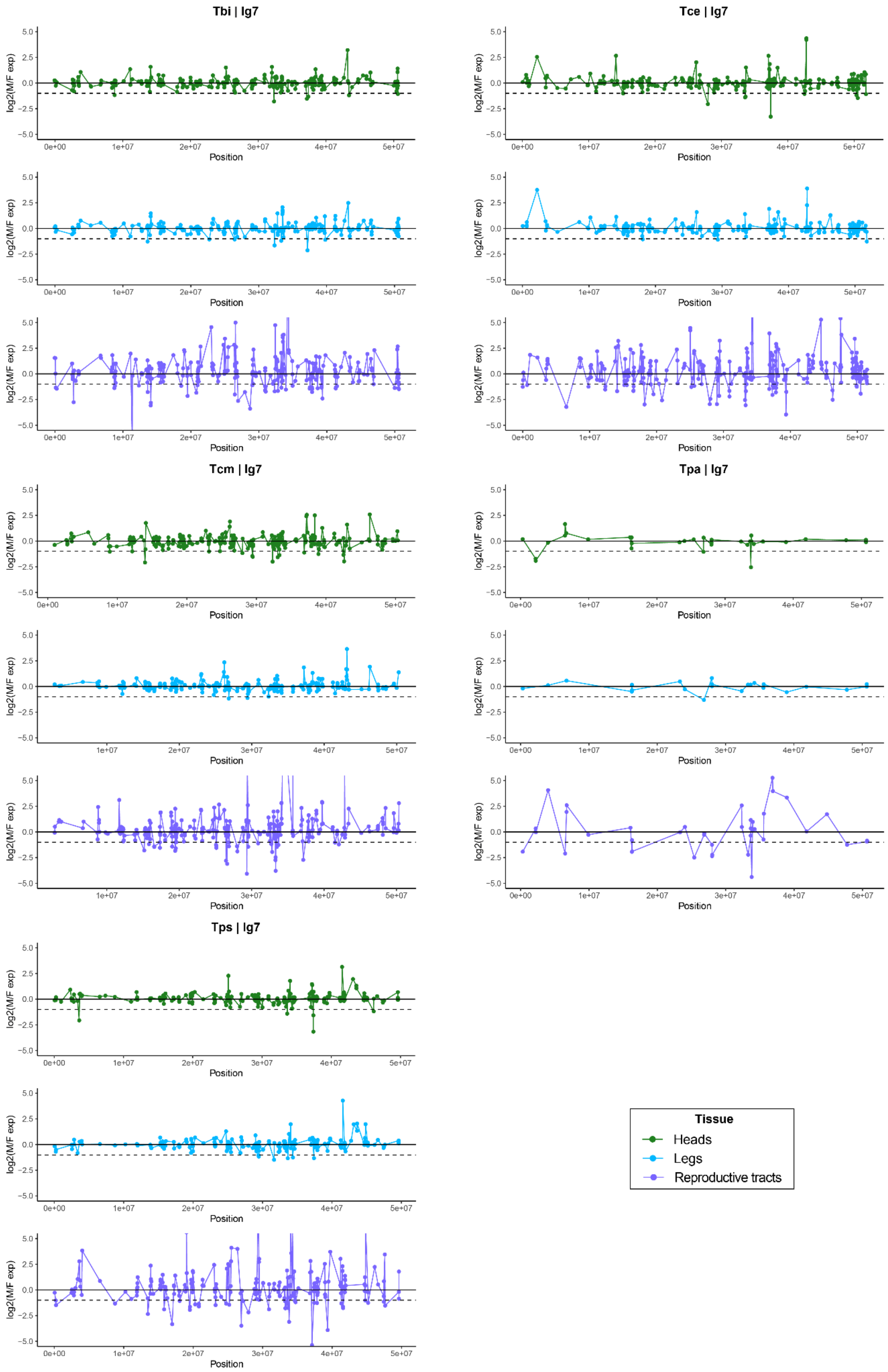


**Fig. S41 | Log$_2$ ratio of male to female expression along linkage group 7 in heads (green, top), legs (blue, middle) and reproductive tracts (purple, bottom).** Species names are abbreviated as Tbi = *T. bartmani*, Tce = *T. cristinae*, Tcm = *T. californicum*, Tps = *T. poppensis*, and Tpa = *T. podura*.

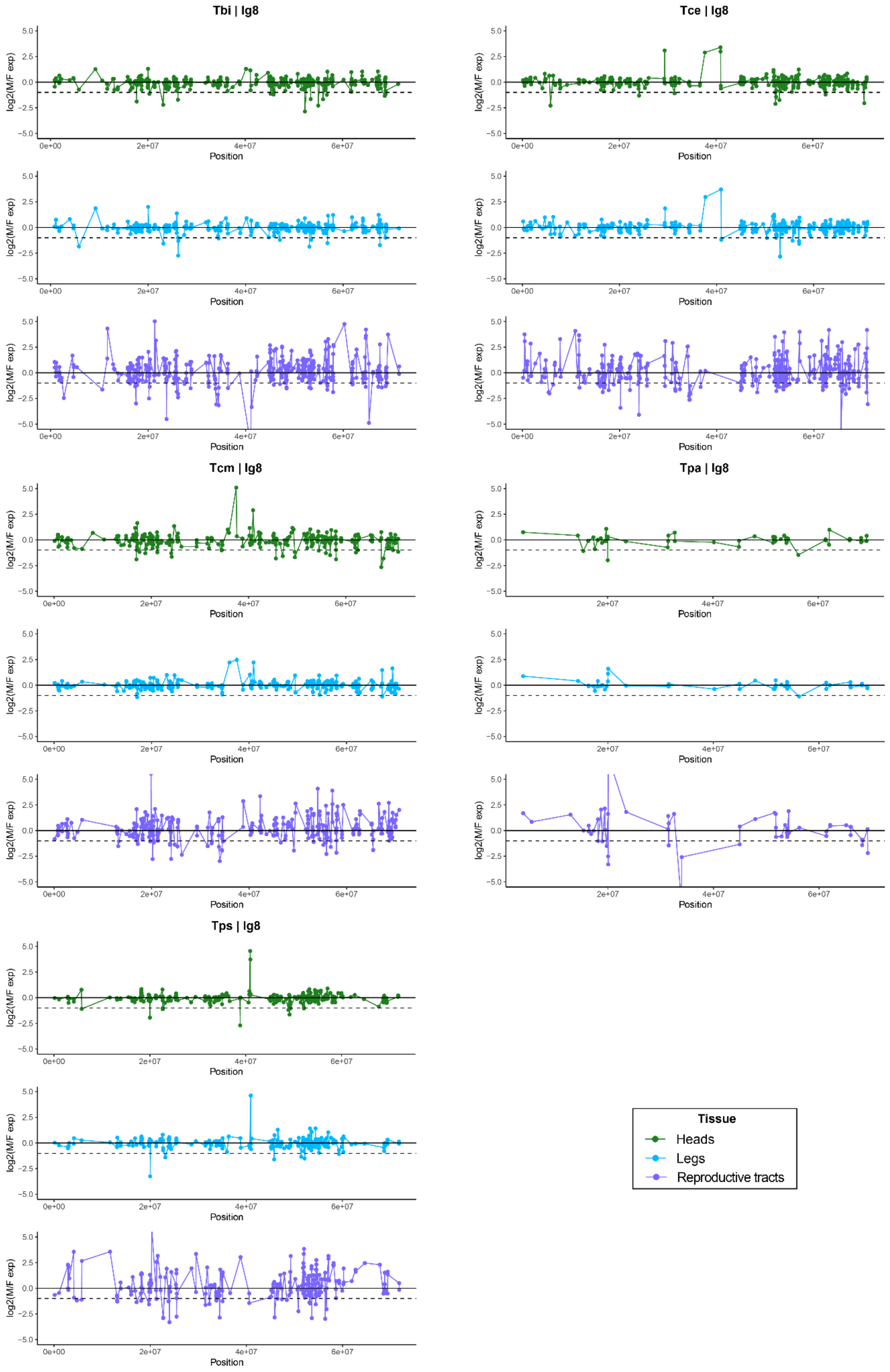


**Fig. S42 | Log$_2$ ratio of male to female expression along linkage group 8 in heads (green, top), legs (blue, middle) and reproductive tracts (purple, bottom).** Species names are abbreviated as Tbi = *T. bartmani*, Tce = *T. cristinae*, Tcm = *T. californicum*, Tps = *T. poppensis*, and Tpa = *T. podura*.

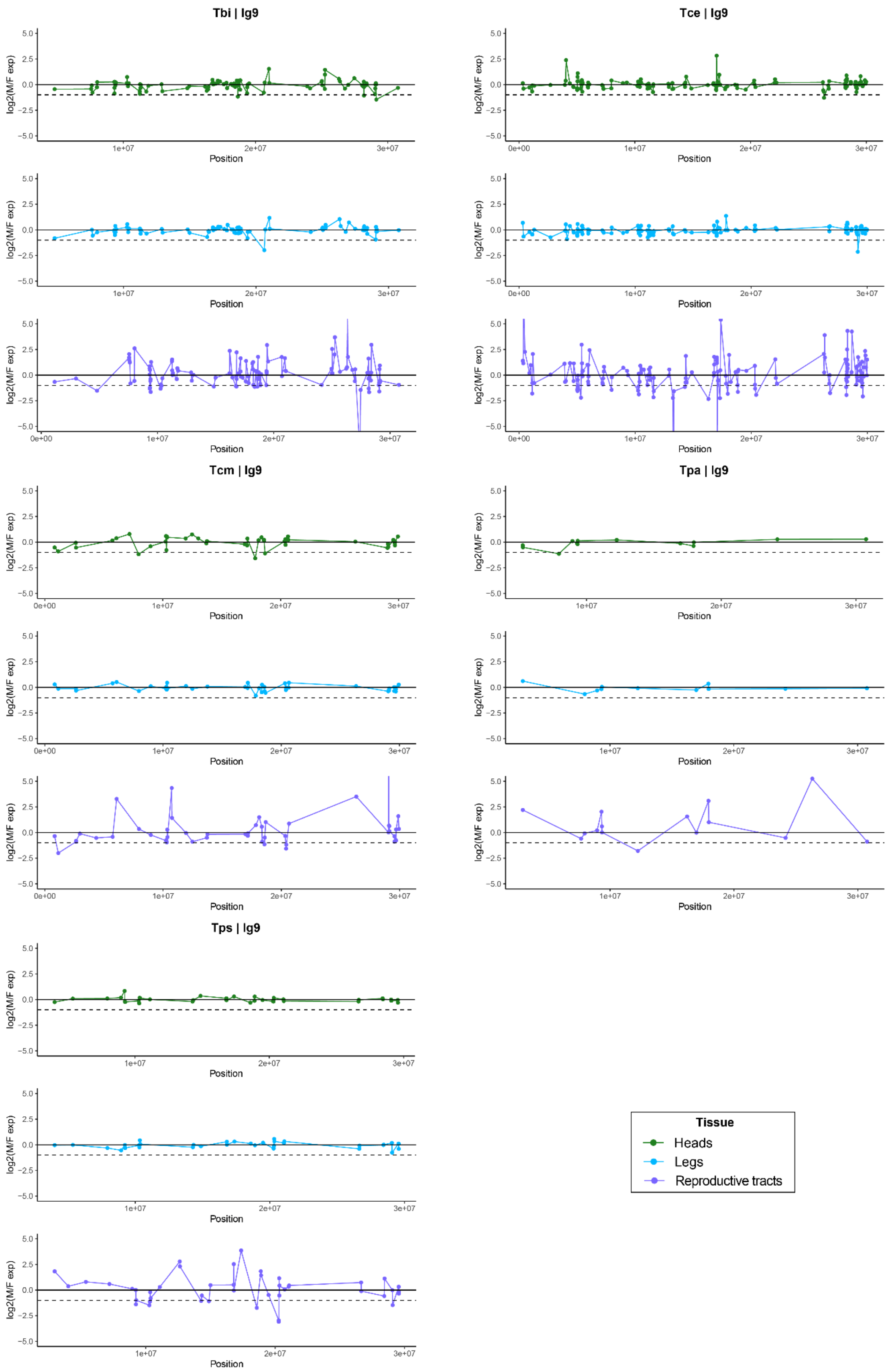


**Fig. S43 | $Log_2$ ratio of male to female expression along linkage group 9 in heads (green, top), legs (blue, middle) and reproductive tracts (purple, bottom).** Species names are abbreviated as Tbi = *T. bartmani*, Tce = *T. cristinae*, Tcm = *T. californicum*, Tps = *T. poppensis*, and Tpa = *T. podura*.

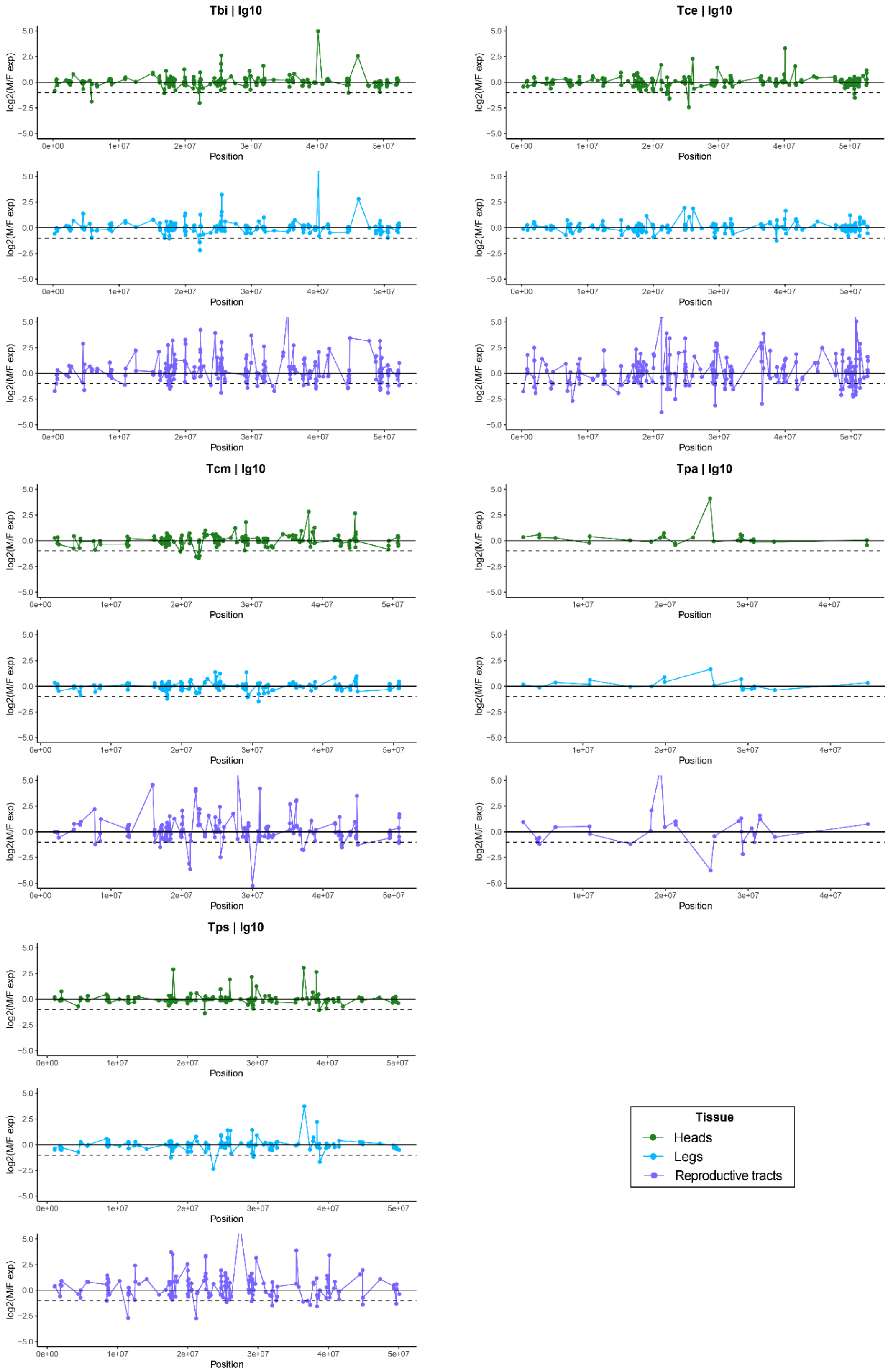


**Fig. S44 | $Log_2$ ratio of male to female expression along linkage group 10 in heads (green, top), legs (blue, middle) and reproductive tracts (purple, bottom).** Species names are abbreviated as Tbi = *T. bartmani*, Tce = *T. cristinae*, Tcm = *T. californicum*, Tps = *T. poppensis*, and Tpa = *T. podura*.

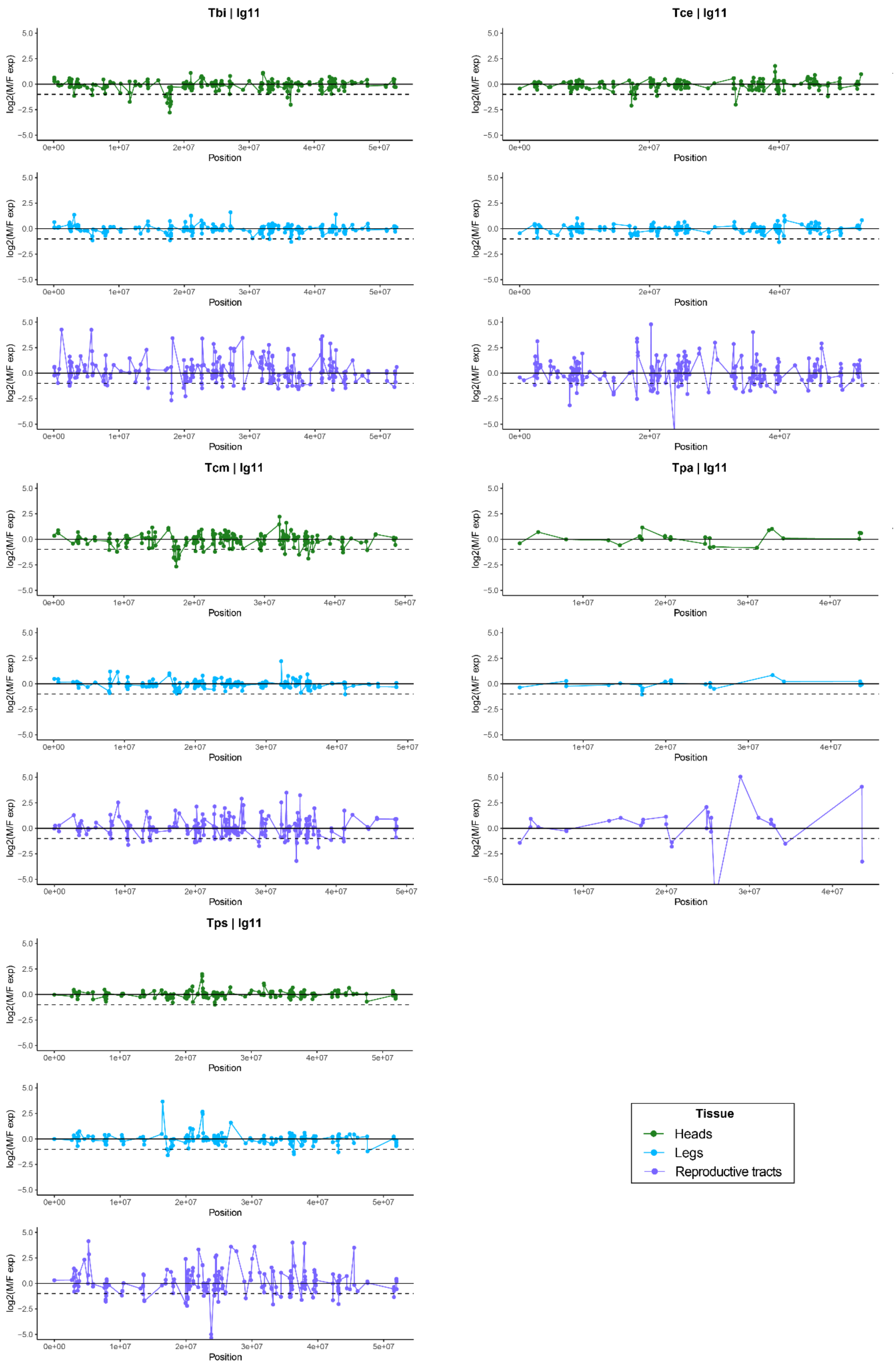


**Fig. S45 | Log$_2$ ratio of male to female expression along linkage group 11 in heads (green, top), legs (blue, middle) and reproductive tracts (purple, bottom).** Species names are abbreviated as Tbi = *T. bartmani*, Tce = *T. cristinae*, Tcm = *T. californicum*, Tps = *T. poppensis*, and Tpa = *T. podura*.

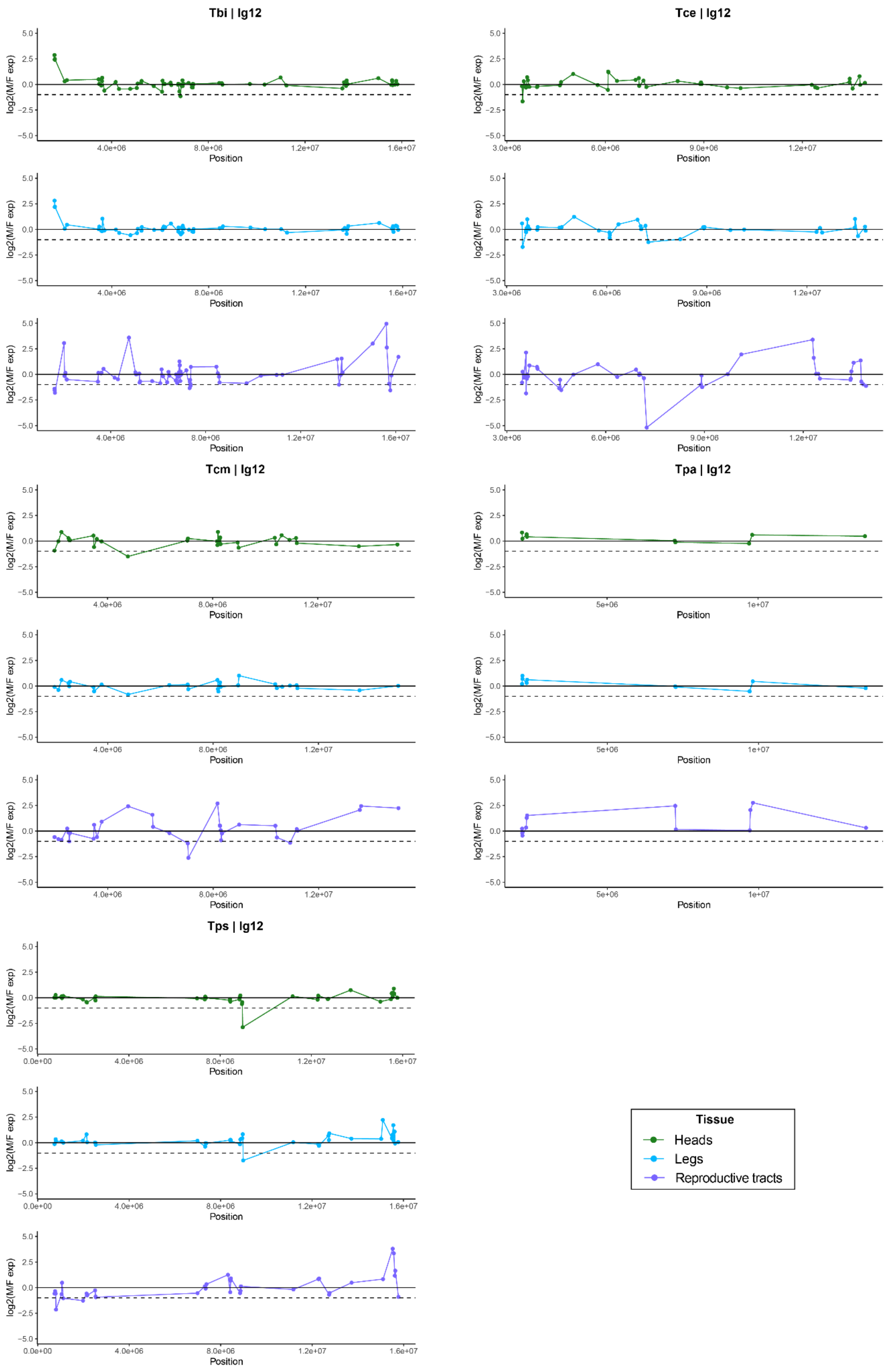


**Fig. S46 | Log$_2$ ratio of male to female expression along linkage group 12 in heads (green, top), legs (blue, middle) and reproductive tracts (purple, bottom).** Species names are abbreviated as Tbi = *T. bartmani*, Tce = *T. cristinae*, Tcm = *T. californicum*, Tps = *T. poppensis*, and Tpa = *T. podura*.

**Table S1 | Mapping statistics for each library**

| Sp | Sex | Sample name | Raw reads | Trimmed reads | Mapped reads | % trimmed reads retained | Median coverage |
|---|---|---|---|---|---|---|---|
| Tbi | F | Tbi_F_CC86B | 163550102 | 104673495 | 122491491 | 58.5 | 13 |
| Tbi | F | Tbi_F_CC86C | 182229510 | 117083861 | 157791863 | 67.4 | 16 |
| Tbi | F | Tbi_F_CC87B | 310822750 | 198768251 | 262877684 | 66.1 | 28 |
| Tbi | F | Tbi_F_CC87C | 189747769 | 120625968 | 163391747 | 67.7 | 17 |
| Tbi | F | Tbi_F_CC88B | 162185335 | 101815571 | 133042229 | 65.3 | 14 |
| Tbi | M | Tbi_M_13_Tbi | 230289434 | 193548911 | 236141867 | 61 | 25 |
| Tbi | M | Tbi_M_14_Tbi | 167317549 | 143798101 | 162886424 | 56.6 | 17 |
| Tbi | M | Tbi_M_15_Tbi | 180468659 | 159784783 | 202394255 | 63.3 | 21 |
| Tbi | M | Tbi_M_16_Tbi | 173403444 | 153356094 | 195854284 | 63.9 | 21 |
| Tce | F | Tce_F_CC22B | 240302203 | 153537662 | 194360412 | 63.3 | 21 |
| Tce | F | Tce_F_CC22C | 183847856 | 113513072 | 141789081 | 62.5 | 15 |
| Tce | F | Tce_F_CC24B | 190707946 | 121517676 | 153918581 | 63.3 | 16 |
| Tce | F | Tce_F_CC24C | 165761611 | 104261537 | 127867746 | 61.3 | 13 |
| Tce | F | Tce_F_CC25B | 173937316 | 113879289 | 143226429 | 62.9 | 15 |
| Tce | M | Tce_M_05_HM15 | 283916393 | 242069828 | 286559073 | 59.2 | 31 |
| Tce | M | Tce_M_06_HM16 | 276053436 | 233286803 | 278105695 | 59.6 | 30 |
| Tce | M | Tce_M_07_HM33 | 253435459 | 210226724 | 238032367 | 56.6 | 26 |
| Tce | M | Tce_M_08_HM61 | 252560771 | 212476439 | 253700845 | 59.7 | 27 |
| Tcm | F | Tcm_F_HM217 | 185583334 | 117066283 | 132790722 | 56.7 | 13 |
| Tcm | F | Tcm_F_HM218 | 186876879 | 119895740 | 134255101 | 56 | 14 |
| Tcm | F | Tcm_F_HM219 | 201939048 | 129727246 | 148122395 | 57.1 | 15 |
| Tcm | F | Tcm_F_HM220 | 165585380 | 103738590 | 117786957 | 56.8 | 12 |
| Tcm | F | Tcm_F_HM221 | 169629803 | 108266912 | 121750485 | 56.2 | 12 |
| Tcm | M | Tcm_M_01_HM148 | 308854833 | 264941657 | 290114343 | 54.8 | 30 |
| Tcm | M | Tcm_M_02_HM149 | 257853423 | 216836107 | 235138570 | 54.2 | 24 |
| Tcm | M | Tcm_M_03_HM150 | 220274671 | 185844153 | 199230991 | 53.6 | 20 |
| Tcm | M | Tcm_M_04_HM151 | 252568065 | 213772777 | 233051440 | 54.5 | 24 |
| Tpa | F | Tpa_F_H54 | 161627958 | 105644951 | 120542760 | 57.1 | 11 |
| Tpa | F | Tpa_F_H56 | 271827902 | 173075827 | 187636415 | 54.2 | 18 |
| Tpa | F | Tpa_F_Pa_AB | 163854265 | 106235524 | 127703269 | 60.1 | 12 |
| Tpa | F | Tpa_F_PA_CD | 172588238 | 110487637 | 128870754 | 58.3 | 13 |
| Tpa | F | Tpa_F_PA_E | 162728669 | 106103692 | 130725613 | 61.6 | 12 |
| Tpa | M | Tpa_M_09_Tpa | 230642771 | 192844114 | 222442319 | 57.7 | 21 |
| Tpa | M | Tpa_M_10_Tpa | 270408484 | 225888550 | 264701771 | 58.6 | 24 |
| Tpa | M | Tpa_M_11_Tpa | 287173420 | 240787111 | 278536113 | 57.8 | 26 |
| Tpa | M | Tpa_M_12_Tpa | 277374266 | 236821164 | 271814085 | 57.4 | 26 |
| Tps | F | Tps_F_ReSeq_Ps08 | 134510710 | 114179907 | 112990604 | 49.5 | 11 |
| Tps | F | Tps_F_ReSeq_Ps12 | 144601431 | 123763989 | 119753101 | 48.4 | 12 |
| Tps | F | Tps_F_ReSeq_Ps14 | 152252645 | 132604275 | 151769843 | 57.2 | 16 |

| | | | | | | | |
|---|---|---|---|---|---|---|---|
| Tps | F | Tps_F_ReSeq_Ps16 | 121136075 | 105142652 | 129026061 | 61.4 | 13 |
| Tps | F | Tps_F_ReSeq_Ps18 | 153052014 | 132974234 | 165681524 | 62.3 | 17 |
| Tps | M | Tps_M_17_HM99 | 174653591 | 154114880 | 182631522 | 59.3 | 19 |
| Tps | M | Tps_M_18_HM100 | 177718034 | 156454093 | 178690981 | 57.1 | 19 |
| Tps | M | Tps_M_19_HM101 | 162978918 | 142304334 | 169701197 | 59.6 | 18 |
| Tps | M | Tps_M_20_15255 | 151428008 | 132551815 | 148456301 | 56 | 16 |

**Table S2 | Number of genes and scaffolds classified as X-linked or autosomal using a more stringent coverage classification**

| Species | Min scaffold length | % genome X-linked | N Autosomal genes | N X-linked genes | N genes not classified |
|---|---|---|---|---|---|
| *T. bartmani* | 1000 | 10.84 | 12923 | 1125 | 18 |
| *T. cristinae* | 1000 | 10.11 | 12730 | 1128 | 24 |
| *T. californicum* | 1000 | 11.02 | 13360 | 1191 | 12 |
| *T. podura* | 1000 | 7.39 | 15680 | 846 | 3 |
| *T. poppensis* | 1000 | 10.79 | 14331 | 1238 | 36 |

**Table S3 | Number of genes and scaffolds classified as X-linked or autosomal with a minimum scaffold length of 5000.** For coverage classification scheme, M = main analysis, S = stringent classification.

| Species | Min scaffold length | % genome X-linked | N Autosomal genes | N X-linked genes | N genes not classified | Classification scheme |
|---|---|---|---|---|---|---|
| *T. bartmani* | 5000 | 11.89 | 11913 | 1184 | 969 | M |
| *T. cristinae* | 5000 | 11.97 | 11620 | 1251 | 1011 | M |
| *T. californicum* | 5000 | 12.57 | 11285 | 1257 | 2021 | M |
| *T. podura* | 5000 | 15.42 | 9873 | 1234 | 5422 | M |
| *T. poppensis* | 5000 | 13.06 | 11587 | 1337 | 2681 | M |
| *T. bartmani* | 5000 | 11.08 | 12012 | 1085 | 969 | S |
| *T. cristinae* | 5000 | 10.32 | 11763 | 1108 | 1011 | S |
| *T. californicum* | 5000 | 11.88 | 11407 | 1135 | 2021 | S |
| *T. podura* | 5000 | 10.41 | 10250 | 857 | 5422 | S |
| *T. poppensis* | 5000 | 12.15 | 11746 | 1178 | 2681 | S |

**Table S4 | Nucleotide diversity (π) and effective population size (Ne) estimates.** Nucleotide diversity estimates are median values of scaffolds weighted by scaffold length. Effective population sizes were calculated using the mutation rate either from *Heliconius melpomene* (H) (Keightley *et al.*, 2015) or *Drosophila melanogaster* (D) (Keightley *et al.*, 2014).

| Species | $\pi_X$ | $\pi_A$ | $\pi_X / \pi_A$ | $Ne_X$ (H) | $Ne_A$ (H) | $Ne_X$ (D) | $Ne_A$ (D) |
|---|---|---|---|---|---|---|---|
| *T. bartmani* | 0.00087 | 0.00328 | 0.26521 | 74939 | 282566 | 77616 | 292658 |

| | | | | | | | |
|---|---|---|---|---|---|---|---|
| *T. cristinae* | 0.00358 | 0.00887 | 0.40386 | 308764 | 764537 | 319792 | 791842 |
| *T. californicum* | 0.00110 | 0.00578 | 0.19034 | 94889 | 498513 | 98278 | 516317 |
| *T. podura* | 0.01049 | 0.02204 | 0.47582 | 903955 | 1899767 | 936239 | 1967616 |
| *T. poppensis* | 0.00078 | 0.00189 | 0.41475 | 67490 | 162724 | 69900 | 168535 |

**Table S5 | Enrichment of SB genes on the X when using the FPKM gene set.** Sp = species (Species names are abbreviated as Tbi = *T. bartmani*, Tce = *T. cristinae*, Tcm = *T. californicum*, Tps = *T. poppensis*, and Tpa = *T. podura*), Tiss = tissue (RT = reproductive tract, HD = head, LD = legs), Chr_c = coverage classification (M = main analysis, S = stringent classification). SB_class indicates how sex-biased genes were classified (FDR = FDR < 0.05 only, FDR + FC = FDR < 0.05 + FC > 2), FBX = number of female-biased genes on the X, FBA = number of female-biased genes on the autosomes, MBX = number of male-biased genes on the X, MBA = number of female-biased genes on the autosomes, AllX = total number of genes expressed on the X, AllA = total number of genes expressed on the autosomes, FB_FDR = fisher's exact test FDR for enrichment of female-biased genes, MB_FDR = fisher's exact test FDR for enrichment of male-biased genes. FDR < 0.05 in bold

| Sp | Tiss | chr_c | SB_class | FBX | FBA | MBX | MBA | AllX | AllA | FB_FDR | MB_FDR |
|---|---|---|---|---|---|---|---|---|---|---|---|
| Tbi | RT | S | FDR | 380 | 2221 | 37 | 2629 | 484 | 7604 | **1.32E-103** | **2.32E-41** |
| Tce | RT | S | FDR | 321 | 1185 | 16 | 1218 | 471 | 7575 | **1.12E-130** | **3.47E-17** |
| Tcm | RT | S | FDR | 299 | 1032 | 20 | 1467 | 451 | 7329 | **1.79E-128** | **3.32E-20** |
| Tpa | RT | S | FDR | 328 | 2324 | 11 | 2660 | 435 | 10149 | **3.54E-112** | **1.03E-39** |
| Tps | RT | S | FDR | 418 | 2377 | 35 | 2776 | 530 | 8353 | **1.16E-118** | **1.10E-45** |
| Tbi | HD | S | FDR | 100 | 698 | 29 | 605 | 611 | 6984 | **8.74E-06** | **1.08E-03** |
| Tce | HD | S | FDR | 15 | 143 | 9 | 154 | 622 | 7033 | 6.40E-01 | 4.14E-01 |
| Tcm | HD | S | FDR | 16 | 143 | 9 | 128 | 626 | 6935 | 5.23E-01 | 7.97E-01 |
| Tpa | HD | S | FDR | 5 | 36 | 3 | 100 | 523 | 9058 | 1.33E-01 | 5.68E-01 |
| Tps | HD | S | FDR | 29 | 277 | 35 | 281 | 694 | 8010 | 4.99E-01 | 9.42E-02 |
| Tbi | LG | S | FDR | 48 | 408 | 36 | 440 | 582 | 6574 | 1.33E-01 | 8.40E-01 |
| Tce | LG | S | FDR | 5 | 46 | 8 | 92 | 568 | 6619 | 6.41E-01 | 9.14E-01 |
| Tcm | LG | S | FDR | 1 | 33 | 11 | 99 | 590 | 6634 | 6.40E-01 | 6.55E-01 |
| Tpa | LG | S | FDR | 7 | 77 | 6 | 113 | 445 | 8012 | 3.53E-01 | 1.00E+00 |
| Tps | LG | S | FDR | 12 | 136 | 22 | 177 | 640 | 7407 | 8.78E-01 | 2.06E-01 |
| Tbi | RT | S | FDR + FC | 265 | 805 | 27 | 1487 | 484 | 7604 | **8.27E-114** | **5.13E-17** |
| Tce | RT | S | FDR + FC | 317 | 1046 | 16 | 1152 | 471 | 7575 | **5.25E-141** | **1.72E-15** |
| Tcm | RT | S | FDR + FC | 280 | 696 | 20 | 1172 | 451 | 7329 | **4.06E-148** | **2.46E-13** |
| Tpa | RT | S | FDR + FC | 320 | 1771 | 10 | 2277 | 435 | 10149 | **2.49E-136** | **5.17E-32** |
| Tps | RT | S | FDR + FC | 344 | 1103 | 26 | 1804 | 530 | 8353 | **1.40E-150** | **6.59E-25** |
| Tbi | HD | S | FDR + FC | 27 | 253 | 13 | 202 | 611 | 6984 | 5.60E-01 | 4.66E-01 |
| Tce | HD | S | FDR + FC | 14 | 120 | 6 | 132 | 622 | 7033 | 5.60E-01 | 2.88E-01 |
| Tcm | HD | S | FDR + FC | 15 | 140 | 7 | 123 | 626 | 6935 | 6.93E-01 | 4.39E-01 |
| Tpa | HD | S | FDR + FC | 5 | 36 | 3 | 99 | 523 | 9058 | 1.77E-01 | 5.15E-01 |
| Tps | HD | S | FDR + FC | 14 | 165 | 16 | 168 | 694 | 8010 | 1.00E+00 | 7.45E-01 |
| Tbi | LG | S | FDR + FC | 9 | 123 | 12 | 204 | 582 | 6574 | 8.00E-01 | 3.83E-01 |
| Tce | LG | S | FDR + FC | 5 | 42 | 8 | 83 | 568 | 6619 | 6.26E-01 | 7.45E-01 |
| Tcm | LG | S | FDR + FC | 1 | 32 | 10 | 98 | 590 | 6634 | 6.93E-01 | 7.45E-01 |
| Tpa | LG | S | FDR + FC | 7 | 71 | 6 | 111 | 445 | 8012 | 4.13E-01 | 1.00E+00 |
| Tps | LG | S | FDR + FC | 6 | 87 | 18 | 145 | 640 | 7407 | 8.00E-01 | 3.06E-01 |
| Tbi | RT | M | FDR | 409 | 2192 | 41 | 2625 | 534 | 7554 | **1.73E-105** | **6.85E-46** |
| Tce | RT | M | FDR | 370 | 1136 | 21 | 1213 | 560 | 7486 | **6.38E-146** | **4.10E-19** |
| Tcm | RT | M | FDR | 322 | 1009 | 24 | 1463 | 489 | 7291 | **3.95E-138** | **3.21E-20** |
| Tpa | RT | M | FDR | 533 | 2119 | 27 | 2644 | 746 | 9838 | **8.64E-169** | **6.71E-60** |
| Tps | RT | M | FDR | 448 | 2347 | 44 | 2767 | 583 | 8300 | **3.03E-120** | **1.79E-46** |

| | | | | | | | | | | | |
|---|---|---|---|---|---|---|---|---|---|---|---|
| Tbi | HD | M | FDR | 113 | 685 | 37 | 597 | 676 | 6919 | **5.98E-07** | **8.80E-03** |
| Tce | HD | M | FDR | 18 | 140 | 10 | 153 | 723 | 6932 | 4.37E-01 | 2.91E-01 |
| Tcm | HD | M | FDR | 17 | 142 | 9 | 128 | 671 | 6890 | 4.37E-01 | 6.09E-01 |
| Tpa | HD | M | FDR | 9 | 32 | 5 | 98 | 886 | 8695 | **2.38E-02** | 2.91E-01 |
| Tps | HD | M | FDR | 32 | 274 | 37 | 279 | 757 | 7947 | 3.84E-01 | 1.42E-01 |
| Tbi | LG | M | FDR | 54 | 402 | 44 | 432 | 643 | 6513 | 6.36E-02 | 8.04E-01 |
| Tce | LG | M | FDR | 7 | 44 | 10 | 90 | 666 | 6521 | 4.37E-01 | 7.81E-01 |
| Tcm | LG | M | FDR | 1 | 33 | 11 | 99 | 624 | 6600 | 4.37E-01 | 7.26E-01 |
| Tpa | LG | M | FDR | 13 | 71 | 12 | 107 | 762 | 7695 | 8.67E-02 | 7.26E-01 |
| Tps | LG | M | FDR | 15 | 133 | 22 | 177 | 693 | 7354 | 4.61E-01 | 3.03E-01 |
| Tbi | RT | M | FDR + FC | 282 | 788 | 28 | 1486 | 534 | 7554 | **2.06E-116** | **6.32E-20** |
| Tce | RT | M | FDR + FC | 366 | 997 | 21 | 1147 | 560 | 7486 | **1.91E-158** | **4.25E-17** |
| Tcm | RT | M | FDR + FC | 301 | 675 | 23 | 1169 | 489 | 7291 | **5.99E-159** | **1.08E-13** |
| Tpa | RT | M | FDR + FC | 519 | 1572 | 25 | 2262 | 746 | 9838 | **2.26E-209** | **4.94E-48** |
| Tps | RT | M | FDR + FC | 368 | 1079 | 31 | 1799 | 583 | 8300 | **1.23E-156** | **6.24E-26** |
| Tbi | HD | M | FDR + FC | 36 | 244 | 15 | 200 | 676 | 6919 | 5.15E-02 | 5.37E-01 |
| Tce | HD | M | FDR + FC | 17 | 117 | 7 | 131 | 723 | 6932 | 3.00E-01 | 1.96E-01 |
| Tcm | HD | M | FDR + FC | 16 | 139 | 7 | 123 | 671 | 6890 | 5.96E-01 | 3.95E-01 |
| Tpa | HD | M | FDR + FC | 9 | 32 | 5 | 97 | 886 | 8695 | **2.77E-02** | 3.57E-01 |
| Tps | HD | M | FDR + FC | 16 | 163 | 17 | 167 | 757 | 7947 | 1.00E+00 | 8.47E-01 |
| Tbi | LG | M | FDR + FC | 11 | 121 | 15 | 201 | 643 | 6513 | 1.00E+00 | 5.01E-01 |
| Tce | LG | M | FDR + FC | 7 | 40 | 10 | 81 | 666 | 6521 | 3.00E-01 | 7.29E-01 |
| Tcm | LG | M | FDR + FC | 1 | 32 | 10 | 98 | 624 | 6600 | 4.90E-01 | 8.44E-01 |
| Tpa | LG | M | FDR + FC | 12 | 66 | 11 | 106 | 762 | 7695 | 1.30E-01 | 8.70E-01 |
| Tps | LG | M | FDR + FC | 8 | 85 | 18 | 145 | 693 | 7354 | 1.00E+00 | 4.31E-01 |

**Table S6 | Enrichment of SB genes on the X when using the TPM gene set.** Sp = species (Species names are abbreviated as Tbi = *T. bartmani*, Tce = *T. cristinae*, Tcm = *T. californicum*, Tps = *T. poppensis*, and Tpa = *T. podura*), Tiss = tissue (RT = reproductive tract, HD = head, LD = legs), Chr_c = coverage classification (M = main analysis, S = stringent classification). SB_class indicates how sex-biased genes were classified (FDR = FDR < 0.05 only, FDR + FC = FDR < 0.05 + FC > 2), FBX = number of female-biased genes on the X, FBA = number of female-biased genes on the autosomes, MBX = number of male-biased genes on the X, MBA = number of female-biased genes on the autosomes, AllX = total number of genes expressed on the X, AllA = total number of genes expressed on the autosomes, FB_FDR = fisher's exact test FDR for enrichment of female-biased genes, MB_FDR = fisher's exact test FDR for enrichment of male-biased genes. FDR values < 0.05 are given in bold.

| Sp | Tiss | chr_c | SB_class | FBX | FBA | MBX | MBA | AllX | AllA | FB_FDR | MB_FDR |
|---|---|---|---|---|---|---|---|---|---|---|---|
| Tbi | RT | S | FDR | 454 | 2400 | 48 | 2887 | 584 | 8310 | **1.19E-121** | **1.63E-47** |
| Tce | RT | S | FDR | 366 | 1107 | 21 | 1466 | 553 | 8076 | **2.82E-158** | **7.11E-23** |
| Tcm | RT | S | FDR | 348 | 1092 | 24 | 1634 | 538 | 7917 | **7.01E-146** | **4.53E-25** |
| Tpa | RT | S | FDR | 316 | 2257 | 11 | 2590 | 419 | 9914 | **5.29E-109** | **1.12E-37** |
| Tps | RT | S | FDR | 476 | 2495 | 39 | 2968 | 607 | 8962 | **2.36E-136** | **1.02E-52** |
| Tbi | HD | S | FDR | 103 | 750 | 34 | 629 | 691 | 7625 | **1.52E-04** | **3.86E-03** |
| Tce | HD | S | FDR | 14 | 141 | 13 | 165 | 683 | 7590 | 7.61E-01 | 8.60E-01 |
| Tcm | HD | S | FDR | 17 | 160 | 9 | 133 | 669 | 7492 | 6.64E-01 | 8.04E-01 |
| Tpa | HD | S | FDR | 5 | 36 | 3 | 99 | 523 | 9061 | 1.51E-01 | 6.29E-01 |
| Tps | HD | S | FDR | 28 | 263 | 38 | 310 | 751 | 8556 | 5.21E-01 | 1.20E-01 |
| Tbi | LG | S | FDR | 45 | 426 | 40 | 460 | 652 | 7080 | 5.21E-01 | 8.60E-01 |
| Tce | LG | S | FDR | 5 | 47 | 9 | 94 | 621 | 7080 | 7.61E-01 | 8.60E-01 |
| Tcm | LG | S | FDR | 2 | 34 | 11 | 105 | 620 | 6901 | 7.66E-01 | 8.30E-01 |
| Tpa | LG | S | FDR | 7 | 79 | 6 | 116 | 455 | 8149 | 4.17E-01 | 1.00E+00 |
| Tps | LG | S | FDR | 10 | 132 | 21 | 173 | 677 | 7743 | 7.66E-01 | 2.68E-01 |
| Tbi | RT | S | FDR + FC | 335 | 946 | 35 | 1754 | 584 | 8310 | **1.96E-141** | **3.22E-22** |
| Tce | RT | S | FDR + FC | 362 | 1022 | 21 | 1396 | 553 | 8076 | **1.10E-164** | **6.16E-21** |
| Tcm | RT | S | FDR + FC | 331 | 771 | 23 | 1348 | 538 | 7917 | **8.29E-169** | **3.47E-18** |
| Tpa | RT | S | FDR + FC | 307 | 1710 | 10 | 2199 | 419 | 9914 | **1.41E-131** | **4.15E-30** |
| Tps | RT | S | FDR + FC | 402 | 1235 | 30 | 2017 | 607 | 8962 | **2.29E-173** | **3.52E-30** |
| Tbi | HD | S | FDR + FC | 33 | 305 | 17 | 229 | 691 | 7625 | 5.89E-01 | 7.15E-01 |
| Tce | HD | S | FDR + FC | 14 | 123 | 10 | 144 | 683 | 7590 | 6.97E-01 | 7.15E-01 |
| Tcm | HD | S | FDR + FC | 16 | 157 | 8 | 128 | 669 | 7492 | 7.42E-01 | 7.15E-01 |
| Tpa | HD | S | FDR + FC | 5 | 36 | 3 | 98 | 523 | 9061 | 1.77E-01 | 7.06E-01 |
| Tps | HD | S | FDR + FC | 14 | 169 | 18 | 185 | 751 | 8556 | 1.00E+00 | 8.02E-01 |
| Tbi | LG | S | FDR + FC | 10 | 145 | 15 | 222 | 652 | 7080 | 6.97E-01 | 6.10E-01 |
| Tce | LG | S | FDR + FC | 5 | 44 | 9 | 87 | 621 | 7080 | 7.42E-01 | 7.15E-01 |
| Tcm | LG | S | FDR + FC | 2 | 34 | 10 | 104 | 620 | 6901 | 8.20E-01 | 9.25E-01 |
| Tpa | LG | S | FDR + FC | 7 | 73 | 6 | 114 | 455 | 8149 | 4.27E-01 | 1.00E+00 |
| Tps | LG | S | FDR + FC | 6 | 90 | 18 | 145 | 677 | 7743 | 8.13E-01 | 3.68E-01 |
| Tbi | RT | M | FDR | 489 | 2365 | 53 | 2882 | 645 | 8249 | **4.85E-124** | **8.33E-53** |
| Tce | RT | M | FDR | 421 | 1052 | 27 | 1460 | 654 | 7975 | **8.43E-178** | **1.48E-25** |
| Tcm | RT | M | FDR | 373 | 1067 | 28 | 1630 | 578 | 7877 | **4.01E-157** | **1.52E-25** |
| Tpa | RT | M | FDR | 516 | 2057 | 24 | 2577 | 713 | 9620 | **4.42E-169** | **9.17E-59** |
| Tps | RT | M | FDR | 515 | 2456 | 49 | 2958 | 673 | 8896 | **3.01E-140** | **2.18E-54** |
| Tbi | HD | M | FDR | 117 | 736 | 42 | 621 | 765 | 7551 | **1.14E-05** | **1.57E-02** |

| | | | | | | | | | | | |
|---|---|---|---|---|---|---|---|---|---|---|---|
| Tce | HD | M | FDR | 18 | 137 | 14 | 164 | 794 | 7479 | 5.09E-01 | 7.09E-01 |
| Tcm | HD | M | FDR | 18 | 159 | 9 | 133 | 715 | 7446 | 5.77E-01 | 5.54E-01 |
| Tpa | HD | M | FDR | 9 | 32 | 5 | 97 | 887 | 8697 | **2.39E-02** | 3.12E-01 |
| Tps | HD | M | FDR | 31 | 260 | 40 | 308 | 823 | 8484 | 4.39E-01 | 1.77E-01 |
| Tbi | LG | M | FDR | 51 | 420 | 48 | 452 | 720 | 7012 | 4.19E-01 | 8.11E-01 |
| Tce | LG | M | FDR | 7 | 45 | 11 | 92 | 727 | 6974 | 4.57E-01 | 7.64E-01 |
| Tcm | LG | M | FDR | 2 | 34 | 11 | 105 | 656 | 6865 | 8.21E-01 | 8.03E-01 |
| Tpa | LG | M | FDR | 13 | 73 | 12 | 110 | 778 | 7826 | 1.07E-01 | 8.03E-01 |
| Tps | LG | M | FDR | 13 | 129 | 21 | 173 | 737 | 7683 | 8.80E-01 | 5.05E-01 |
| Tbi | RT | M | FDR + FC | 358 | 923 | 38 | 1751 | 645 | 8249 | **6.81E-146** | **4.03E-25** |
| Tce | RT | M | FDR + FC | 417 | 967 | 27 | 1390 | 654 | 7975 | **6.99E-186** | **2.30E-23** |
| Tcm | RT | M | FDR + FC | 354 | 748 | 26 | 1345 | 578 | 7877 | **2.35E-181** | **7.52E-19** |
| Tpa | RT | M | FDR + FC | 500 | 1517 | 22 | 2187 | 713 | 9620 | **1.07E-206** | **7.75E-47** |
| Tps | RT | M | FDR + FC | 435 | 1202 | 36 | 2011 | 673 | 8896 | **3.27E-183** | **5.90E-32** |
| Tbi | HD | M | FDR + FC | 43 | 295 | 19 | 227 | 765 | 7551 | 5.74E-02 | 6.26E-01 |
| Tce | HD | M | FDR + FC | 18 | 119 | 11 | 143 | 794 | 7479 | 3.09E-01 | 5.60E-01 |
| Tcm | HD | M | FDR + FC | 17 | 156 | 8 | 128 | 715 | 7446 | 7.33E-01 | 5.60E-01 |
| Tpa | HD | M | FDR + FC | 9 | 32 | 5 | 96 | 887 | 8697 | **2.79E-02** | 4.14E-01 |
| Tps | HD | M | FDR + FC | 16 | 167 | 19 | 184 | 823 | 8484 | 1.00E+00 | 9.26E-01 |
| Tbi | LG | M | FDR + FC | 12 | 143 | 18 | 219 | 720 | 7012 | 7.33E-01 | 6.26E-01 |
| Tce | LG | M | FDR + FC | 7 | 42 | 11 | 85 | 727 | 6974 | 3.33E-01 | 6.26E-01 |
| Tcm | LG | M | FDR + FC | 2 | 34 | 10 | 104 | 656 | 6865 | 8.84E-01 | 1.00E+00 |
| Tpa | LG | M | FDR + FC | 12 | 68 | 11 | 109 | 778 | 7826 | 1.40E-01 | 9.36E-01 |
| Tps | LG | M | FDR + FC | 8 | 88 | 18 | 145 | 737 | 7683 | 1.00E+00 | 5.60E-01 |

**Table S7 | Wilcoxon tests comparing expression between X and autosomes in males and females for the FPKM and TPM gene sets using either the main coverage classification or the more stringent one.** Sp = species (Species names are abbreviated as Tbi = *T. bartmani*, Tce = *T. cristinae*, Tcm = *T. californicum*, Tps = *T. poppensis*, and Tpa = *T. podura*), Tiss = tissue (RT = reproductive tract, HD = head, LD = legs), Comparison = gene expression being compared (FemaleA - MaleA = expression on female autosomes vs expression on male autosomes, FemaleA - MaleX = expression on female autosomes vs expression on male X, FemaleX - FemaleA = expression on female X vs expression on female autosomes, FemaleX - MaleA = expression on female X vs expression on male autosomes, FemaleX - MaleX = expression on female X vs expression on male X, MaleX - MaleA = expression on male X vs expression on male autosomes), FPKM Main FDR = Wilcoxon test FDR for the expression comparison when classing genes with the main coverage classification and FPKM, FPKM Stringent FDR = Wilcoxon test FDR for the expression comparison when classing genes with the stringent coverage classification and FPKM, TPM Main FDR = Wilcoxon test FDR for the expression comparison when classing genes with the main coverage classification and TPM, TPM Stringent FDR = Wilcoxon test FDR for the expression comparison when classing genes with the stringent coverage classification and TPM. FDR values < 0.05 are given in bold.

| Sp | Tiss | Comparison | FPKM Main FDR | FPKM Stringent FDR | TPM Main FDR | TPM Stringent FDR |
|---|---|---|---|---|---|---|
| Tbi | HD | FemaleA - MaleA | 1.89E-01 | 1.96E-01 | **2.42E-02** | **2.28E-02** |
| Tbi | HD | FemaleA - MaleX | **1.02E-03** | **9.39E-04** | **6.10E-07** | **9.44E-07** |
| Tbi | HD | FemaleX - FemaleA | 5.49E-01 | 4.73E-01 | 5.82E-02 | 6.26E-02 |
| Tbi | HD | FemaleX - MaleA | 9.45E-01 | 8.80E-01 | 4.42E-01 | 4.22E-01 |
| Tbi | HD | FemaleX - MaleX | 8.36E-02 | 9.80E-02 | **2.75E-02** | **3.63E-02** |
| Tbi | HD | MaleX - MaleA | **1.19E-02** | **1.14E-02** | **1.83E-04** | **2.25E-04** |
| Tbi | LG | FemaleA - MaleA | 8.48E-01 | 8.18E-01 | 7.25E-02 | 8.02E-02 |
| Tbi | LG | FemaleA - MaleX | 4.10E-01 | 2.85E-01 | 1.25E-01 | 9.00E-02 |
| Tbi | LG | FemaleX - FemaleA | 3.18E-01 | 2.59E-01 | **1.18E-02** | **7.63E-03** |
| Tbi | LG | FemaleX - MaleA | 2.67E-01 | 2.23E-01 | **1.10E-03** | **7.75E-04** |
| Tbi | LG | FemaleX - MaleX | 9.71E-01 | 9.84E-01 | 5.56E-01 | 5.83E-01 |
| Tbi | LG | MaleX - MaleA | 3.38E-01 | 2.47E-01 | **2.42E-02** | **1.57E-02** |
| Tbi | RT | FemaleA - MaleA | **1.02E-16** | **5.28E-16** | **1.19E-16** | **6.17E-16** |
| Tbi | RT | FemaleA - MaleX | **5.93E-19** | **1.99E-17** | **3.39E-31** | **3.55E-29** |
| Tbi | RT | FemaleX - FemaleA | **3.58E-03** | **2.18E-03** | 2.06E-01 | 1.29E-01 |
| Tbi | RT | FemaleX - MaleA | 9.81E-01 | 7.95E-01 | 8.38E-02 | 2.41E-01 |
| Tbi | RT | FemaleX - MaleX | **4.25E-22** | **8.97E-21** | **2.25E-24** | **1.75E-23** |
| Tbi | RT | MaleX - MaleA | **2.16E-34** | **4.92E-31** | **2.90E-51** | **4.94E-47** |
| Tce | HD | FemaleA - MaleA | 9.31E-01 | 8.80E-01 | 5.58E-01 | 5.08E-01 |
| Tce | HD | FemaleA - MaleX | **9.55E-03** | **2.01E-02** | **6.99E-04** | **2.06E-03** |
| Tce | HD | FemaleX - FemaleA | 1.05E-01 | 1.34E-01 | **4.44E-02** | **4.70E-02** |
| Tce | HD | FemaleX - MaleA | 1.30E-01 | 1.63E-01 | 9.19E-02 | 1.01E-01 |
| Tce | HD | FemaleX - MaleX | 5.96E-01 | 6.92E-01 | 3.77E-01 | 5.06E-01 |
| Tce | HD | MaleX - MaleA | **1.24E-02** | **2.76E-02** | **2.45E-03** | **6.79E-03** |
| Tce | LG | FemaleA - MaleA | 4.23E-01 | 4.49E-01 | **1.56E-08** | **1.58E-08** |
| Tce | LG | FemaleA - MaleX | 9.78E-02 | 1.65E-01 | 7.42E-01 | 7.97E-01 |
| Tce | LG | FemaleX - FemaleA | 9.78E-02 | 1.34E-01 | **1.68E-02** | **2.14E-02** |
| Tce | LG | FemaleX - MaleA | **3.68E-02** | 6.13E-02 | **6.20E-07** | **2.71E-06** |

| | | | | | | |
|---|---|---|---|---|---|---|
| Tce | LG | FemaleX - MaleX | 9.82E-01 | 9.50E-01 | 1.19E-01 | 1.33E-01 |
| Tce | LG | MaleX - MaleA | **3.68E-02** | 8.13E-02 | **7.00E-03** | **1.59E-02** |
| Tce | RT | FemaleA - MaleA | **1.24E-03** | **7.06E-03** | **5.68E-31** | **1.74E-28** |
| Tce | RT | FemaleA - MaleX | **2.08E-47** | **2.19E-44** | **1.24E-42** | **9.71E-41** |
| Tce | RT | FemaleX - FemaleA | **2.99E-03** | **1.04E-02** | **4.13E-02** | 9.81E-02 |
| Tce | RT | FemaleX - MaleA | 1.23E-01 | 1.63E-01 | **1.69E-02** | **3.12E-02** |
| Tce | RT | FemaleX - MaleX | **1.56E-43** | **1.04E-39** | **1.32E-36** | **1.71E-33** |
| Tce | RT | MaleX - MaleA | **1.54E-56** | **4.11E-51** | **3.55E-74** | **2.42E-67** |
| Tcm | HD | FemaleA - MaleA | **4.38E-03** | **4.44E-03** | **2.85E-14** | **2.15E-14** |
| Tcm | HD | FemaleA - MaleX | **1.22E-02** | **2.14E-02** | **1.03E-04** | **1.51E-04** |
| Tcm | HD | FemaleX - FemaleA | 9.45E-01 | 9.77E-01 | 6.86E-01 | 7.14E-01 |
| Tcm | HD | FemaleX - MaleA | 3.70E-01 | 3.54E-01 | **5.85E-04** | **9.40E-04** |
| Tcm | HD | FemaleX - MaleX | 1.01E-01 | 1.34E-01 | **1.71E-03** | **2.53E-03** |
| Tcm | HD | MaleX - MaleA | 2.19E-01 | 2.85E-01 | 4.43E-01 | 4.57E-01 |
| Tcm | LG | FemaleA - MaleA | 9.20E-01 | 8.91E-01 | 6.38E-01 | 6.34E-01 |
| Tcm | LG | FemaleA - MaleX | 5.96E-01 | 5.64E-01 | 4.43E-01 | 4.18E-01 |
| Tcm | LG | FemaleX - FemaleA | 9.71E-01 | 9.77E-01 | 8.01E-01 | 7.97E-01 |
| Tcm | LG | FemaleX - MaleA | 9.45E-01 | 9.50E-01 | 6.65E-01 | 6.41E-01 |
| Tcm | LG | FemaleX - MaleX | 7.01E-01 | 6.92E-01 | 6.91E-01 | 6.77E-01 |
| Tcm | LG | MaleX - MaleA | 6.50E-01 | 6.25E-01 | 3.40E-01 | 3.03E-01 |
| Tcm | RT | FemaleA - MaleA | **1.87E-13** | **5.91E-13** | **4.13E-39** | **2.93E-38** |
| Tcm | RT | FemaleA - MaleX | **3.91E-25** | **3.88E-25** | **8.20E-27** | **1.08E-27** |
| Tcm | RT | FemaleX - FemaleA | **5.31E-03** | **9.11E-03** | 1.99E-01 | 3.24E-01 |
| Tcm | RT | FemaleX - MaleA | 8.78E-01 | 8.51E-01 | **2.79E-04** | **2.43E-04** |
| Tcm | RT | FemaleX - MaleX | **7.56E-28** | **4.44E-27** | **1.05E-22** | **4.64E-22** |
| Tcm | RT | MaleX - MaleA | **5.08E-42** | **7.78E-41** | **1.28E-58** | **3.27E-58** |
| Tpa | HD | FemaleA - MaleA | 6.75E-01 | 6.81E-01 | **2.05E-04** | **1.33E-04** |
| Tpa | HD | FemaleA - MaleX | 5.12E-02 | 8.83E-02 | **2.79E-04** | **2.20E-03** |
| Tpa | HD | FemaleX - FemaleA | 4.25E-01 | 6.04E-01 | 3.42E-01 | 5.08E-01 |
| Tpa | HD | FemaleX - MaleA | 5.96E-01 | 7.47E-01 | 6.25E-01 | 6.41E-01 |
| Tpa | HD | FemaleX - MaleX | 4.83E-01 | 4.73E-01 | 5.90E-02 | 1.11E-01 |
| Tpa | HD | MaleX - MaleA | 9.78E-02 | 1.52E-01 | 5.79E-02 | 1.01E-01 |
| Tpa | LG | FemaleA - MaleA | 9.45E-01 | 9.53E-01 | 6.77E-01 | 6.77E-01 |
| Tpa | LG | FemaleA - MaleX | 5.96E-01 | 8.34E-01 | 3.18E-01 | 5.13E-01 |
| Tpa | LG | FemaleX - FemaleA | 8.51E-01 | 8.80E-01 | 6.80E-01 | 9.26E-01 |
| Tpa | LG | FemaleX - MaleA | 8.78E-01 | 8.80E-01 | 8.01E-01 | 8.11E-01 |
| Tpa | LG | FemaleX - MaleX | 8.78E-01 | 7.82E-01 | 6.54E-01 | 5.91E-01 |
| Tpa | LG | MaleX - MaleA | 6.15E-01 | 8.49E-01 | 4.16E-01 | 6.00E-01 |
| Tpa | RT | FemaleA - MaleA | **5.63E-22** | **2.09E-15** | **4.80E-160** | **5.95E-144** |
| Tpa | RT | FemaleA - MaleX | **1.40E-93** | **2.24E-58** | **1.88E-42** | **6.93E-28** |
| Tpa | RT | FemaleX - FemaleA | 9.20E-01 | 4.07E-01 | 6.22E-01 | 3.77E-01 |
| Tpa | RT | FemaleX - MaleA | **9.92E-04** | 3.17E-01 | **9.64E-23** | **1.52E-10** |
| Tpa | RT | FemaleX - MaleX | **3.05E-58** | **1.04E-39** | **3.13E-29** | **1.28E-20** |
| Tpa | RT | MaleX - MaleA | **6.31E-140** | **1.27E-81** | **1.11E-134** | **8.02E-80** |
| Tps | HD | FemaleA - MaleA | 9.44E-01 | 8.91E-01 | 2.57E-01 | 2.76E-01 |
| Tps | HD | FemaleA - MaleX | 8.66E-02 | 2.85E-01 | 5.90E-02 | 2.76E-01 |
| Tps | HD | FemaleX - FemaleA | 1.89E-01 | 4.04E-01 | **4.44E-02** | 1.76E-01 |
| Tps | HD | FemaleX - MaleA | 2.17E-01 | 4.49E-01 | **1.15E-02** | 6.92E-02 |

| | | | | | | |
|---|---|---|---|---|---|---|
| Tps | HD | FemaleX - MaleX | 8.51E-01 | 8.98E-01 | 9.23E-01 | 8.58E-01 |
| Tps | HD | MaleX - MaleA | 9.78E-02 | 3.14E-01 | **1.69E-02** | 1.14E-01 |
| Tps | LG | FemaleA - MaleA | 1.77E-01 | 1.75E-01 | **1.03E-04** | **8.69E-05** |
| Tps | LG | FemaleA - MaleX | **2.64E-02** | 8.13E-02 | **4.73E-05** | **7.75E-04** |
| Tps | LG | FemaleX - FemaleA | 1.88E-01 | 2.85E-01 | 5.06E-02 | 1.42E-01 |
| Tps | LG | FemaleX - MaleA | 5.46E-01 | 6.86E-01 | 7.46E-01 | 9.32E-01 |
| Tps | LG | FemaleX - MaleX | 6.15E-01 | 7.15E-01 | 1.43E-01 | 2.04E-01 |
| Tps | LG | MaleX - MaleA | 1.37E-01 | 2.85E-01 | **2.42E-02** | 1.05E-01 |
| Tps | RT | FemaleA - MaleA | **3.98E-18** | **1.99E-17** | **3.32E-70** | **1.30E-68** |
| Tps | RT | FemaleA - MaleX | **1.84E-37** | **2.36E-36** | **6.55E-29** | **5.58E-27** |
| Tps | RT | FemaleX - FemaleA | 8.66E-02 | 6.38E-02 | 4.42E-01 | 2.50E-01 |
| Tps | RT | FemaleX - MaleA | 2.64E-01 | 4.63E-01 | **3.04E-10** | **4.40E-08** |
| Tps | RT | FemaleX - MaleX | **2.02E-33** | **1.68E-33** | **5.63E-23** | **2.35E-23** |
| Tps | RT | MaleX - MaleA | **2.98E-61** | **3.60E-58** | **2.84E-75** | **5.86E-69** |

**Table S8 | Sample information and accession numbers for the males sequenced in this study.** Reads are deposited under the bioproject accession PRJNA725673. Species names (sp) are abbreviated as follows: *Timema bartmani* = Tbi, *Timema cristinae* = Tce, *Timema californicum* = Tcm, *Timema podura* = Tpa, *Timema poppensis* = Tps

| Library Name | Sp | Population name | Population coord | Colour morph | Biosample accession | SRA run accession |
|---|---|---|---|---|---|---|
| Tbi_M_13_Tbi | Tbi | Jenks | 34.1700 N 117.0020 W | green | SAMN18898464 | SRR14340024, SRR14340035, SRR14340046, SRR14340057, SRR14340067, SRR14340078, SRR14340089, SRR14340162, SRR14340239, SRR14340250, SRR14340261, SRR14340272-SRR14340274 |
| Tbi_M_14_Tbi | Tbi | Jenks | 34.1700 N 117.0020 W | green | SAMN18898465 | SRR14340106, SRR14340117, SRR14340128, SRR14340139, SRR14340150, SRR14340161, SRR14340173, SRR14340184, SRR14340195, SRR14340206, SRR14340217, SRR14340228 |
| Tbi_M_15_Tbi | Tbi | Jenks | 34.1700 N 117.0020 W | grey | SAMN18898466 | SRR14340081-SRR14340088, SRR14340090-SRR14340092, SRR14340095 |
| Tbi_M_16_Tbi | Tbi | Jenks | 34.1700 N 117.0020 W | grey | SAMN18898467 | SRR14340068-SRR14340077, SRR14340079, SRR14340080 |
| Tce_M_05_HM15 | Tce | Ojai A007 | 34.5363 N 119.2444 W | brown | SAMN18898468 | SRR14340014, SRR14340053-SRR14340056, SRR14340058-SRR14340066 |
| Tce_M_06_HM16 | Tce | Ojai A007 | 34.5363 N 119.2444 W | green | SAMN18898469 | SRR14340038-SRR14340045, SRR14340047-SRR14340052 |
| Tce_M_07_HM33 | Tce | Ojai A007 | 34.5363 N 119.2444 W | green | SAMN18898470 | SRR14340022, SRR14340023, SRR14340025-SRR14340034, SRR14340036, SRR14340037 |
| Tce_M_08_HM61 | Tce | Ojai A007 | 34.5363 N 119.2444 W | green | SAMN18898471 | SRR14340013, SRR14340015-SRR14340021, SRR14340266-SRR14340271 |
| Tcm_M_01_HM148 | Tcm | Hamilton | 37.3432 N 121.6365 W | green | SAMN18898472 | SRR14340254, SRR14340257-SRR14340260, SRR14340262-SRR14340265 |
| Tcm_M_02_HM149 | Tcm | Hamilton | 37.3432 N 121.6365 W | green | SAMN18898473 | SRR14340241-SRR14340249, SRR14340251-SRR14340253, SRR14340255, SRR14340256 |
| Tcm_M_03_HM150 | Tcm | Hamilton | 37.3432 N 121.6365 W | green | SAMN18898474 | SRR14340225-SRR14340227, SRR14340229-SRR14340238, SRR14340240 |
| Tcm_M_04_HM151 | Tcm | Hamilton | 37.3432 N 121.6365 W | green | SAMN18898475 | SRR14340210-SRR14340216, SRR14340218-SRR14340224 |
| Tpa_M_09_Tpa | Tpa | Vista | 33.7976 N 116.7769 W | brown | SAMN18898476 | SRR14340194, SRR14340196-SRR14340205, SRR14340207- |

| | | | | | | |
|---|---|---|---|---|---|---|
| | | | | | | SRR14340209 |
| Tpa_M_10 _Tpa | Tpa | Vista | 33.7976 N 116.7769 W | brown | SAMN18898477 | SRR14340179-SRR14340183, SRR14340185-SRR14340193 |
| Tpa_M_11 _Tpa | Tpa | Vista | 33.7976 N 116.7769 W | brown | SAMN18898478 | SRR14340164-SRR14340172, SRR14340174-SRR14340178 |
| Tpa_M_12 _Tpa | Tpa | Vista | 33.7976 N 116.7769 W | brown | SAMN18898479 | SRR14340147-SRR14340149, SRR14340151-SRR14340160, SRR14340163 |
| Tps_M_17 _HM99 | Tps | Bear Creek | 37.1655 N 122.0156 W | green | SAMN18898480 | SRR14340133-SRR14340138, SRR14340140-SRR14340146 |
| Tps_M_18 _HM100 | Tps | Bear Creek | 37.1655 N 122.0156 W | green | SAMN18898481 | SRR14340120-SRR14340127, SRR14340129-SRR14340132 |
| Tps_M_19 _HM101 | Tps | Bear Creek | 37.1655 N 122.0156 W | green | SAMN18898482 | SRR14340107-SRR14340116, SRR14340118, SRR14340119 |
| Tps_M_20 _15255 | Tps | Summit Road | 37.2223 N 122.0878 W | green | SAMN18898483 | SRR14340093, SRR14340094, SRR14340096-SRR14340105 |

**Table S9 | Number of genes with evidence for positive selection when comparing site-models M8 and M8a on the X and autosomes.** The proportion of genes with positively selected sites on the X and autosomes was not significantly different (Fisher's exact test p-value = 0.1077).

| Chromosome type | N genes with positively selected sites | Total genes |
|---|---|---|
| Autosomes | 15 | 4269 |
| X | 3 | 297 |
